# Impact of *Fkbp5* × Early Life Adversity × Sex in Humanized Mice on Multidimensional Stress Responses and Circadian Rhythmicity

**DOI:** 10.1101/2021.07.06.450863

**Authors:** Verena Nold, Michelle Portenhauser, Dolores Del Prete, Andrea Blasius, Isabella Harris, Eliza Koros, Tatiana Peleh, Bastian Hengerer, Iris-Tatjana Kolassa, Michal Slezak, Kelly Ann Allers

**Author notes:** **Corresponding Author**, Boehringer Ingelheim Pharma GmbH & Co KG, CNSDR (and Ulm University, Clinical & Biological Psychology), Birkendorferstraße 65, 88397 Biberach an der Riß, Germany.

## Abstract

The cumulative load of genetic predisposition, early life adversity (ELA) and lifestyle shapes the prevalence of psychiatric disorders. Single nucleotide polymorphisms (SNPs) in the human *FKBP5* gene were shown to modulate disease risk. To enable investigation of disease-related SNPs in behaviorally relevant context, we generated humanized mouse lines carrying either the risk (AT) or the resiliency (CG) allele of the rs1360780 locus and exposed litters of these mice to maternal separation. Behavioral and physiological aspects of their adult stress responsiveness displayed interactions of genotype, early life condition and sex. In humanized females carrying the CG-but not the AT-allele, ELA led to altered HPA-axis functioning, exploratory behavior and sociability. These changes correlated with differential expression of genes in the hypothalamus, where synaptic transmission, metabolism, and circadian entrainment pathways were deregulated. Our data suggest an integrative role of *FKBP5* in shaping the sex-specific outcome of ELA in adulthood.

## 1 Introduction

Stress responses are essential to adjust physiology and behaviour to recurrently changing environmental demands [1], but corrupted stress responses are a hallmark feature of psychiatric conditions [2]. The susceptibility or resilience to develop psychiatric disorders can be attributed to interactions of genetic predispositions and environmental factors [3]. Among environmental factors, early life adversity (ELA) is found to be especially detrimental given that aberrations during development will influence the affected individuals throughout life [4]. Childhood maltreatment is common in the history of many psychiatric patients and comprises experiences of physical, sexual and emotional abuse, as well as physical and emotional neglect [5]. Such experiences during development shape disease prevalence in later life through alterations in HPA-axis programming, stress coping strategies and brain connectivity [6].

With respect to genetic predispositions, the regulation of glucocorticoid signaling is a prominent research target since glucocorticoids are a key messenger for the spread and initiation of stress responsive signaling. This regulation is fine-tuned in a timing- and dose-dependent manner and depends on the individual cellular set-up such as the relative expression of glucocorticoid receptors and its regulators [7]. Expression levels of *FKBP5*, a potent negative regulator of glucocorticoid signaling, is part of this cellular identity and is itself a target of glucocorticoid-mediated gene transcription [8]. Single nucleotide polymorphisms (SNPs) inside the human *FKBP5* gene are associated with differential induction of the FKBP51 protein upon glucocorticoid stimulation [9] and add to the variability of stress perception and response in the population [10]. Carriers of the high induction allele rs1360780-A/T of *FKBP5* who suffered from ELA are more prone to develop psychiatric symptoms in later life than individuals without such preconditioning [11]. Importantly, sex-dependent differences in the interaction of *FKBP5* and life adversities have been associated to a higher prevalence of depression in females [12]. Despite the strong negative impact of psychiatric disorders on quality of life and productivity, the underlying processes linking *FKBP5* genotypes, stress regulation and pathological transitions are not fully understood. Animal models offer a possibility to investigate gene × environment interactions in a timely resolved manner. In depth analyses of laboratory mouse sequences in-house indicated numerous *Fkbp5* SNPs that vary by strain. However, no SNPs at the same location or with the same functional impact as found in humans occur naturally in rodents.

This lack of an animal model suited to exploring human *FKBP5* SNPs hinders elucidation of causal relationships and mechanisms underlying disease development and progression. Therefore, we previously generated *Fkbp5*-humanized mice carrying either the risk-associated high induction AT-allele of rs1360780 or the resiliency-associated CG-allele. Initial characterization of primary CNS-cell types derived from these mice revealed that the presence of the AT-allele results in the increased expression of *Fkbp5* upon stimulation of the glucocorticoid receptor compared to the CG-allele [7]. This initial characterization prompted us to exploit this new model to examine the *Fkbp5* × ELA interactions on the stress response system in adulthood. We exposed AT and CG-allele carrying mice to prolonged maternal separation stress, since this paradigm is broadly used to mimic ELA in rodents [13]. When mice reached adulthood, the performance of the HPA-axis and behavioral response of *Fkbp5*-humanized mice to mild stressors were measured. Furthermore, we investigated the transcriptomic profiles in several brain regions engaged in stress processing. Lastly, astrocytes and neurons derived from human induced pluripotent stem cells (hiPSCs) were analyzed for SNP-based differens in their expression profiles.

The goals of the study were to validate the *Fkbp5* × ELA model by 1) determining wether ELA would cause alterations in the offspring’s adult behaviour and physiology compared to controls, 2) determining whether risk AT-allele carriers would respond differently to ELA than CG-allele carriers, 3) assessing which pathways are involved in the adaptation to ELA in context of risk and resilience associated SNPs. A more far-reaching aim was to demonstrate that the humanized *Fkpb5* × ELA mouse model can be used to further investigate the influence of the human *FKBP5* gene variants on the risk and resilience to stress and to further elucidate their contribution to psychiatric disorders.

## 2 Results

Prolonged separation from mothers and peers was performed for the first three weeks of postnatal life to model ELA. In parallel, control mice were housed with littermates and received undisturbed maternal care until weaning. An overview of the group sizes of the cohort is provided in Tab. 1. On postnatal day 21, pups were weaned and grown to adulthood with physiological and behavioral examination starting at 10 weeks of age (Fig. 1). Exploration of novel environments offers an easily accessible measure of mild stress in rodents [14]. Therefore, we challenged control and ELA-exposed mice with novel situations to probe for their stress coping strategies. The same procedures were simultaneously carried out in wild type mice of both sexes. The data on their HPA-axis functioning (Fig. S1) and behavior (Fig. S2) are visualized in the supplements for reference. Statistical analyses were performed jointly for males and females to address differences between sex, ELA exposure, *Fkbp5*-genotypes and the interaction thereof. Details of the descriptive analyses, model summaries and analysis of variance (ANOVA) results are provided in the supplements 7. A significant effect of sex × genotype × treatment interaction and significant two-way interactions in the vast majority of measured parameters were detected and are detailed in the following paragraphs.

**Table 1:**
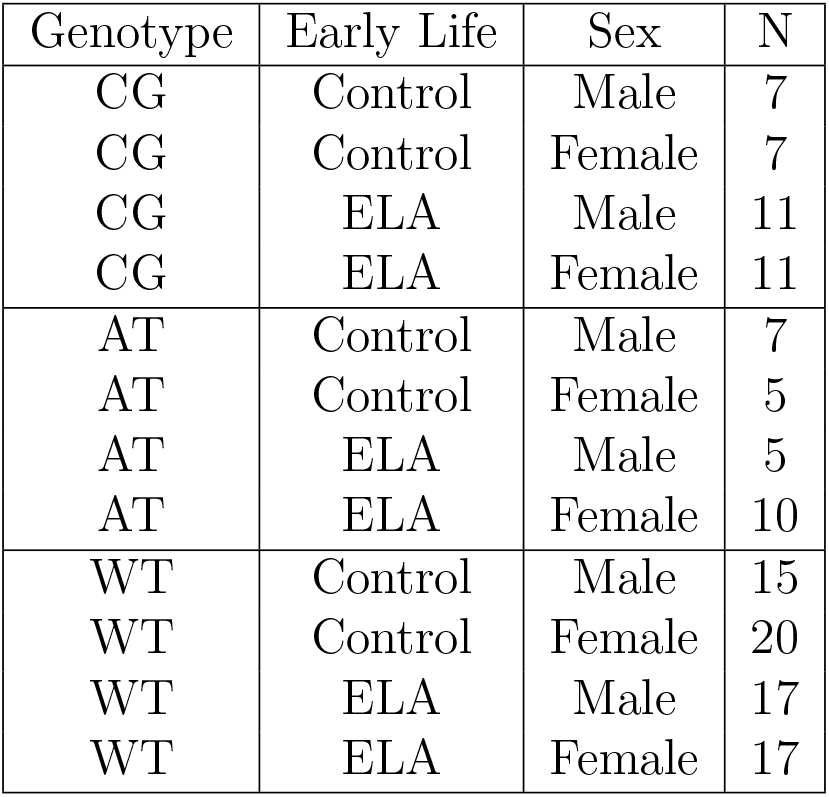
Overview of the Study Cohort

**Figure 1:**
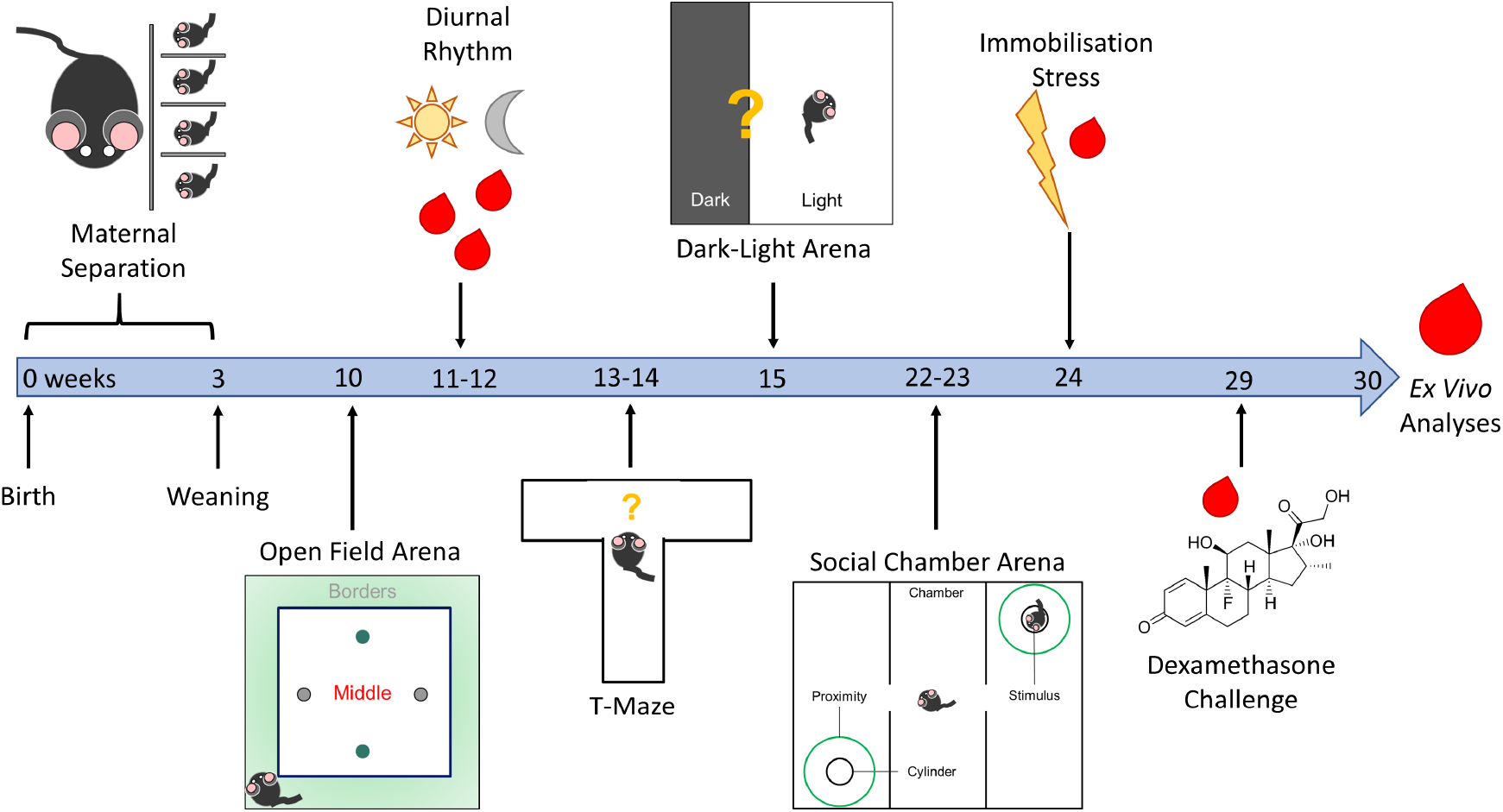
Time Line of Experiments. Study overview of *in vivo* and *ex vivo* experiments during the life time of *Fkbp5*-humanized mice with ELA. The same time line, except for the maternal separation, was applied to control mice in parallel.

### 2.1 Early Life Adversity and *Fkbp5*-Variants Shift and Attenuate Diurnal HPA-Axis Rhythmicity

To measure the impact of *Fkbp5* SNPs in combination with ELA on the diurnal performance of the HPA-axis, the plasma corticosterone concentration of samples collected at three time points was assessed. As confirmed in the wild type mice (Fig. S1), these timepoints were reflecting the diurnal nadir (morning), peak (evening) and one intermediate state (noon). In control females carrying the CG-allele, the expected increase of plasma corticosterone over the course of the day was observed, with a clear peak towards the evening (Fig. 2a, Tab. S1, Tab. S2, Tab. S3). Following ELA exposure, the highest concentration was instead measured at noon. The increase of plasma corticosterone levels in AT-allele carrying control females was not statistically significant, regardless of ELA exposure.

**Figure 2:**
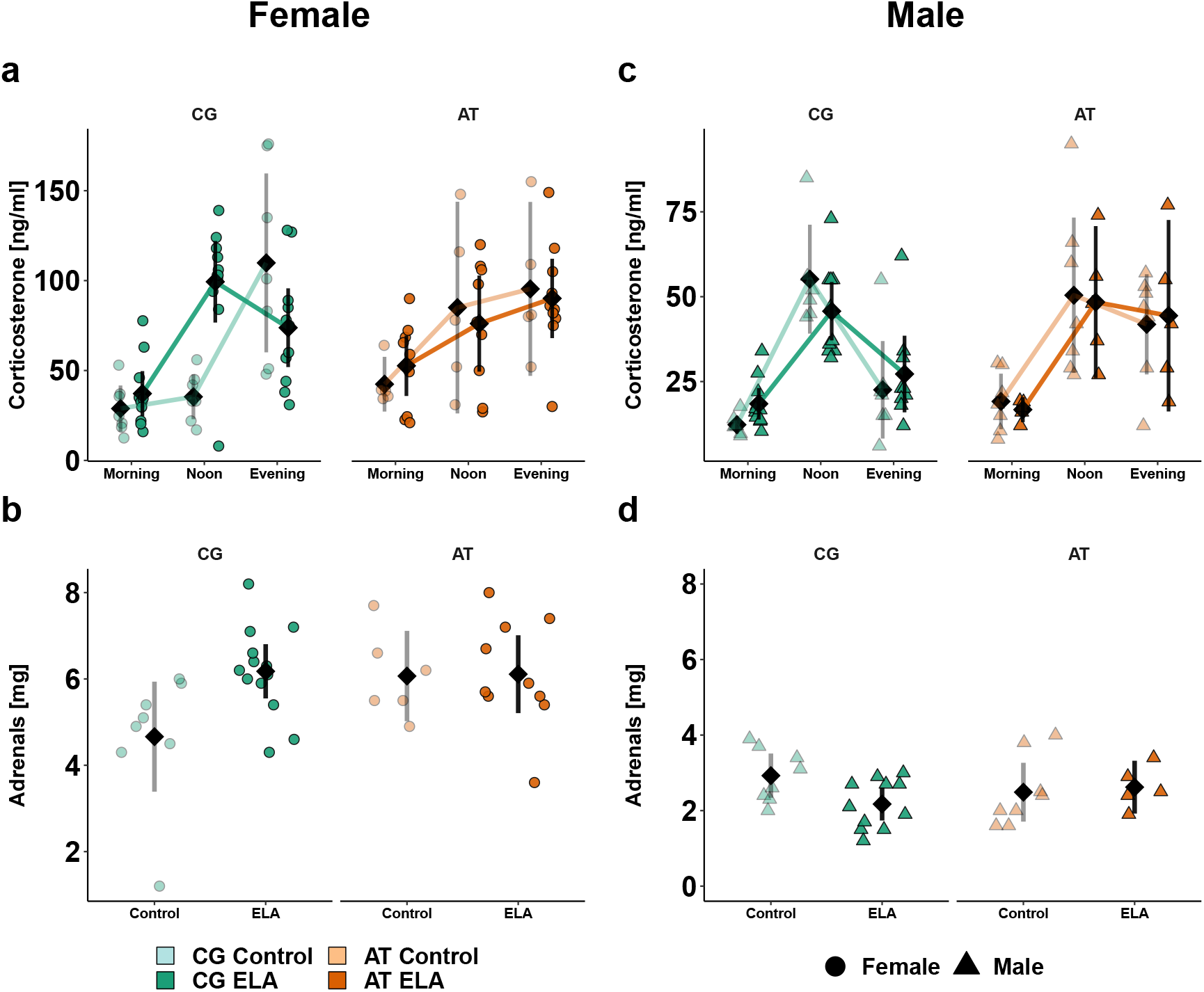
*Fkbp5*-Genotype × ELA Influence the Unstimulated HPA-Axis Activity in a Sex-Dependent Manner. Individual animal data is shown alongside with the mean (black diamond) ± 95% confidence interval. Diurnal rhythmicity of corticosterone plasma levels in female (a) and male (c) *Fkbp5*-humanized control or ELA-exposed mice. A different scale for males than females was used to make the pattern better visible. Comparison of adrenal weights in females (b) and males (d). Descriptive statistics, model summaries and ANOVA results are provided in Tab. S1, Tab. S2 and Tab. S3 for diurnal corticosterone and in Tab. S4, Tab. S5 and Tab. S6 for adrenal weight.

In line with these findings, the adrenal weight was increased by ELA in CG-allele carrying females, while in the AT-allele carrying females, the adrenal weight tended to be already increased in control mice compared to CG controls without further increase upon ELA (Fig. 2b, Tab. S4, Tab. S5, Tab. S6).

In *Fkbp5*-humanized males, the diurnal plasma corticosterone concentration peaked towards noon with CG-vs-AT-allele carriers showing a decrease towards the evening, regardless of ELA exposure (Fig. 2c). The detected diurnal amplitude of corticosterone was smaller in males than females. No significant differences in the adrenal weights were observed among males (Fig. 2d), but male vs. female adrenal weights were significantly lower.

Taken together, female AT-vs. CG-allele carriers are genetically predisposed to less pronounced diurnal HPA-axis rhythmicity resulting in elevated corticosterone levels at time points were mice usually would rest. Lower diurnal corticosterone amplitudes and adrenal weights in males vs. females suggest a different corticosterone secretion capacity between sexes.

### 2.2 Early Life Adversity Increases Responsiveness to Novel Environments Dependent on *Fkbp5*-Genotype and Sex

Exposure to novel environments as mild stress was applied to determine natural behavior and coping strategies. First, behavior in open field test arenas was assessed to obtain a measure of locomotor activity at the beginning of and throughout the murine active phase (18:30 – 05:30). Overall activity within the first 15 minutes, including running and rearing, was assessed by measuring the frequency of crossing light beams (Fig. 3a, Tab. S7, Tab. S8, Tab. S9). During this period, the activity decreased over time with early life condition and sex showing an interaction with time. As in wild type females (Fig. S2a), the group of CG control females displayed habituation in the shape of a strong decrease in activity, while the exposure to ELA led to flattening of the 15 minutes activity profile and thus slower habituation (Fig. 3a). Habituation in AT-allele carrying females tended to be slower than in CG-controls, regardless of early life condition.

**Figure 3:**
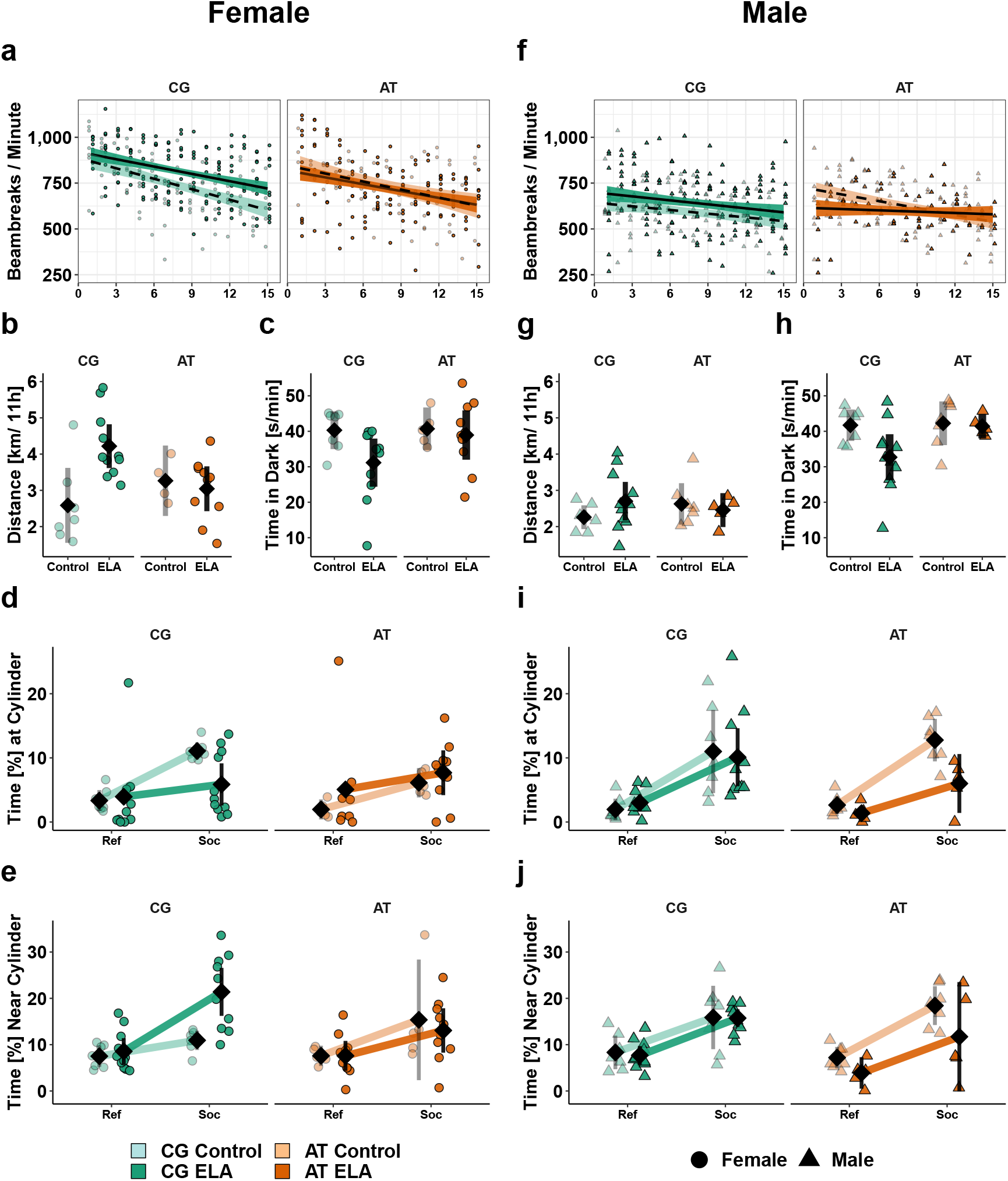
Sex × *Fkbp5*-Genotype × ELA Interactions Alter Activity in Humanized Mice with Impact on Behavioral Responses to Mild Stress. Individual data is shown alongside with the mean ± 95% confidence intervals. Exploration activity (light beams crossing / minute) during the first 15 minutes in a novel environment in females (**a**) and males (**f**). Total distance [km] females (**b**) and males (**g**) moved during the night. Average time [s/min] females (**c**) and males (**h**) spent in the dark compartment. Time [%] females (**d**) and males (**i**) spent at the cylinder with (Soc) or without (Ref) an unfamiliar mouse. Time [%] females (**e**) and males (**j**) spent in the area surrounding the cylinder with (Soc) or without (Ref) an unfamiliar. Descriptive statistics, model summary and ANOVA for the habituation activity are provided in Tab. S7, Tab. S8 and Tab. S9, for the nocturnal distance in Tab. S10, Tab. S11 and Tab. S12, for the Dark-Light-Test in Tab. S19, Tab. S20 and Tab. S21, for social interaction time in Tab. S22, Tab. S23 and Tab. S24, while for the time in social distance in Tab. S25, Tab. S26 and Tab. S27.

Analyses of the total nocturnal distance revealed main effects of sex and early life condition, as well as an interaction effect of ELA × genotype (Fig. 3b, Tab. S10, Tab. S11, Tab. S12). While ELA-exposed female CG-allele carriers were more active than controls, AT-allele carrying females with ELA experience were indistinguishable from controls.

In the spontaneous alternations T-maze, ELA did not affect the fraction of alternations between left or right side of the maze, irrespective of genotype or sex (Fig. S3a, Tab. S14, Tab.S13, Tab. S15), suggesting no impact on working memory performance. However, ELA-exposed mice performed the task significantly faster than the respective control group and females were quicker than males (Fig. S3b, Tab.S16, Tab. S17, Tab. S18).

In the dark-light test, ELA decreased the mean time spent in the dark compartment. Moreover, a trend for *Fkbp5*-genotype related effects was seen, with CG-allele carrying females compared to controls showing the ELA effect, while in the AT-allele carriers the control group was indistinguishable from ELA-exposed females (Fig. 3c, Tab. S19, Tab. S20, Tab. S21).

Finally, we measured social preference in the 3-social-chamber test. Pairwise comparisons of compartment effects separated by early life conditions, genotype and sex revealed significant differences: CG-allele control females showed social preference, measured by the time the mouse spent in the nearest vicinity of the cylinder with the social stimulus (Fig. 3d, Tab. S22, Tab. S23, Tab. S24). The exposure to ELA led to decrease of this parameter, while simultaneously we observed a significant increase in the time spent in the chamber, but in 5 cm distance from the occupied cylinder (Fig. 3e, Tab. S25, Tab. S26, Tab. S27). In contrast, AT-allele carrying control females spent less time interacting with the unfamiliar mouse, as compared to CG-allele carrying controls. ELA did not further change this parameter, and the time of direct interaction vs. time in ‘social distance’ was similar in the AT-allele carrying controls and ELA-exposed females.

In males, the activity measured in the open field arena (Fig. 3f and g) and working memory assessed as spontaneous alternations in the T-maze (Fig. S3a) were similar among groups. Like in females, male CG-allele carriers with ELA tended to spend less time in the dark compartment (Fig. 3h), and to complete the T-maze test faster (Fig. S3b) than controls. In contrast to females, the social preference was not affected by ELA in male CG-allele carriers, but decreased in ELA-exposed AT-allele carrying males (Fig. 3i, Fig. 3j).

Overall, the data on behavioral responses to mild stress elicited by novel environments suggest that the effects of ELA on these read outs depend on the genetic variants of *Fkbp5 ×* sex.

### 2.3 HPA-Axis Responses are Stronger in Females than Males

To probe the HPA-axis reactivity to acute induction and negative feedback, we measured plasma corticosterone after 5 minutes of restraint stress and 6 h after a single intraperitoneal injection of the synthetic glucocorticoid dexamethasone. In all mice, corticosterone increased in response to restraint stress without a differential effect of genotype. However, the slope was steeper in females (Fig. 4a) compared to males (Fig. 4b) among ELA-exposed mice. Similarly, all mice responded to dexamethasone with reduced corticosterone levels, suggesting a suppression of the endogenous corticosterone secretion. *Post hoc* analyses revealed that the slope of decrease was overall steeper in females exposed to ELA relative to controls (Fig. 4c). In males, no effect of ELA or *Fkbp5*-genotype on the HPA-axis responsiveness to negative feedback was statistically significant (Fig. 4d).

**Figure 4:**
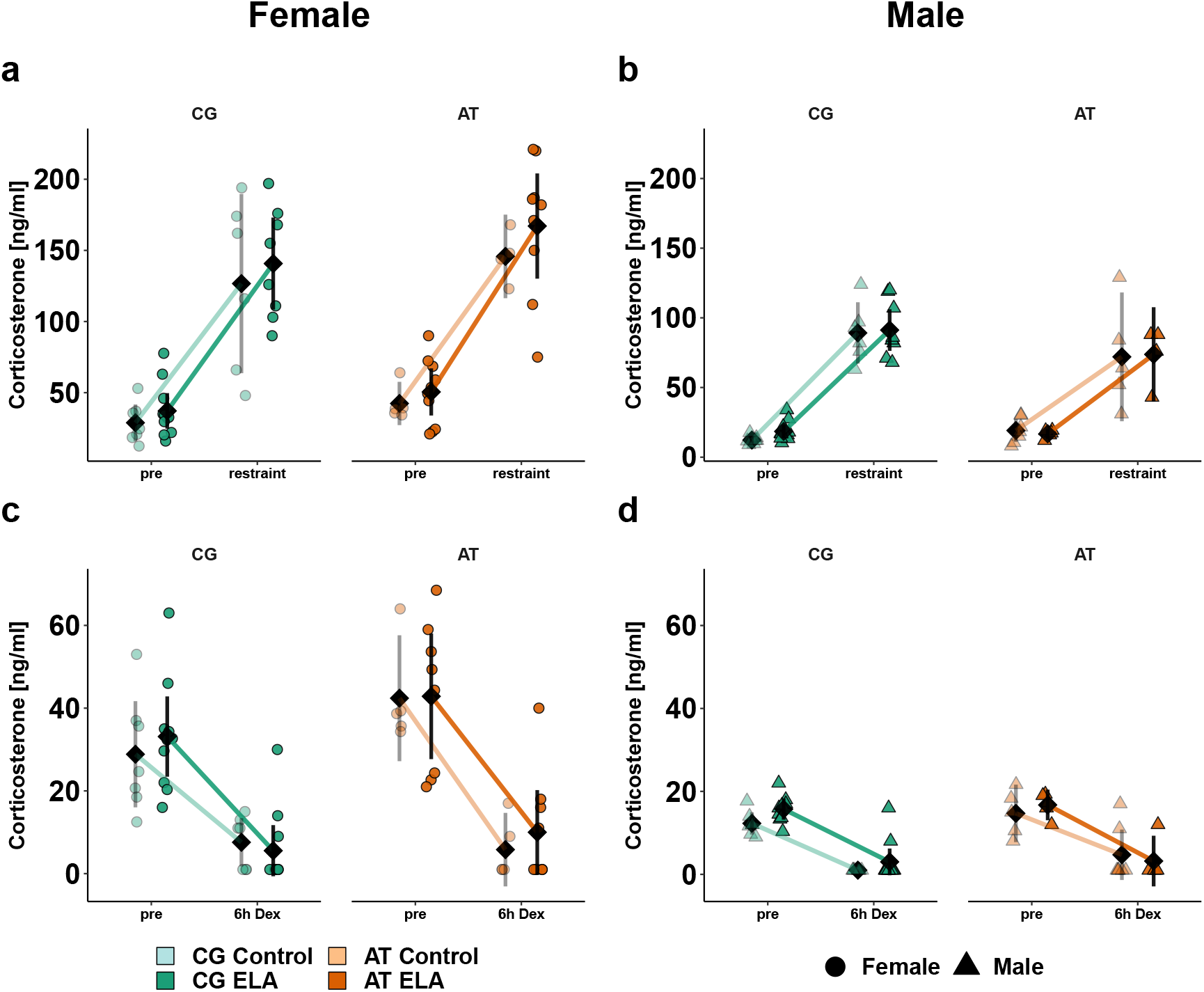
Stimulated HPA-Axis Reactivity in *Fkbp5*-Humanized Females is Greater than in Males. Individual animal data is shown alongside with the mean ± 95% confidence interval (black). Acute responsiveness of the HPA-axis assessed by comparison of plasma corticosterone levels before and 5 minutes after restraint stress in females (**a**) and males (**b**). Suppression of endogenous corticosterone production 6 hours after dexamethasone injection in females (**c**) and males (**d**). Descriptive statistics (Tab. S1) as well as model summaries and ANOVA results for restraint stress corticosterone are available in Tab. S28 and Tab. S29, and in Tab. S30 and Tab. S31 for dexamethasone suppression.

In summary, the responsiveness of the HPA-axis is preserved in *Fkbp5*-humanized mice.

### 2.4 Transcription in Stress-Responsive Brain Regions is Affected by *Fkbp5* × ELA

To identify transcriptional differences that could be related to the observed differences in behavior and HPA-axis physiology of *Fkbp5*-humanized mice × ELA, mRNA sequencing and analyses of differential gene expression were carried out. Given the sexual dimorphism in the *in vivo* experiments, analyses were limited to females and focused on hypothalamus, ventral and dorsal hippocampus as brain regions engaged in stress regulation [15].

In the SNP-comparison among controls, more differentially expressed genes (DEGs) were found in the hypothalamus (579), followed by ventral (41) and dorsal (2) hippocampus (Tab. 2). Among ELA-exposed individuals, substantially more DEGs between the SNP variants were detected than in controls, underscoring the interaction of ELA × *Fkbp5*-genotype. Looking at the effect of ELA, fewer differences were detected in AT-allele carriers (114) than in CG-allele (903) carriers. This matches to the behavior and HPA-axis data, where the AT-allele alone was found to predispose to a ‘stress-like’ phenotype, with few additional impact of ELA.

**Table 2:** Counts of Differentially Expressed Genes in Subgroups of *Fkbp5*-Humanized Female Mice.

| Comparison | Tissue | Direction | Control | ELA |
| --- | --- | --- | --- | --- |
| *Fkbp5*-Genotype | Hypothalamus | AT > CG | 349 | 561 |
|  | Hypothalamus | CG > AT | 230 | 855 |
|  | Ventral Hippocampus | AT > CG | 18 | 468 |
|  | Ventral Hippocampus | CG > AT | 23 | 457 |
|  | Dorsal Hippocampus | AT > CG | 1 | 798 |
|  | Dorsal Hippocampus | CG > AT | 1 | 844 |
| Comparison | Tissue | Direction | CG | AT |
| Early Life Condition | Hypothalamus | ELA > Con | 410 | 4 |
|  | Hypothalamus | Con > ELA | 195 | 29 |
|  | Ventral Hippocampus | ELA > Con | 16 | 0 |
|  | Ventral Hippocampus | Con > ELA | 10 | 0 |
|  | Dorsal Hippocampus | ELA > Con | 145 | 27 |
|  | Dorsal Hippocampus | Con > ELA | 127 | 54 |

Adopting knowledge from the SNP effects in humans, the overlap and uniqueness of the DEGs from the two comparisons and subgroups were analyzed for nomination of potential resiliency- or vulnerability-related genes. Genes linked to CNS-development such as *Mab21l2, Gart* and *Lipt2* were spotted as potentially vulnerability-related and were changed in opposite directions, with AT- vs. CG-allele carriers displaying a lower expression.

A second analysis focussing on gene clusters related to (developmental) neurological disorders using a two-step core and comparison analysis of the commercial software *Ingenuity* (Qiagen) confirmed that the ELA-responsive DEGs in both mouse lines have an impact on neurological and psychiatric symptoms (Fig. S4). In eight of the shown 30 deregulated clusters e.g. comprising ‘congenital neurological disorder’ or ‘learning’, the effects were opposite between AT- vs. CG-allele carriers.

In sum, the counts of DEGs and their accordant vs. discordant overlap between the analyzed subgroups suggest that the *Fkbp5* × ELA interaction on gene expression may have relevance for neurologic and psychiatric symptomatology.

### 2.5 The AT-Allele and ELA Reduce CNS Communication but Increase Metabolism

To identify how the DEGs might be linked to disorders via their role in cellular pathways, their over-representation in metabolism and signaling-related pathways listed in the Kyoto Encyclopedia of Genes and Genomes (KEGG) was assessed. The analyses revealed significantly altered pathways in the hypothalamus and ventral hippocampus (Tab. 3). The direction of change between *Fkbp5*-genotypes differed dependent on function, with pathways related to neuronal communication rather showing a downregulation, and pathways related to metabolism rather showing an upregulation in AT- vs. CG-allele carriers.

**Table 3:**
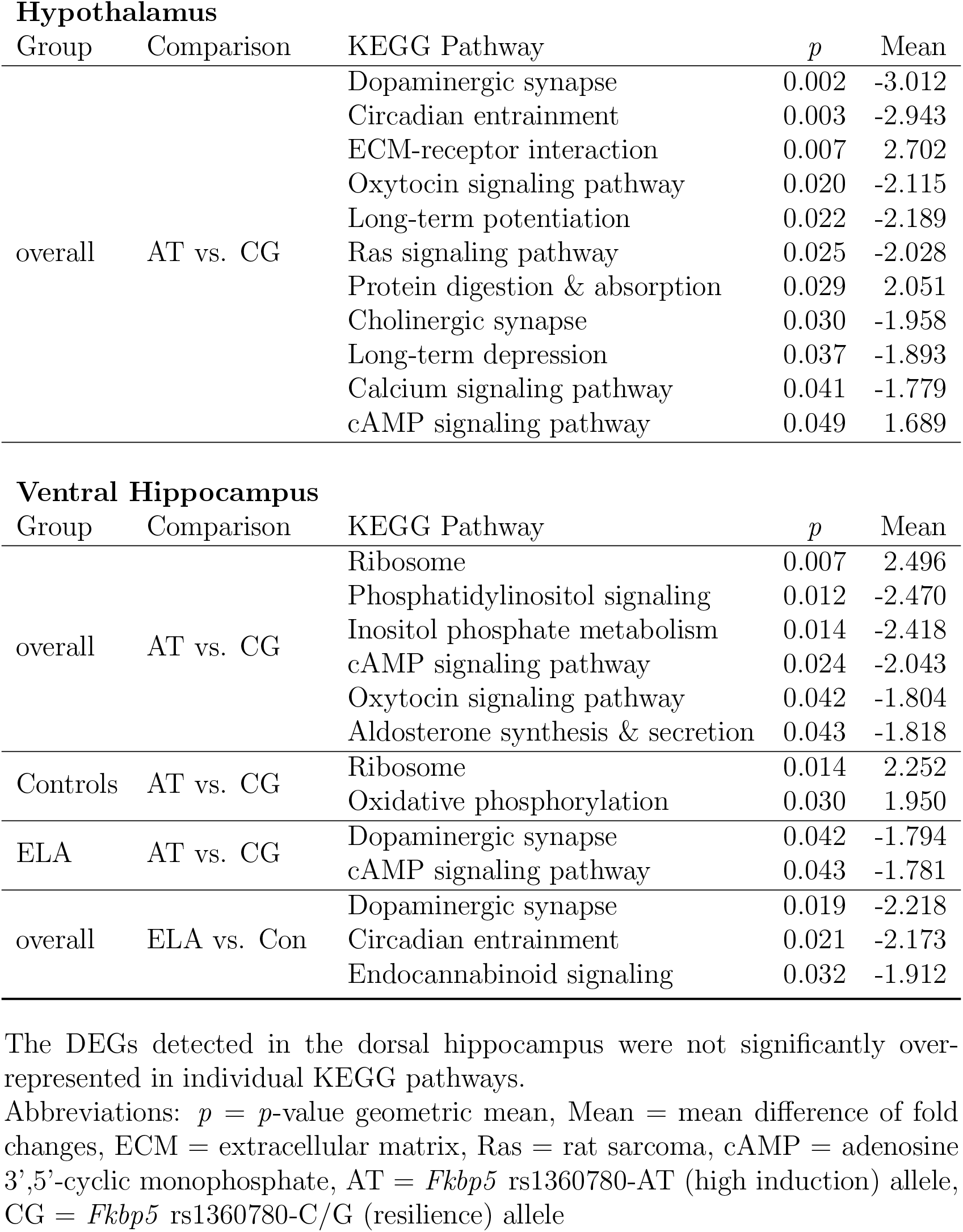
Enriched KEGG Pathways in *Fkbp5*-Humanized Females.

| Hypothalamus |  |  |  |  |
| --- | --- | --- | --- | --- |
| Group | Comparison | KEGG Pathway | *p* | Mean |
| overall | AT vs. CG | Dopaminergic synapse | 0.002 | -3.012 |
|  |  | Circadian entrainment | 0.003 | -2.943 |
|  |  | ECM-receptor interaction | 0.007 | 2.702 |
|  |  | Oxytocin signaling pathway | 0.020 | -2.115 |
|  |  | Long-term potentiation | 0.022 | -2.189 |
|  |  | Ras signaling pathway | 0.025 | -2.028 |
|  |  | Protein digestion & absorption | 0.029 | 2.051 |
|  |  | Cholinergic synapse | 0.030 | -1.958 |
|  |  | Long-term depression | 0.037 | -1.893 |
|  |  | Calcium signaling pathway | 0.041 | -1.779 |
|  |  | cAMP signaling pathway | 0.049 | 1.689 |
| Ventral Hippocampus |  |  |  |  |
| Group | Comparison | KEGG Pathway | *p* | Mean |
| overall | AT vs. CG | Ribosome | 0.007 | 2.496 |
|  |  | Phosphatidylinositol signaling | 0.012 | -2.470 |
|  |  | Inositol phosphate metabolism | 0.014 | -2.418 |
|  |  | cAMP signaling pathway | 0.024 | -2.043 |
|  |  | Oxytocin signaling pathway | 0.042 | -1.804 |
|  |  | Aldosterone synthesis & secretion | 0.043 | -1.818 |
| Controls | AT vs. CG | Ribosome | 0.014 | 2.252 |
|  |  | Oxidative phosphorylation | 0.030 | 1.950 |
| ELA | AT vs. CG | Dopaminergic synapse | 0.042 | -1.794 |
|  |  | cAMP signaling pathway | 0.043 | -1.781 |
| overall | ELA vs. Con | Dopaminergic synapse | 0.019 | -2.218 |
|  |  | Circadian entrainment | 0.021 | -2.173 |
|  |  | Endocannabinoid signaling | 0.032 | -1.912 |

The DEGs detected in the dorsal hippocampus were not significantly over-represented in individual KEGG pathways.
Abbreviations: *p* = *p*-value geometric mean, Mean = mean difference of fold changes, ECM = extracellular matrix, Ras = rat sarcoma, cAMP = adenosine 3',5'-cyclic monophosphate, AT = *Fkbp5* rs1360780-AT (high induction) allele, CG = *Fkbp5* rs1360780-C/G (resilience) allele

In the hypothalamus, the most significantly downregulated pathways included circadian entrainment, regulation of synaptic plasticity via long-term potentiation and depression as well as activity of dopaminergic and cholinergic synapses together with changes in calcium, cAMP and oxytocin signaling. In the ventral hippocampus, reduced expression of synaptic communication in AT- vs. CG-allele carriers was repeated. Especially in the ELA-subgroup, lower expression of genes related to cAMP signaling and dopaminergic synapses were found in AT-allele carriers compared to CG-allele carriers. Independent of strain, ELA was linked to lower expression of transcripts related to endocannabinoid and circadian entrainment relative to controls. For genes in pathways related to metabolism, such as protein absorption and digestion in the hypothalamus or ribosome activity and oxidative phosphorylation in the ventral hippocampus of controls, higher expression in AT-allele carriers relative to CG-allele carriers was observed.

The mRNA of neurons and astrocytes derived from human induced pluripotent stem cells (hiPSCs) of rs1360780 SNP carriers was sequenced and used to qualitatively validate the SNP-dependence of the observed differences in an independent expression system. In both cell types, comparable SNP-based expression differences, which might indicate less synaptic communication in AT- vs. CG-allele carriers, were seen. However, the distribution within the pathways differed between hiPSC- and mouse derived samples. For example, more differential expression in the upstream vs. downstream members of the circadian entrainment pathway was seen in the *Fkbp5*-humanized mice, while in the hiPSCs rather the expression of downstream targets was changed (Fig. S5). Moreover, the expression patterns in astrocytes vs. neurons were more similar to the patterns seen in mice.

The KEGG pathway analyses imply that ELA and the AT-allele both lead to less entrainment of diurnal HPA-axis rhythmicity. This lower entrainment of sleep-wake states may interact with the decreased ability of AT- vs. CG-allele carriers to process incoming inputs via synaptic communication.

### 2.6 Lower Glucocorticoid Sensitivity of the Hippocampus is Modulated by *Fkbp5* Genotype

To estimate how much impact the potentially altered glucocorticoid exposure due to differences in circadian entrainment and synaptic signaling might exert on the hypothalamus, ventral and dorsal hippocampus, the expression levels of genes related to glucocorticoid signaling were compared (Fig. 5). This analysis provides insights in the likelihood of the brain regions to respond to glucocorticoid stimulation. While expression levels of the glucocorticoid receptor (*Nr3c1*) and heat shock protein 90 (*Hsp90ab1*) were comparable between all three brain regions, the mineralocorticoid receptor (*Nr3c2*) was less expressed in the hypothalamus than in the hippocampus, with the ventral hippocampus displaying the highest expression. Moreover, *Fkbp5* was less expressed in the hypothalamus than hippocampus and the AT- vs. GC-allele was associated with a lower *Fkbp5* expression in dorsal and ventral hippocampus. **Considering the gene functions, the hypothalamus appears to be more sensitive to glucocorticoid receptor mediated signaling than the hippocampus, with CG- vs. AT-allele hippocampi being more protected.** The decreased cerebral expression of genes related to synaptic communication in AT- vs. CG-allele carriers might be a compensatory mechanism to prevent excessive excitation. To test whether the expression levels of the identified DEGs and *Fkbp5* could be linked to the observed behavioral and physiological differences, tissue-wise correlation analyses were carried out. For each brain region, the top 10 correlations are provided in Tab. S34 (the full list of correlations will be provided upon request). In all three brain regions, the majority of DEGs correlated with *Fkbp5*. In the hypothalamus, gap junction protein *β* 1 (*Gjb1*) showed a correlation with the time spent in the dark compartment of the test arena, while the membrane-associated, tyrosine-specific kinase 1 (*Pkmyt1*) and the nicotinic acetylcholine receptor subunit 7 (*Chrna7*, regression shown in S6) were linked to morning corticosterone levels. This could indicate an association between some hypothalamic DEGs and differences in HPA-axis functioning and behavior. **The correlation analyses suggest a linkage between expression levels of *Fkbp5* and DEGs in brain regions relevant for stress processing.**

**Figure 5:**
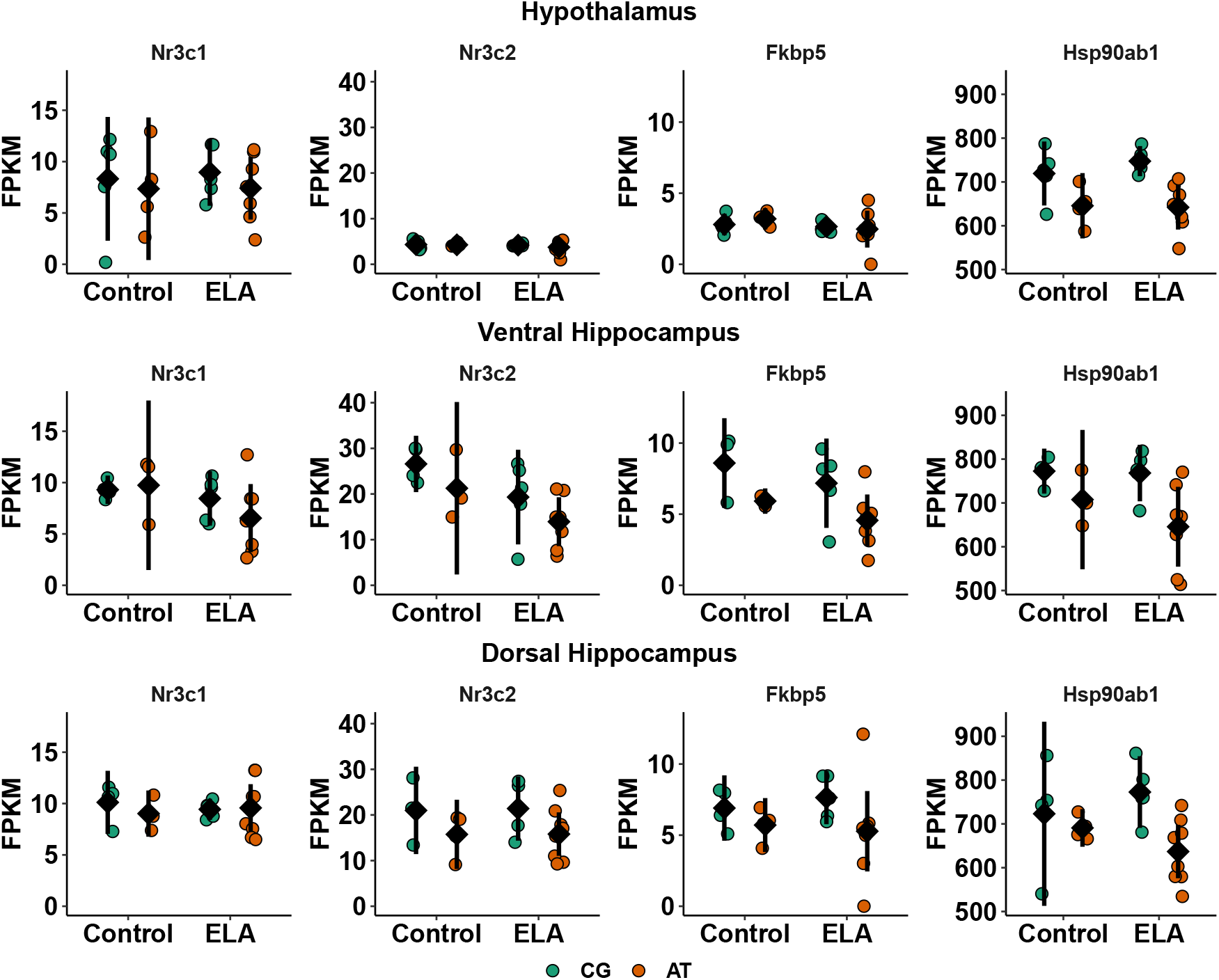
Brain Region Specific Expression Levels of Glucocorticoid Signalling Regulators. Expression levels [FPKM] of the glucocorticoid receptor (*Nr3c1*, GR), mineralocorticoid receptor (*Nr3c2*), *Fkbp5* and heat shock protein 90 (*Hsp90ab1*) of individual female AT- or CG-allele carriers that experienced ELA or undisturbed maternal care (control) are visualized alongside the mean ± 95% confidence interval. Plots are shown separate for hypothalamus (**top**), ventral hippocampus (**middle**) and dorsal hippocampus (**bottom**). Descriptive statistics and an overview of the significant model terms in the ANOVA are provided in Supplementary Tab. S32 and S33, respectively.

## 3 Discussion

The present study has demonstrated a gene × environment interaction in novel *Fkbp5*-humanized mice indicating that the model is suited to investigate the effects of ELA in the context of risk- and resiliency-related SNPs. Early life adversity elicited by maternal separation has differential impact on adult physiology and behavior based on genetic predisposition imparted by *Fkbp5* alleles. This is demonstrated by changes in locomotor, social, and anxious behavior. Additionally, diurnal corticosterone rhymicity is moderately altered as seen at a functional level via HPA-profiling and on molecular levels through altered gene expression in the circadian entrainment pathway. Differential gene expression in brain regions relevant to stress regulation shows an enrichment for pathways linked to neural communication and brain disorders. Many of the differentially expressed genes are correlated with *Fkbp5* levels. In the tests utilized here, the impact of *Fkbp5* SNPs and ELA was greater in females than males.

These stronger effects of *Fkbp5* × ELA in female compared to male mice match previously reported sexual dimorphism in responsiveness to ELA in animals [16] and was discussed in humans [17]. Notably, ELA and sex hormones both influence maturation kinetics and thus the development of cerebral regions implicated in glucocorticoid regulation [18]. The interactions of the SNP rs1360780, sex and ELA observed in the mice presented here and in humans [11, 12] could thus be explained by the regulatory capacity of *Fkbp5* on glucocorticoid signaling. Based on sex-dependent correlations between *FKBP5* levels and depression and anxiety scores as well as with nadir cortisol levels, *FKBP5* was suggested as a female-specific biomarker for prolongued cortisol load and the associated risk of psychiatric disorders [19]. In line with this correlation, we observed associations between genotype and nadir corticosterone levels in *Fkbp5*-humanized mice, with AT-allele carrying females displaying higher morning glucocorticoid levels than CG-allele carrying females. The sexual dimorphism in the effect of ELA indicate that the novel *Fkbp5*-humanized mouse model offers the possibility to further investigate the networking of ELA, sex and disease-related SNPs.

In addition, the data provide mechanistic insights into how *Fkbp5* SNPs may contribute to the shaping of overall physiology and the stress response system. As negative modulator of glucocorticoid receptor maturation, *Fkbp5* holds the potential to inhibit glucocortocid signaling. At the same time, its expression depends on recent glucocorticoid exposure since *Fkbp5* itself harbors glucocorticoid response elements [8]. The higher induction of the AT-allele in CNS cell types of *Fkbp5*-humanized mice upon glucocorticoid stimulation could thus be expected to result in stronger or longer inhibition of subsequent glucocorticoid signaling [7]. *In vivo*, this stronger induction of inhibitory potential via *Fkbp5* in AT-allele carriers could lead to dampened negative feedback to the HPA-axis and a prolonged interval of elevated glucocorticoid levels. The negative feedback loop is furthermore critical for the maintenance of oscillation patterns and function [20]. The reduction in the complexity of ultradian fluctuation and the resulting decreased variability of HPA-axis reactivity in AT-allele carriers could decrease their flexibility to respond to novel environments. Behavioral evidence of this differential responsiveness could include the alterations in light-dark box testing, locomotor habituation, and abnormal social behavior as seen in this study. In humans, differences in HPA-axis responsivess to environmental stimuli, e.g. in the Trier Social Stress Test, between human AT- and CG-allele carriers has been demonstrated [10]. The findings imply that *Fkbp5* genotype dependent regulation of ultradian HPA-axis activity might be a core molecular mechanism that contributes to the variability seen in human stress responsiveness, which ultimately plays a role in distinction between healthy adaptation or pathological alteration in the aftermath of stress [21].

Another environmental stimulus that can affect glucocorticoid rhythms is the light-dark cycle [22]. One commonly investigated manifestation of this circadian rhythmicity is the pronounced increase of glucocorticoids prior to awakening [23]. Mechanistically, the ability to detect light in the retinal ganglia and to signal this via the suprachiasmatic nucleus to the periphery is a crucial trigger for the awakening response [24]. In AT- vs. CG-allele carriers, flatter diurnal glucocorticoid profiles were paralelled by lower expression of circadian entrainment related genes even though histological analyses of the eyes (data not shown) indicated no differences in the ability to detect light. This underscores the relevance of self-maintaining feedforward and feedback loops in regulating overall physiology throughout the day. While external light signals can synchronize individuals to a 24 h cycle [25], the internal gene expression driven clock seems to define the shape of the circadian glucocorticoid profile and thus when and how strong individuals are likely to respond to challenges. In humans, modulation of the cortisol awakening response was reported to influence their performance during the upcoming day and was dependent on the anticipation of challenges [26]. The awakening response is used clinically to identify individuals with certain personality traits that are vulnerable to develop psychiatric disorders [27], and for the diagnosis of depression [28].

Besides impaired awakening responses, differences in kinetic and responsiveness of the HPA-axis, e.g. to acute stress or dexamethasone exposure, between psychiatric patients and healthy controls have been demonstrated [29]. In the present study, no dysfunction of HPA-axis responsiveness was observed, which indicates that the combination of ELA and genetic predisposition via the AT-allele of *Fkbp5* alone might not be sufficient to cause full pathology. This is in agreement with the Research Domain Criteria framework proposing a continuum between ‘normal’ and ‘pathological’ which needs to be better understood in order to alleviate symptoms. Accordingly, the transition to pathology occurs over a life time and is a multidimensional process shaped by numerous genetic and environmental factors that introduce subtle changes which jointly alter networking of physiological systems [30]. As in humans, the *Fkbp5*-humanized mouse model demonstrates changes in basal HPA-axis activity dependent on genotype and early life experience, with more prominent effects in females than males. These alterations in non-stimulated HPA-axis functioning were suggested to have an impact on sleep-wake states, responsiveness to environmental stimuli and *vice versa* [31]. In the long run, insufficient adaptation could contribute to allostatic load and finally development of disorders [21]. However, the cumulative stress load in this study was low since the animals were not exposed to any severe or chronic stressors during later life.

Nevertheless, the *Fkbp5* × ELA model shows indications of changes in the psycho-immune-neuro-endocrine system that are commonly seen in response to chronic stress. Reduced expression of immediate early genes as markers of plasticity in the prefrontal cortex and hippocampus as well as elevated mitochondrial respiration in response to repeated mild stress during adulthood was previously reported [32]. In the present study, the increased expression of genes related to oxidative phosphorylation in the hippocampus of AT- vs. CG-allele carriers is an interesting parallel, as is the reduction of genes related to synaptic communication. Reduced neural communication and plasticity might become maladaptive since dendritic retraction has been described to render the hippocampus more vulnerable to neurotoxic or metabolic challenges [33, 34]. The longer the time window of decreased plasticity and increased vulnerability exists, the higher is the likelihood of a co-incidential high metabolic demand. Stressful situations only transiently elevate energetic demands while simultaneously decreasing the neuronal supply with glucose [35]. Unique stress events may thus not cause irreversible harm to the hippocampus, and AT-allele carriers might even benefit from their inherent higher expression of mitochondrial genes. Under prolonged exposure to glucocorticoids, increased oxidative phosphorylation in AT-allele carriers might produce excessive amounts of neurotoxic reactive oxygen species which may damage the hippocampus. Findings of this study imply more glucocorticoid signaling in the hippocampus of AT-relative to CG-allele carriers since the glucocorticoid signaling inhibitor *Fkbp5* had a lower expression level while nadir corticosterone levels were increased in female AT- vs. CG-allele carriers. Cumulatively, this mechanism could contribute to the loss of hippocampal volume in stress-related disorders such as depression and would explain why AT-allele carriers are more prone to develop disorders than CG-allele carriers [36]. The proposed sequence of alterations on cellular and circuitry level from healthy to allostatic load and allostatic overload conditions is outlined in Fig. 6. Assessment of behavior and physiologic read outs in *Fkbp5*-humanized mice that experienced both, ELA and more severe or chronic stress paradigms, would resolve these questions.

**Figure 6:**
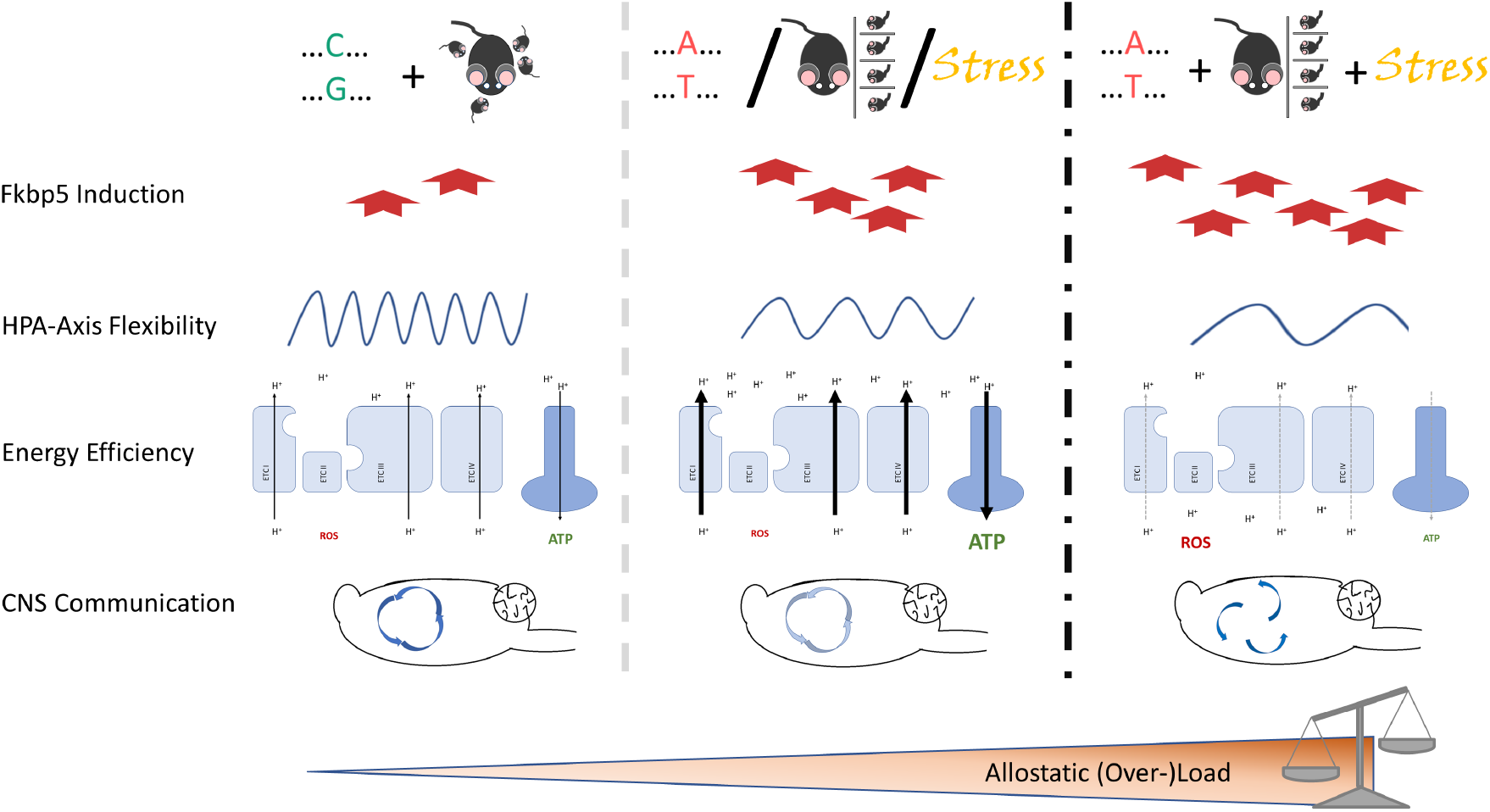
Proposed Sequence of Alterations in the Stress Response System on Cellular and Brain Circuit Level in Health, Allostasis and Allostatic Overload. The normal induction of *Fkbp5* upon challenge in CG-allele carriers with undisturbed maternal care allows for dynamic ultradian and circadian rhythms of the HPA -axis (**left**). In parallel, the electron transport chain (ETC) in the mitochondrial membrane produces energy in the form of adenosine-tri-phosphate (ATP) and few reactive oxygen species (ROS), while brain regions involved in stress regulation such as hypothalamus, hippocampus, pre-frontal cortex and amygdala engage in interconnected communication. Carriers of the AT-allele, or individuals exposed to early life adversity or mild chronic stress show signs of allostatic load (**center**). The affected individuals display a higher induction of *Fkbp5* and an attenuated rhythmicity of the HPA-axis. The associated increase in nadir glucocorticoid levels is linked to higher expression of genes related to oxidative phosphorylation, resulting in elevated mitochondrial respiration and ATP production, and to a lower expression of genes involved in synaptic communication. In the proposed triple-hit condition, a further increase in the levels of *Fkbp5* could interfere with the negative feedback to the HPA-axis and delay the termination of the stress response (**right**). As consequence of prolonged stress, the ETC might suffer from wear and tear resulting in a decreased efficiency in ATP production combined with elevated ROS generation and oxidative stress. Morevover, the reduced communication between stress-regulating brain regions could manifest in uncoupling of the brain circuits and asynchronous neural signaling. The here described *Fkbp5*-humanized mice will support future work to validate this scenario.

Moreover, the combination of *Fkbp5*-SNPs and ELA with simultaneous or sequential stress hits could enable prediction of and intervention at critical transition points during the development and progression of psychiatric symptoms.

## 4 Conclusion

The cumulative load of genetic predisposition, unfavorable environmental influences during development and repeated exposure to stressful events increases the prevalence of psychiatric disorders in affected individuals. The glucocorticoid-induced expression of *Fkbp5* is a hub for integrating lifetime and recent stressful experiences. Simultaneously, *Fkbp5* modulates responsiveness to acute stressors as negative modulator of glucocorticoid signaling. The naturally occurring *Fkbp5*-SNPs in laboratory rodents do not feature comparable functional effects as rs1360780 in humans, where the AT- vs. CG-allele is more strongly induced by glucocorticoids and linked to the etiology of psychiatric disorders. To enable studying in more detail the mechanistic impact of the human SNP on stress physiology and the etiology of psychiatric disorders, *Fkbp5*-humanized mouse lines carrying either the AT- or CG-allele of this SNP were generated. Characterization of the *Fkbp5 ×* ELA mouse model showed mechanistic and face validity with aspects of psychiatric disorders. Female AT- vs. CG-allele carriers after ELA showed attenuated diurnal rhythmicity of glucocorticoids, lower activity, and less responsiveness to novel environments. On a molecular level, reduced expression of genes related to circadian entrainment and synaptic communication as well as increased expression of genes related to mitochondrial respiration between AT- vs. CG-allele carriers imply a genetic predisposition of their psycho-immune-neuro-endocrine system to allostatic changes reported in a mild chronic stress settings. Since ELA lead to decreased circadian entrainment in the hippocampus, which in turn influences the circadian entrainment in the hypothalamus, the combination of ELA and *Fkbp5* SNPs could synergistically modify the HPA-axis to respond less to stimuli. Given that dynamic variability in glucocorticoid levels and plasticity are required for adaptation to challenges, this predisposition increases the risk of an unsuccessful resolution of allostatic loads and thus elevates the risk of developing stress-related disorders. In combination with severe or chronic stress exposure, the observed *Fkbp5* × ELA interactions likely contribute to the etiology of stress-related pathology. Taken together, we are confident that this novel animal model will contribute to more comprehensive analyses of *FKBP5*-induced alterations in the stress response network that causally lead to the development of pathology.

## 5 Methods

### 5.1 Generation of Transgenic Mice

Two novel transgenic mouse models were created by and are publicly available at Taconic Biosciences, carrying either the cytosine (C)/guanine (G) variant at position 3622 in the human *FKBP5* gene (C57BL/6NTac-Fkbp54571 (FKBP5) Tac) or the high-induction adenine (A)/thymine (T) version of rs1360780 (C57BL/6NTac-Fkbp5tm4570 (FKBP5) Tac). In short, the murine *Fkbp5* was exchanged with the human *FKBP5* coding region, keeping the 3’ and 5’ UTR of the mouse. Full details of the method are available at [7]. Homozygote mice were bred in-house to be used in the experiments.

### 5.2 Animal Husbandry

Standard laboratory conditions were adhered to (20-24°C, 45-55% humidity, 12-hour light/dark cycles (sunrise 6:00, sunset 18:00), *ad libitum* access for standard laboratory chow and water, enrichment (wooden block, red plastic shelter house and tube and paper stripes) provided). Mice had not been used in any other study prior to sacrifice. All animal experiments were performed under allowance of the regional council for animal welfare (Regierungspräsidium Tübingen, Baden-Württemberg, Germany, licence VVH 17-009) and in compliance with directive 2020/63/EU and ARRIVE guidelines. An overview of the study cohort is provided in Tab. 1. Sample size estimation was based on empirical knowledge regarding measurement accuracy or reproduction errors of *in vivo* and *ex vivo* methods (noise) and the expected effect sizes. Since we are the first to characterize these novel transgenic mice, no data was available for exact effect size estimations. Instead, published and in-house observed effect sizes of stress manipulations that were deemed to be biologically relevant were used as reference. As result of these considerations and based on our previous experience with animal experiments, n = 8 was agreed upon to be a suitable trade off between power for effect detection and the amount of animals required. Scheduled mating was used for breeding of the animals. On the day of birth, the litters were assigned to control or maternal separation in a way that group sizes were balanced as far as possible. Sex balance and equal litter sizes were not enforced since culling of littermates / offspring would introduce counfounding stressors and unnecessary suffering. As a result and since litter sizes and sex ratio within litters are non deterministic, final group sizes varied. After testing for equivalence, the WTs of the AT- and CG-allele strains were pooled which resulted in double the number of the transgenic groups.

### 5.3 Maternal Separation

Separation from mothers and litter mates at different times of the day for three hours starting from post-natal day two until 21 was carried out. During separation, heating pads were placed below the cages to keep the pups warm despite the lack of nesting material and low amount of saw dust. Mothers were kept at the other side of the room in their home cages. Maternal behaviour during separation and after reunion was observed to decrease after the first few separation sessions. Blinding of the experimenter was not possible given these obvious manipulations. After weaning on post-natal day 21, adolescent mice were group-housed until an age of 6 weeks. Afterwards, if not required earlier due to aggressive behavior, animals were single housing.

### 5.4 Behavioral Test Battery

Locomotion, dark-light preference, sociability and spatial working memory was assessed after mice were grown up to young adults. An overview of the timelines of the behavioral test battery is provided in Fig. 1. Within measurement sessions, mouse strains were mixed but controls and ELA-exposed mice as well as males and females were separated to avoid olfactory or auditory cues being transferred between groups, introducing distress or confounding effects. The experiments were first performed in males (controls, then ELA-exposed) and subsequently in females. Assignment to arenas or order of measurements within the day was randomized and for the social chamber and T-maze test experimenters were blinded regarding test groups. Arenas were extensively cleaned between each measurement and between groups. One day pause was kept to eliminate potentially distracting olfactory cues from the room.

#### 5.4.1 Open Field Test

A maximum of 30 arenas of 45 by 45 cm size were evenly illuminated with 267 lx, water gel and food pellet were placed around the borders of the center zone and one handful of saw dust from each mouse’ home cage were distributed inside the arena. Mice were brought to the measurement room at least two hours before the session for recovery from transport and habituation to the room. Shortly before 17:00, the actimot (TSE Systems, Bad Homburg, Germany), which detects movement of animals via breaking of light beams in x, y, z direction, was switched on, mice were placed in the middle of the measurement chambers and their locomotion was recorded over night until 07:00 to obtain measures of their activity in the light and dark phase (lights off 18:00, lights on 06:00).

#### 5.4.2 Dark Light Test

Measurement of preference for the dark or light compartment were also performed with the actimot system, while a 1/3 of the arena was kept dark (2.2 lx) using a black plastic house with a circular door to allow mice to freely travel between both compartments. At beginning of the measurement, mice were placed in the dark compartment facing the corner away from the door. Experiments were performed between 08:00 to 11:00 in the morning and lasted for 30 minutes.

#### 5.4.3 Three Social Chamber Test

Sociability was tested in an arena divided into three compartments of equal size (60 × 40 × 22 cm, Ugo Basile) with sliding doors between the compartments. The left and right section contained a mesh cylinder (7 cm in diameter, 15 cm height). Mice were habituated to the measurement room 1 hour prior testing and experiments were performed in the morning from 7:00-12:00. The arena was evenly illuminated with 23 lx. After recording the 5 minutes habituation phase of the tested mouse to the arena, an unfamiliar stimulus mouse was placed in one of the cylinders. Choice of side was evenly distributed across groups. Stimulus mice were juvenile, of the same sex as the tested mouse, habituated to the cylinder and used twice per day with 1-hour break between measurements. Behavior of the tested mouse was videotaped for 10 minutes and analyzed using an automated tracking software (TopScan CleverSys Inc., USA). Main readouts were the time spent, the amount of entries, the activity during the visit measured by distance and speed, the latency to first enter and the latency to end the first visit. This was assessed for the chamber as a whole and a zone surrounding the cylinders. As secondary readouts, locomotion and immobility were measured.

#### 5.4.4 Alternations T-Maze

Mice were moved to the testing room the day before their performance in the T-maze was assessed. At beginning of the measurement, the mouse was placed in the starting box for 5 seconds before the door to the arena was opened. In the first trial, the animal was either forced to enter the right or the left arm by closing the door to the respective other arm. Starting sides were evenly distributed across sexes, early life conditions and mouse lines. Every mouse was tested once and had to complete 14 trials consisting of entering one arm, closing the door to the non-chosen arm, returning of the animal to the starting zone and opening all passages to enable free choice of side for the next trial. The maximal allowed duration was set to 14 minutes and if an animal completed less than 7 trials it was excluded from further analysis. This was the case for 1 CG-allele carrying male control. Dimensions of the arena were 20 cm height, 8.5 cm corridor width, 30 cm lengths of each arm, 54 cm length of the starting zone. The test room was illuminated with 230 lx while above the T-maze light intensity was set to 50 lx. To enable spatial discrimination, navigation objects with differing shape and color were placed outside of the left and right arm of the arena.

### 5.5 HPA-Axis Performance

Blood was sampled from the *vena saphena* by immobilizing the mouse (Broome Rodent Restrainers, Harvard Apparatus, Cat.No.52-0460, MA 01746, USA), shaving and anointing the left leg and stinging into the vein with a lancet (Solofix, B. Braun, Cat.No. 6182003, Melsungen, Germany). Blood droplets were collected in K2-EDTA-containing capillaries (Microvette, Sarstedt, Cat.No.16.444, Nümbrecht, Germany) and stored on ice prior to centrifugation at 20000 g for 20 min at 4°C. The whole process from cage opening to collection of the last drop was carried out within less than 1.5 minutes to avoid a procedure-associated rise of corticosterone [37]. Plasma aliquots were frozen immediately.

For tracking of basal diurnal rhythmicity, blood was collected in the morning (06:45 – 07:15), afternoon (12:45 – 13:15) and evening (18:45 – 19:15). To obtain a measure of plasma corticosterone levels after stress, on another day at about the same time when the morning blood sample was drawn, mice were kept inside the restrainer for 5 minutes before puncturing the vein. Negative feedback to the HPA-axis was investigated by comparing rise or fall of plasma corticosterone levels between morning and 6 hours after injection of saline (NaCl 0.9%, B. Braun, Cat.No. FREU950) or with 0.001 mg/kg dexamethasone (DexaHexal 4 mg/ml diluted in saline, Hexal, Holzkirchen, Germany) two days after saline injection. An overview of blood sampling time points is provided in Fig. 1. Concentrations of corticosterone were quantified using an enzyme-linked immunosorbent assay (DetectX Corticosterone Enzyme Immunoassay Kit, Cat.No. CEA540Ge, Abor Assays, TX 77494, USA) following the manufacturer’s instructions. The data on morning corticosterone levels showed strong accordance between replicates and was therefore pooled for each mouse.

### 5.6 Gene Expression

#### 5.6.1 Tissue Collection

Mice were sacrificed in the morning under isoflurane anesthesia by rapid decapitation. Organs were collected within 10 minutes after death and immediately stored in cooled RNAlater or for histology in 4% formaldehyde supplemented with 20% sucrose over night.

#### 5.6.2 Generation of hiPSCs

Lines were derived from healthy patients genotyped for FKBP5 SNP rs1360780 and *FKBP5* InDel rs9470080CNV. In both cases, the AT genotype corresponds to the ‘high induction’, or ‘risk’ allele, while the CG genotype corresponds to ‘low induction’ or ‘resilience’ allele. Lines were derived from peripheral blood mononuclear cells collected from 2 females and 2 males homozygote for the AT- or CG-allele, with even distribution of both genotypes. Reprogramming was performed with episomal plasmids [38]. Comparison of the genome wide CNV in the parental material and the emanated hiPSC showed no chromosomal aberrations. Pluripotency markers were detected immunocytochemically.

#### 5.6.3 hiPSC differentiation

All hiPSC lines were cultured in mTSER1 (Stem Cell Technologies, Cat.No. 058509) on Matrigel Matrix High Concentration (Corning, Cat.No. 354263). Neural induction was performed based on a published protocol [39] with a few modifications. hiPS cells were maintained in Matrigel coated vessels, with mTeSR1 media and split by passing complete colonies using a non-enzymatic approach (EDTA, Versene Solution). hiPSCs were dissociated to single cells with Accutase (Stem Cell Technologies, Cat.No. 07920) and plated at 3 × 106 cells/well to allow the embryoid body (EB) formation in Neural Induction Media (NIM) + 10 *μ*M Y-27632 (Stem Cell Technologies, Cat.No. 72308). They were allowed to attach for at least overnight, and then the medium was replaced by NIM (without Y-27632), consisting of a 1:1 mix of N2 supplement (Life Technologies, Cat.No. 17502048) in DMEM/F12 (Life Technologies, Cat.No. 31331028) and B27 supplement (Life Technologies Cat.No. 17504044) in Neurobasal (Life Technologies Cat.No. 21103049), supplemented with 10*muM* SB431542 (Millipore, 616461) and 1 *μ*M Dorsomorphin (Tocris Bioscience, Cat.No. 3093). In days in vitro (DIV) 1-4, NIM was replaced twice a day. On day 4, the EB suspension were made and moved with a 5 ml serological pipette into a 6-well Clear Flat Bottom Ultra Low Attachment Multiple Well Plates (Corning, Cat.No. 3741), and cultured in NIM replaced daily for 10 days. On 10 DIV, the EBs were plated on tissue culture plates coated with Matrigel. On DIV 14-16, the neuroepithelial sheet was detached from the plate using STEMdiff Neural Rosette Selection Reagent (Stem Cell Technologies, Cat.No. 05832). From the following day until DIV 27 cultures were grown in Neuronal Maintenance Media (N2B27 supplemented with 20ng/*μ*l hFGF) replaced daily or on alternate days. Between DIV 17 and 30, any non-neural differentiation present was removed by passaging with STEMdiff Neural Rosette Selection Reagent, and the neural cultures were then dissociated to single cells using Accutase. When cultures reached 80%–90% confluency, they were passaged again until a final passage between DIV 33-40, when they were plated for long-term culture, after which N2B27 medium was replaced every second day. At DIV 60 and 90, in each line, cells were detached with Accutase from several wells and filtered through 40 μm cell strainer (Corning, Cat.No. 352340) for FACS sorting evaluation.

#### 5.6.4 FACS sorting

The media from 4 wells per line, containing astrocyte-neuron co-cultures were removed and the RLT buffer was added to wells for the RNA extraction. In parallel, at least 1 well per line (limited by the number of wells containing differentiated cells, variable between lines) was proceeded for separation of astrocytes and neurons from co-cultures using the positive selection approach with anti-CD44 (BD-bIoscience, Cat.No. 555478) antibodies [40]. Cells were gently detached from the well surface with Versene (3-5 minutes at 37°C), to avoid the epitope damage. Mechanical dissociation with p1000 pipette (5 times gentle up- and down strokes) was applied for obtaining single cells suspension. Cells were counted and up to 3×105 cells were incubated in flow cytometry (FC) wash buffer consisting of 1% FBS, 1× penicilin-streptomycin, nuclease free water and RNasin Plus RNase inhibitor 0.2 U/*μ*l. Next, cells were incubated in the FC wash buffer (2h at 4 °C) containing FITC-coupled anti-CD44 antibody (1 to 80 BD Pharmingen Cat.No. 555478) or its isotype control (FITC Mouse IgG2b, *κ* Isotype Control, BD Pharmingen Cat.No. 555478). After incubation, cells were washed in wash buffer (RNAase-free PBS, pH 7.4 + 0.2U/*μ*l RNase inhibitor), spin down (300 g × 3’) and resuspended in 500*μ*l of wash buffer. Separation was performed on FACSAria Machine (ZMBH, Heidelberg) at 4 °C using 100 *μ*m nozzle (optimized for droplet stream). Based on side scatter pulse width and height (SSC-W and SSC-H), 200000-300000 events of singlets were sorted directly to Low binding tube coated with FBS overnight (Corning, CLS3207-250EA), spun down (400 g × 10’ at 4 °C) and the pellet was resuspended in 600*μ* of Qiazol Lysis Reagent (Qiagen, Cat.No. 79306) and kept at −80 °C before sequencing.

#### 5.6.5 Next Generation Sequencing

RNA was isolated using RNeasy Plus kit (Qiagen Cat.No. 74192) following the manufacturer’s recommendations. RNA purity was checked spectrophotometrically using the NanoPhotometer (IMPLEN, CA, USA) and QIAxpert(Qiagen). Concentration was measured using Qubit RNA Assay Kit in Qubit 2.0 Flurometer (Life Technologies, CA, USA), while integrity was assessed using the standard sensitivity RNA kit (Cat.No. DNF-471, Advanced Analytical) on a Fragment Analyzer (Thermo Fisher Scientific, Langenselbold, Germany) and RNA Nano 6000 Assay Kit of the Bioanalyzer 2100 system (Agilent Technologies, CA, USA). High quality RNA samples with RIN >7.5 were eligible for further processing.

A total amount of 1 *μ*g RNA per sample was used as input material for the RNA sample preparations. Enrichment of mRNA from eukaryotic organisms was performed using oligo(dT) beads from NEBNext Poly(A) mRNA Magnetic Isolation Module (Cat.No. E7490L, NEB, USA). Subsequently, sequencing libraries were generated using NEBNext Ultra II Directional RNA Library Prep Kit for Illumina (Cat.No. E7770L, NEB, USA) following manufacturer’s recommendations. Briefly, fragmentation was carried out using divalent cations under elevated temperature in NEBNext First Strand Synthesis Reaction Buffer (5×). First strand cDNA was synthesized using random hexamer primer and M-MuLV Reverse Transcriptase (RNaseH-). Second strand cDNA synthesis was subsequently performed using DNA Polymerase I and RNase H. In the reaction buffer, dNTPs with dTTP were replaced by dUTP. Remaining overhangs were converted into blunt ends via exonuclease/polymerase activities. After adenylation of 3’ ends of DNA fragments, NEBNext Adaptor with hairpin loop structure were ligated to prepare for hybridization.

In order to select cDNA fragments of preferentially 250-300 bp in length, the library fragments were purified with AMPure XP beads (Cat.No. A63987 Beckman Coulter, Beverly, USA). Then 3*μ*l USER Enzyme (NEB, USA) was used with size-selected, adaptor-ligated cDNA at 37°C for 15 min followed by 5 min at 95°C before PCR. Then PCR was performed with Phusion High-Fidelity DNA polymerase, Universal PCR primers and Index (X) Primer. At last, products were purified (AMPure XP beads) and library quality was assessed using the Agilent High Sensitivity DNA Kit (Cat.No. 5067-4626) on the Agilent Bioanalyzer 2100 system (Agilent Technologies, CA, USA).

The clustering of the index-coded samples was performed on a cBot Cluster Generation System (Cat.No. SY401-2015, Illumina) using TruSeq PE Cluster Kit v3-cBot-HS (Cat.No. PE-401-3001, Illumia) according to the manufacturer’s instructions. After cluster generation, the libraries were sequenced on a NovaSeq 6000 Illumina platform using NovaSeq 6000 S2 Reagent Kit v1.5 cat. 20028314 -(300 cycles) and 150 bp paired-end reads were generated (minimum 12 Gb and 40 M).

RNA-Seq reads were aligned to the mouse genome using Hisat2 software, version 2.1.0 with the corresponding Ensembl GRCm38.p6 reference genome (http://www.ensembl.org). Confirmation of genotyping was done by aligning the NGS reads to the human reference genome GRCh38.p13. Sequenced read quality and duplications were checked with FastQC software, version 0.11.9 and alignment quality metrics were calculated using Samtools flagstat software, version 1.10. Gene and transcripts expression profiles were quantified using Cuffquant and Cuffnorm, version 2.2.1 and GTF file from the Ensembl (v. 100) database to obtain Fragments Per Kilobase Million mapped reads (FPKM). Library preparation, sequencing and initial data processing was carried out at Intelliseq (Poland).

### 5.7 Statistical Analyses

Data processing and analysis was carried out using R (version 4.0.2). For analysis of read outs with repeated measurements (open field, dark-light and social chamber test, HPA-axis performance), nested models using early life condition group, mouse strain and sex as between factors and compartment or time point as within subject component were defined and their quality was inspected visually and using one-point cross validation (R packages *nlme* and *afex*). Data without a temporal or spatial component were modeled linearly. Confidence intervals for the coefficient estimates were obtained using the non-centrality parameter method and the Greenhouse-Geiser method for approximation of the degrees of freedom was applied for nested models [41]. Analysis of variance and effect size estimation were performed using partial sum of squares type II (R packages *car* and *effectsize*). Besides the generalized eta squared (*gη*^2^), the partial epsilon squared (*pϵ*^2^) effect size were reported to reduce potential bias by small sample size [42, 43, 44]. In addition, the relative explanation of variance was assessed (R packages *MuMIn* and *r2glmm*). If significant model terms were suggestive, pairwise two-sided *post hoc* tests with Tukey contrasts were performed (package *emmeans*). Effect sizes in the descriptive analyses between subgroups were computed as Cohen’s d estimates using pooled variance.

Regarding data obtained from next generation sequencing, the following additional analysis steps were performed: The obtained data was filtered for tissue-wise median and mean expression to be above 1 FPKM. In addition, selection criteria for fold changes and signal-to-noise ratios bigger than ± 30% and ± 1.5 were applied, respectively. Comparisons were made between control and ELA-exposed females independent of strain as well as separate for the subgroups of AT- and CG-allele carriers. Furthermore, putative differences between AT- and CG-allele carriers independent of early life experiences, and within the control and maternally separated subset were investigated. To all transcripts where the row-wise t-test was significant, a false-discovery rate filter of 10% was applied and only genes of which the related transcripts indicated fold changes in the same direction were considered as differentially expressed genes (DEGs) in subsequent analyses. Based on the association of the CG-allele with resiliency and the AT-allele with risk to develop disorders, DEGs which were unique to CG-allele carriers when comparing effects of early life conditions (coping) were labeled as potentially resiliency-associated genes, while an overlap of DEGs from the early life comparison with DEGs from the SNP-comparison in the control subgroup were labeled as potential vulnerability-related genes. In addition, transcripts were the 2-way ANOVA suggested an interaction of early life condition × *Fkbp5*-genotype at an *α* level of 5% were included in gene set enrichment analyses.

Using Ingenuity (IPA, Qiagen), the FPKM values of the listed DEGs were subjected to ‘core analysis’ of ELA vs. control for the AT- and CG-allele carrying subgroup applying a threshold of absolute fold changes bigger than 1.5. The results of these ‘core analyses’ were entered into a ‘comparison analysis’ to investigate the deregulation of genes due to ELA between strains. The ‘comparison analysis’ was limited to the term ‘Diseases and Biological Functions’, and further limited to sub-categories of neurological relevance. The *z*-score *p*-value was set to <0.0001. The final comparison list was then filtered to the top 10 results, ordered by *z*-score.

Pearson correlations were computed tissue-wise for the normalized expression levels of *Fkbp5* and the DEGs with HPA-axis and behavior-related read outs. The cutoff for meaningful correlations was *a priori* set to > |0.6|. Among those correlations, the FDR was fixed to 5%.

Generally applicable gene-set enrichment analyses (package *gage*) for metabolic pathways listed in the Kyoto Encyclopedia of Genes and Genomes (KEGG) were performed and visualized in case of significant over representation (package *pathview*). For comparison to the hiPSC-derived astrocytes and neurons, the genes listed in the circadian entrainment pathway (hsa004713) were extracted, filtered based on the above-mentioned effect size and significance criteria and visualized without prior checks on pathway enrichment due to the limitations in sample size.

## 6 Accompanying Statements

## 6.1 Acknowledgements

The authors wish to thank their colleagues at Boehringer Ingelheim Pharma GmbH & Co KG Margot Weiland, Sonja Diehl, Nadine Richter, Marion Trautmann, Werner Rust, Birgit Stierstorfer, Tanja Schönberger for their excellent support in processing of the obtained samples as well as Silke Laack-Reinhardt, Yvonne Schneider, Sonja Hofbauer, Ralf Weber, Britta Gerth and Lukas Schmidt for their help with breeding of the animals. In addition, the hiPSC work would not have been possible without the exceptional support of Susanne Zach (Boehringer Ingelheim Pharma GmbH & Co KG), Shringarika Singh and Santiago Tena (BioMedX). We furthermore thank Michal Korostynski, Slawomir Golda, Dzesika Hoinkis and Marcin Piechota at Intelliseq for carrying out the next generation sequencing and discussing the thereof obtained data. Special thanks go to Michael Schuler for his support during the realization of the novel mouse lines together with Susie Mikkelsen at Taconic Biosciences. Lastly, the authors are greateful for the support of Elisabeth Binder at the Max Planck Institute for Psychiatry in Munich for her guidance in the conceptualization of the humanized mice and the provision with hiPSCs from patients and healthy controls.

## 6.2 Author Contributions

Allers, KA: conceptualization, maternal separation, data interpretation, manuscript revision

Blasius, A: T maze

Del Prete, D: cultivation, FACS and NGS of hiPSCs

Harris, I: support RNA isolation

Hengerer, B: conceptualization, revision of the manuscript

Kolassa, IT: data interpretation, revision of the manuscript

Koros, E: support with maternal separation

Nold, V: conceptualization and execution, sample and data analysis, manuscript

Peleh, T: support social chamber test

Portenhauser, M: support RNA isolation, corticosterone assay, social chamber test

Slezak, M: conceptualization hiPSC experiments, manuscript revision

## 6.4 Ethics Statement

Experiments were performed under the allowance of the regional council for animal welfare (Regierungspräsidium Tübingen, Baden-Württemberg, Germany) and adhere to ARRIVE guidelines.

## 6.5 Data Availability Statement

All raw data files and processed summary data frames will be made available by the corresponding author upon request. The accession code for the murine NGS data set on NCBI’s Sequence Read Archive is PRJNA743189. The hiPSC data sets will be made available upon request.

## 6.6 Code Availability Statement

All R code written to process, analyze and visualize the herein contained data will be made available by the corresponding author upon request.

## 7 Supplementary Material

**Supplementary Table S1:** Descriptive Corticosterone Concentration [ng/ml] in *Fkbp5*-Humanized and Wild Type Mice.

| Strain | Sex | Group | Time | N | Mean | SD | 95% CI<br>low | 95% CI<br>up |
| --- | --- | --- | --- | --- | --- | --- | --- | --- |
| AT | Female | Control | Morning | 6 | 40.50 | 11.90 | 28.01 | 52.99 |
|  |  |  | Noon | 6 | 99.50 | 55.30 | 41.46 | 157.54 |
|  |  |  | Evening | 6 | 109.83 | 49.63 | 57.75 | 161.92 |
|  |  |  | restraint | 5 | 135.20 | 28.49 | 99.82 | 170.58 |
|  |  |  | 6h Dex | 6 | 6.83 | 6.88 | -0.39 | 14.06 |
|  |  | ELA | Morning | 11 | 52.42 | 22.27 | 37.46 | 67.38 |
|  |  |  | Noon | 10 | 78.40 | 33.56 | 54.39 | 102.41 |
|  |  |  | Evening | 11 | 86.82 | 31.22 | 65.85 | 107.79 |
|  |  |  | restraint | 11 | 180.00 | 52.19 | 144.94 | 215.06 |
|  |  |  | 6h Dex | 10 | 9.10 | 12.82 | -0.07 | 18.27 |
| AT | Male | Control | Morning | 7 | 19.14 | 8.85 | 10.96 | 27.33 |
|  |  |  | Noon | 7 | 50.43 | 24.73 | 27.56 | 73.30 |
|  |  |  | Evening | 7 | 41.86 | 15.94 | 27.11 | 56.60 |
|  |  |  | restraint | 7 | 103.71 | 62.87 | 45.57 | 161.86 |
|  |  |  | 6h Dex | 7 | 4.71 | 6.58 | -1.37 | 10.80 |
|  |  | ELA | Morning | 5 | 16.73 | 2.97 | 13.05 | 20.42 |
|  |  |  | Noon | 5 | 48.40 | 18.04 | 26.01 | 70.79 |
|  |  |  | Evening | 5 | 44.40 | 22.71 | 16.20 | 72.60 |
|  |  |  | restraint | 4 | 73.75 | 21.27 | 39.91 | 107.59 |
|  |  |  | 6h Dex | 5 | 3.20 | 4.92 | -2.91 | 9.31 |
| CG | Female | Control | Morning | 7 | 28.86 | 13.87 | 16.03 | 41.69 |
|  |  |  | Noon | 7 | 35.43 | 13.48 | 22.96 | 47.89 |
|  |  |  | Evening | 7 | 109.86 | 53.76 | 60.14 | 159.58 |
|  |  |  | restraint | 7 | 147.14 | 77.04 | 75.89 | 218.40 |
|  |  |  | 6h Dex | 7 | 7.57 | 6.29 | 1.75 | 13.39 |
|  |  | ELA | Morning | 11 | 37.18 | 18.60 | 24.69 | 49.68 |
|  |  |  | Noon | 11 | 99.36 | 33.76 | 76.68 | 122.04 |
|  |  |  | Evening | 11 | 73.82 | 32.58 | 51.93 | 95.71 |
|  |  |  | restraint | 11 | 170.64 | 60.70 | 129.86 | 211.41 |
|  |  |  | 6h Dex | 11 | 5.55 | 9.20 | -0.64 | 11.73 |
| CG | Male | Control | Morning | 7 | 12.29 | 2.89 | 9.61 | 14.96 |
|  |  |  | Noon | 7 | 107.43 | 138.97 | -21.10 | 235.96 |
|  |  |  | Evening | 7 | 22.57 | 15.52 | 8.22 | 36.93 |
|  |  |  | restraint | 7 | 97.29 | 28.77 | 70.67 | 123.90 |
|  |  |  | 6h Dex | 7 | 1.00 | 0.00 | 1.00 | 1.00 |
|  |  | ELA | Morning | 12 | 17.64 | 7.17 | 13.08 | 22.20 |
|  |  |  | Noon | 12 | 45.75 | 12.20 | 38.00 | 53.50 |
|  |  |  | Evening | 10 | 31.40 | 19.13 | 17.72 | 45.08 |
|  |  |  | restraint | 12 | 102.92 | 30.43 | 83.59 | 122.25 |
|  |  |  | 6h Dex | 12 | 2.83 | 4.61 | -0.10 | 5.76 |
| WT | Female | Control | Morning | 20 | 29.39 | 12.97 | 23.32 | 35.46 |
|  |  |  | Noon | 20 | 96.35 | 72.41 | 62.46 | 130.24 |
|  |  |  | Evening | 19 | 137.05 | 81.53 | 97.76 | 176.35 |
|  |  |  | restraint | 19 | 152.58 | 39.29 | 133.64 | 171.52 |
|  |  |  | 6h Dex | 20 | 9.90 | 4.73 | 7.68 | 12.12 |
|  |  | ELA | Morning | 17 | 37.69 | 16.44 | 29.24 | 46.14 |
|  |  |  | Noon | 16 | 79.81 | 45.35 | 55.65 | 103.98 |
|  |  |  | Evening | 16 | 66.50 | 28.68 | 51.22 | 81.78 |
|  |  |  | restraint | 16 | 168.44 | 49.57 | 142.02 | 194.85 |
|  |  |  | 6h Dex | 17 | 8.12 | 6.49 | 4.78 | 11.45 |
| WT | Male | Control | Morning | 15 | 17.69 | 7.01 | 13.81 | 21.57 |
|  |  |  | Noon | 13 | 84.62 | 66.73 | 44.29 | 124.94 |
|  |  |  | Evening | 13 | 82.77 | 47.74 | 53.92 | 111.62 |
|  |  |  | restraint | 15 | 93.60 | 53.65 | 63.89 | 123.31 |
|  |  |  | 6h Dex | 14 | 2.79 | 5.03 | -0.12 | 5.69 |
|  |  | ELA | Morning | 16 | 22.79 | 8.19 | 18.43 | 27.15 |
|  |  |  | Noon | 17 | 81.00 | 34.87 | 63.07 | 98.93 |
|  |  |  | Evening | 17 | 31.88 | 18.55 | 22.34 | 41.42 |
|  |  |  | restraint | 16 | 102.88 | 64.52 | 68.49 | 137.26 |
|  |  |  | 6h Dex | 17 | 11.59 | 12.00 | 5.42 | 17.76 |

**Supplementary Figure S1:**
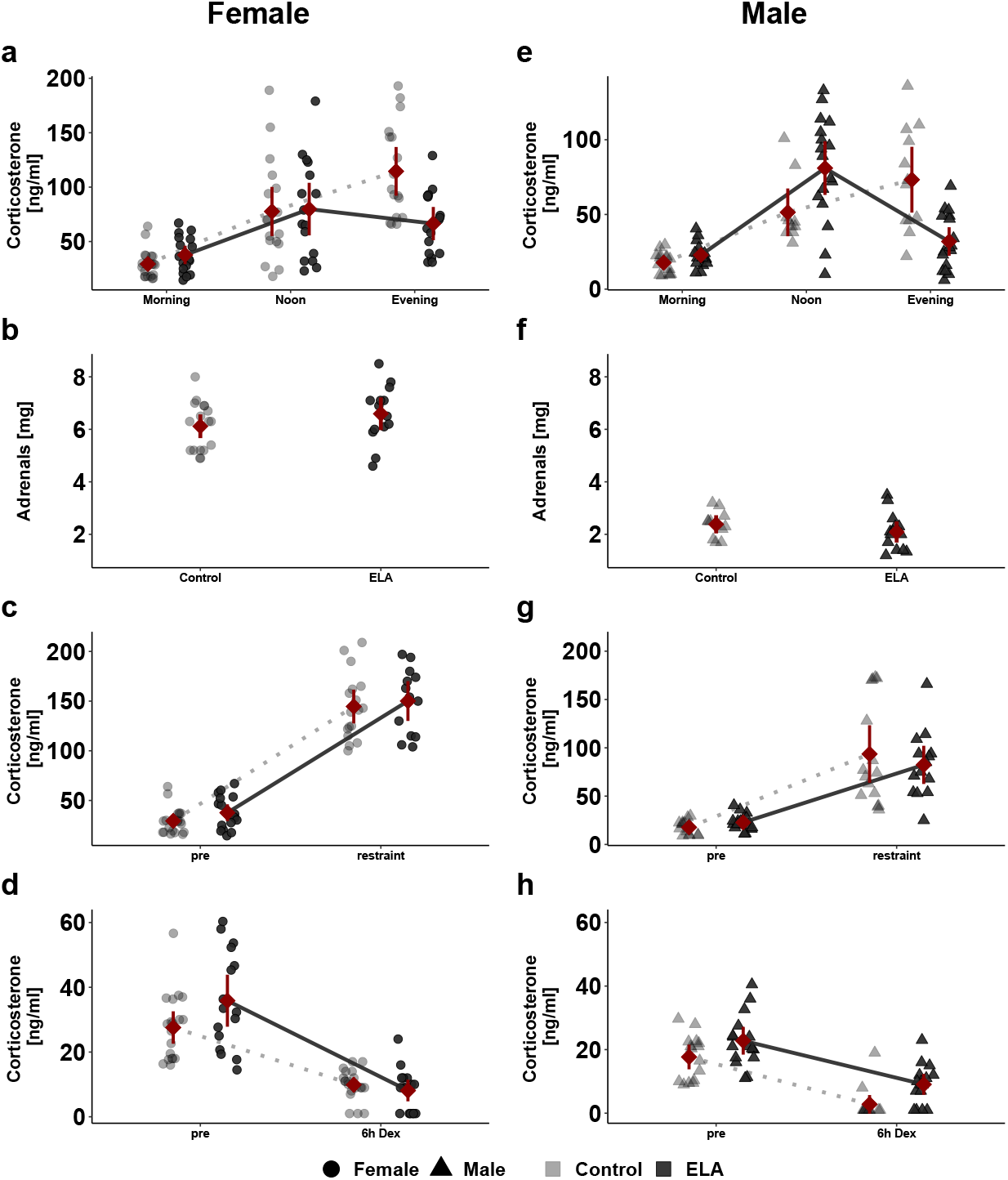
Plasma Corticosterone Levels in Wild Type Females Fluctuate more than in Males. Individual animal data (grey scale) is shown alongside with the mean ± 95% confidence interval (red). Diurnal rhythmicity of corticosterone plasma levels in female (**a**) and male (**e**) control mice peaks in the evening while at noon in ELA-exposed mice. A different scale for males than females was used to make the pattern better visible. Adrenal weights in wild type females (**b**) are higher than in males (**f**). Acute responsiveness of the HPA-axis to 5 minutes of immobilisation stress is higher in females (**c**) than males (**g**). Dexamethasone induced suppression of endogenous corticosterone 6 hours after injection is stronger in females (**d**) than males (**h**). Descriptive statistics, model summaries and ANOVA outcomes for diurnal, dexamethasone-suppressed and stress-induced corticosterone are provided in Tab. S1, Tab. S2, Tab. S3, Tab. S30, Tab. S31, Tab. S28 and Tab. S29, while Tab. S4, Tab. S5 and Tab. S6 contain the data for adrenal weight.

**Supplementary Figure S2:**
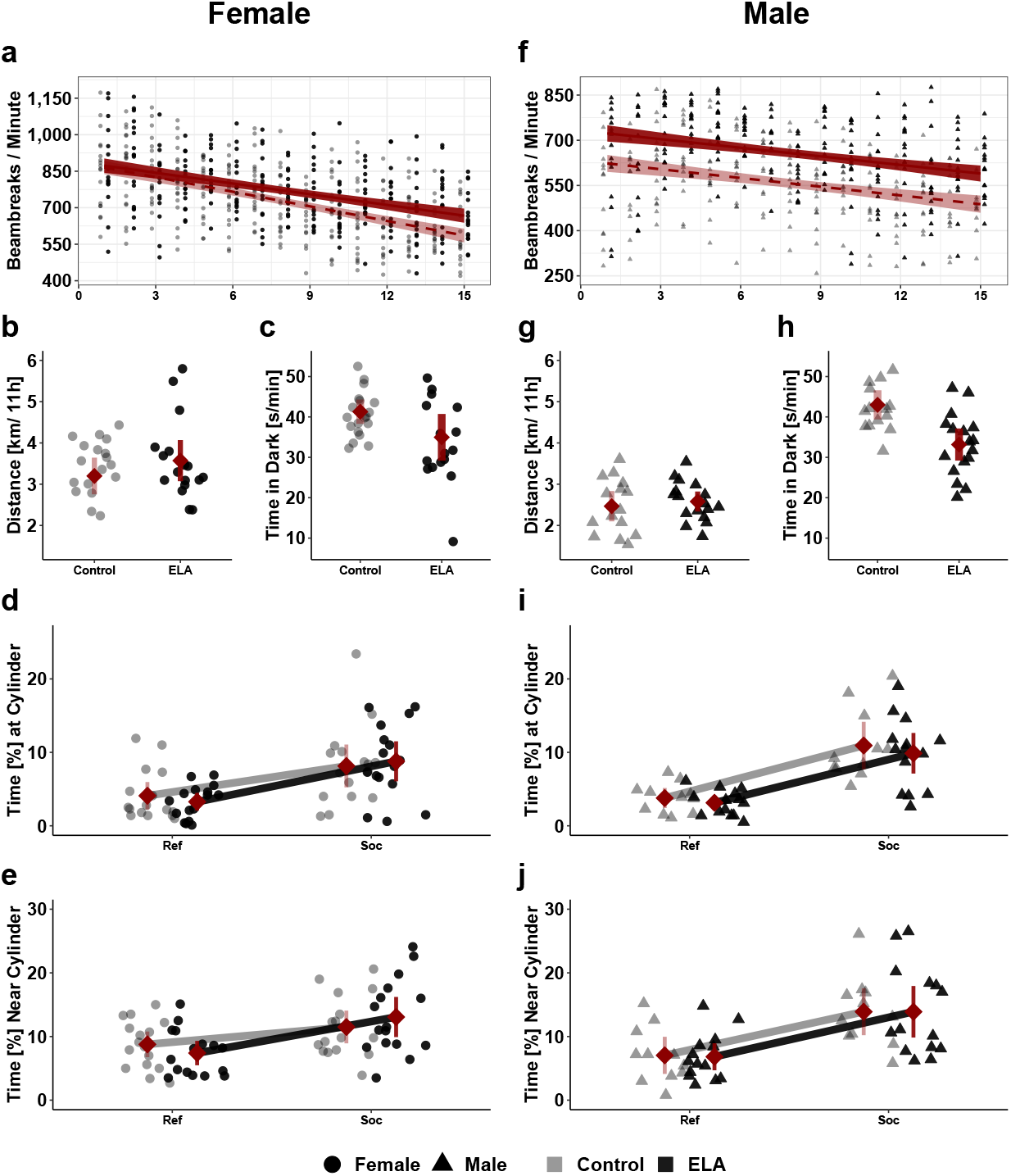
ELA Increases Exploration Behavior During Habituation in Wild Type Mice. Individual animal data is shown alongside with the mean ± 95% confidence interval (red). Increased breaking of light beams during the first 15 minutes in a novel environment in females (**a**) and males (**f**) with ELA. Females (**b**) run more distance during the night than males (**g**). Average time per minute spent in the dark compartment of the arena decreases after ELA with males (**h**) rather than females (**c**). Comparison of the percent of time females (**d**) and males (**i**) spent at the cylinder with (Soc) or without (Ref) an unfamiliar mouse separated for. Relative time [%] spent in the donut-shaped area surrounding the cylinder shown for females (**e**) and males (**j**). Descriptive statistics, model summary and ANOVA for the exploration activity are provided in Tab. S7, Tab. S8 and Tab. S9, for the nocturnal distance in Tab. S10, Tab. S11 and Tab. S12, for the Dark-Light-Test in Tab. S19, Tab. S20 and Tab. S21, for social interaction time in Tab. S22, Tab. S23 and Tab. S24, while for the time in social distance in Tab. S25, Tab. S26 and Tab. S27.

**Supplementary Figure S3:**
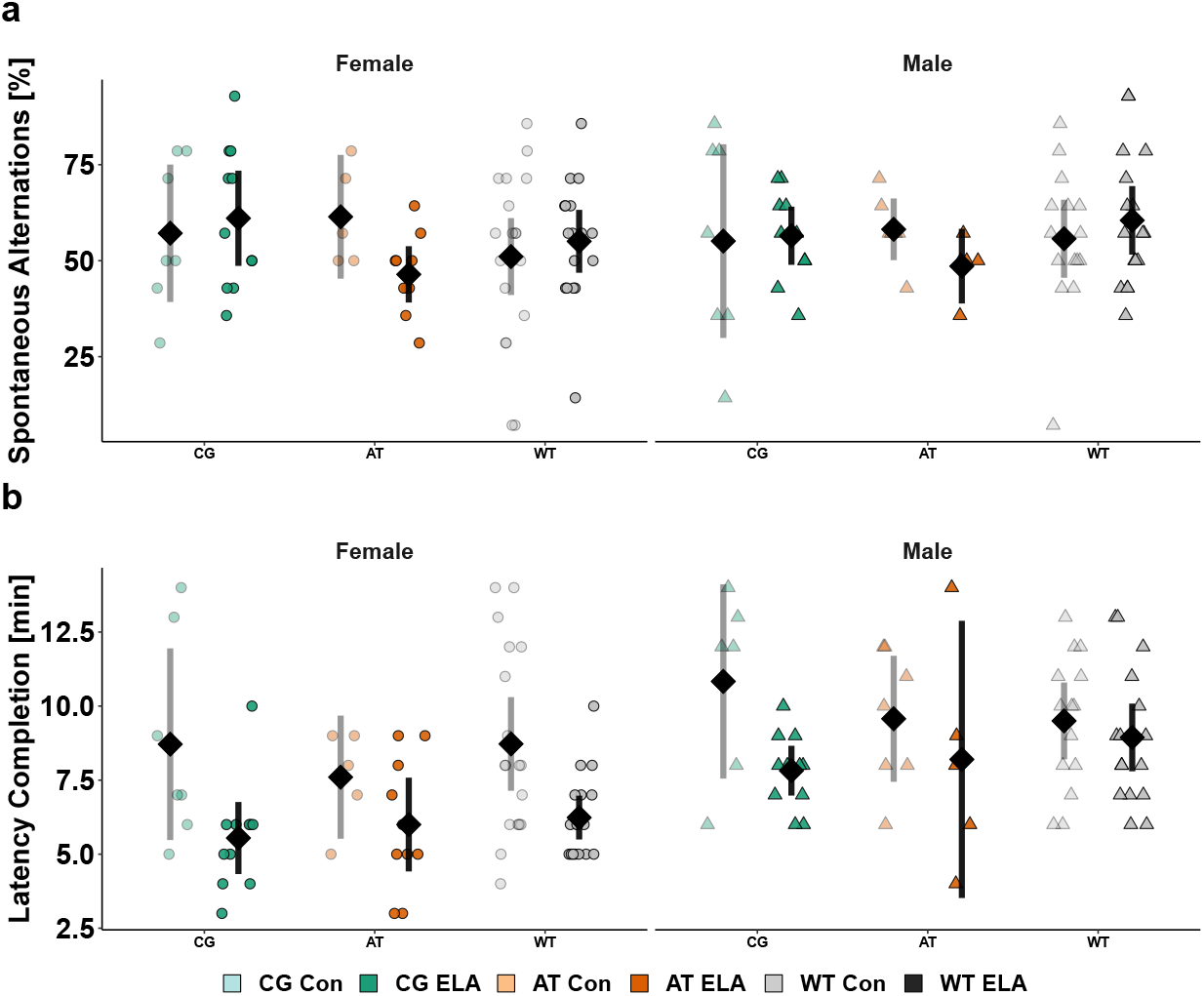
ELA Shortens the Time to Completion of the T-Maze Test in *Fkbp5*-Humanized and Wild Type Mice. Individual animal data is shown alongside with the mean (black diamond) ± 95% confidence interval. The percentage of spontaneous switching between arms of the T-shaped arena is performed at chance level with no statistically significant effect of Early Life condition, *Fkbp5*-genotype or sex (**a**). The time [min] needed to complete the 15 trials differed between controls and ELA-exposed animals and was further modulated by sex (**b**). Descriptive statistics, model summaries and ANOVA results are provided in Tab. S13, Tab S14, and Tab. S15 for the spontaneous alternations and in Tab. S16, Tab. S17 and Tab. S18 for the latency of T-maze completion.

**Supplementary Table S2:** Model Summary Diurnal Corticosterone.

| **Model Terms** | $\beta$ | *SE* | Df | $CI_{low}$ | $CI_{up}$ | *t* | *p* | $R^2_{part}$ |
| --- | --- | --- | --- | --- | --- | --- | --- | --- |
| Intercept | 12.29 | 15.57 | 235 | -16.95 | 41.52 | 0.79 | 0.4308 | 0.4283 |
| ELA | 5.35 | 19.59 | 123 | -31.61 | 42.32 | 0.27 | 0.7851 | 0.0502 |
| AT | 6.86 | 22.02 | 123 | -34.69 | 48.4 | 0.31 | 0.756 | 0.0266 |
| WT | 5.4 | 18.85 | 123 | -30.17 | 40.98 | 0.29 | 0.7749 | 0.0224 |
| Female | 16.57 | 22.02 | 123 | -24.97 | 58.12 | 0.75 | 0.4531 | 0.0223 |
| Noon | 95.14 | 21.09 | 235 | 55.53 | 134.76 | 4.51 | 0 | 0.0187 |
| Evening | 10.29 | 21.09 | 235 | -29.33 | 49.9 | 0.49 | 0.6263 | 0.0186 |
| ELA:AT | -7.76 | 31.07 | 123 | -66.39 | 50.87 | -0.25 | 0.8031 | 0.0163 |
| ELA:WT | -0.08 | 24.55 | 123 | -46.41 | 46.24 | 0 | 0.9973 | 0.0159 |
| ELA:Female | 2.97 | 27.94 | 123 | -49.74 | 55.68 | 0.11 | 0.9155 | 0.0144 |
| AT:Female | 4.79 | 31.78 | 123 | -55.18 | 64.75 | 0.15 | 0.8805 | 0.0118 |
| WT:Female | -4.87 | 26.13 | 123 | -54.17 | 44.43 | -0.19 | 0.8525 | 0.0115 |
| ELA:Noon | -67.03 | 26.54 | 235 | -116.88 | -17.19 | -2.53 | 0.0122 | 0.008 |
| ELA:Evening | 3.4 | 27.06 | 235 | -47.42 | 54.23 | 0.13 | 0.9 | 0.0079 |
| AT:Noon | -63.86 | 29.83 | 235 | -119.88 | -7.84 | -2.14 | 0.0333 | 0.0066 |
| WT:Noon | -28.45 | 25.88 | 235 | -77.05 | 20.15 | -1.1 | 0.2727 | 0.004 |
| AT:Evening | 12.43 | 29.83 | 235 | -43.59 | 68.45 | 0.42 | 0.6773 | 0.0031 |
| WT:Evening | 54.62 | 25.88 | 235 | 6.02 | 103.23 | 2.11 | 0.0359 | 0.0016 |
| Female:Noon | -88.57 | 29.83 | 235 | -144.59 | -32.55 | -2.97 | 0.0033 | 0.0016 |
| Female:Evening | 70.71 | 29.83 | 235 | 14.69 | 126.74 | 2.37 | 0.0186 | 0.0008 |
| ELA:AT:Female | 11.36 | 42.42 | 123 | -68.67 | 91.4 | 0.27 | 0.7892 | 0.0007 |
| ELA:WT:Female | 0.05 | 34.41 | 123 | -64.88 | 64.98 | 0 | 0.9988 | 0.0006 |
| ELA:AT:Noon | 67.41 | 42.1 | 235 | -11.65 | 146.47 | 1.6 | 0.1107 | 0.0005 |
| ELA:WT:Noon | 58.38 | 33.45 | 235 | -4.43 | 121.2 | 1.75 | 0.0822 | 0.0003 |
| ELA:AT:Evening | 1.55 | 42.43 | 235 | -78.13 | 81.23 | 0.04 | 0.9709 | 0.0002 |
| ELA:WT:Evening | -59.39 | 33.86 | 235 | -122.98 | 4.2 | -1.75 | 0.0808 | 0.0002 |
| ELA:Female:Noon | 122.64 | 37.85 | 235 | 51.56 | 193.72 | 3.24 | 0.0014 | 0.0002 |
| ELA:Female:Evening | -47.77 | 38.22 | 235 | -119.54 | 24 | -1.25 | 0.2126 | 0.0002 |
| AT:Female:Noon | 116.29 | 43.06 | 235 | 35.43 | 197.15 | 2.7 | 0.0074 | 0.0001 |
| WT:Female:Noon | 88.84 | 35.64 | 235 | 21.9 | 155.78 | 2.49 | 0.0134 | 0.0001 |
| AT:Female:Evening | -24.1 | 43.06 | 235 | -104.96 | 56.76 | -0.56 | 0.5763 | 0 |
| WT:Female:Evening | -27.91 | 35.71 | 235 | -94.96 | 39.15 | -0.78 | 0.4352 | 0 |
| ELA:AT:Female:Noon | -155.97 | 57.6 | 235 | -264.14 | -47.8 | -2.71 | 0.0073 | 0 |
| ELA:WT:Female:Noon | -138.62 | 46.82 | 235 | -226.54 | -50.7 | -2.96 | 0.0034 | 0 |
| ELA:AT:Female:Evening | 7.88 | 57.71 | 235 | -100.5 | 116.26 | 0.14 | 0.8916 | 0 |
| ELA:WT:Female:Evening | 24.51 | 47.16 | 235 | -64.06 | 113.07 | 0.52 | 0.6038 | 0 |

Model formula:
`lme( ~ Group*Strain*Sex*Time, random = ~ 1|ID/Time)`
$R^2_{marg} = 42.57\%$ , $R^2_{cond} = 95.17\%$
Abbreviations: $\beta$ = $\beta$ coefficient estimate, *SE* = standard error, Df = degrees of freedom, CI = 95% confidence interval, part = partial, group = early life condition group, marg = marginalized, cond = conditioned,

**Supplementary Figure S4:**
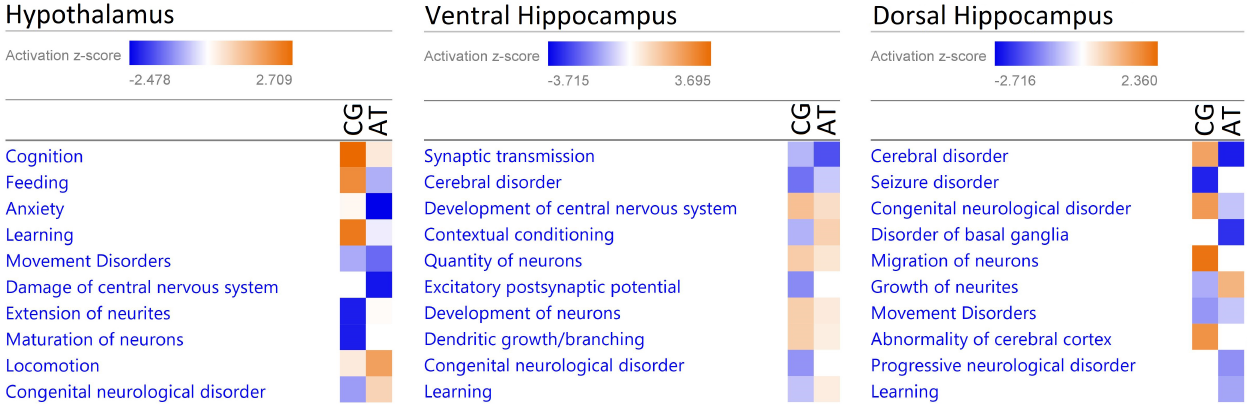
Comparison of ELA-Induced Changes in Neurological and Developmental Disease Pathways between *Fkbp5*-Genotypes. Clustered top 10 activation *z*-scores between CG-allele (left colum) and AT-allele (right colum) carriers in hypothalamus (**left**), ventral hippocampus (**middle**) and dorsal hippocampus (**right**).

**Supplementary Figure S5:**
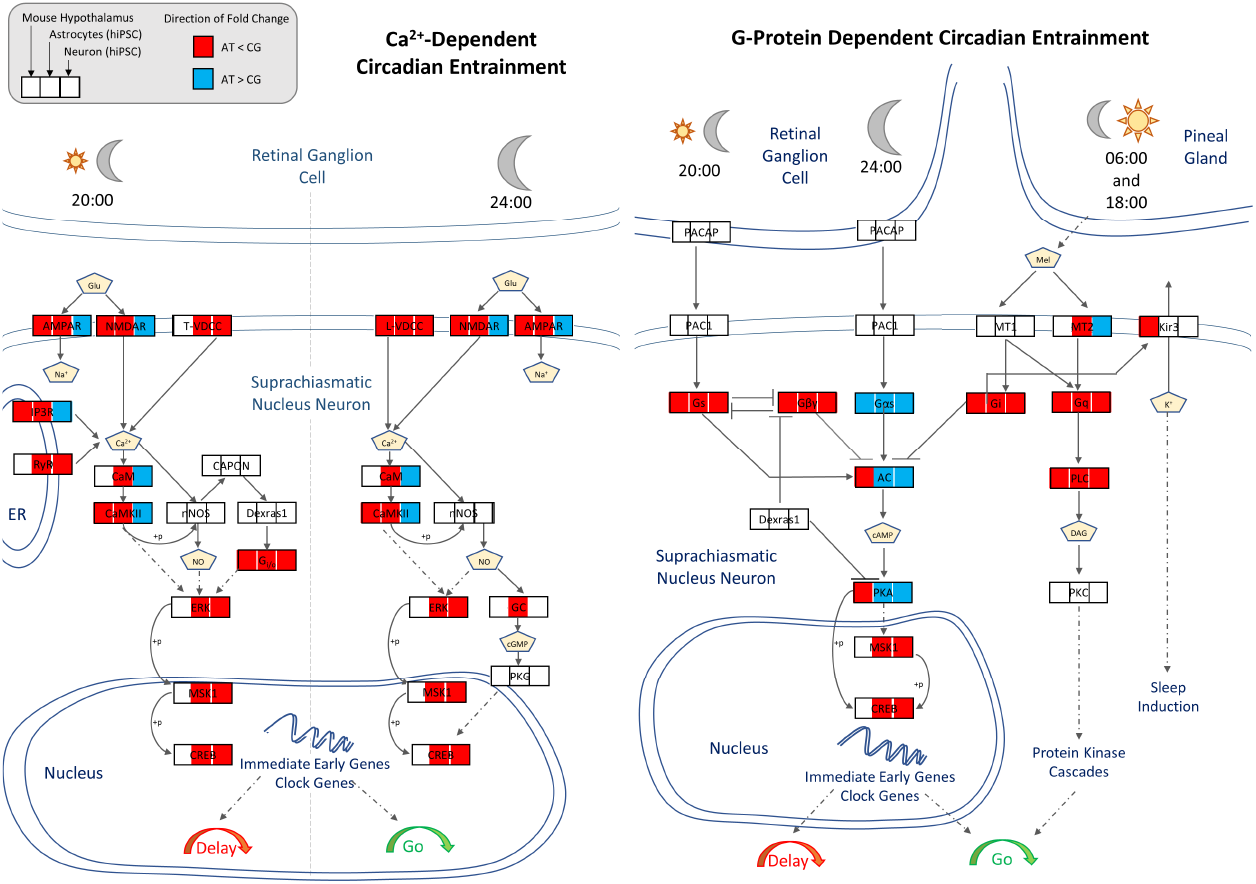
Differential Expression of Genes Related to Circadian Entrainment in *Fkbp5*-Mice and in hiPSC-Derived Astrocytes and Neurons of Human *FKBP5*-SNP Carriers. Visualization of significant fold changes between AT- vs. CG-allele shown for mouse hypothalamus, hiPSC-derived astrocytes and neurons. The calcium-dependent and the G-protein dependent arm of the circadian entrainment KEGG pathway is shown on the left and right, respectively.

**Supplementary Figure S6:**
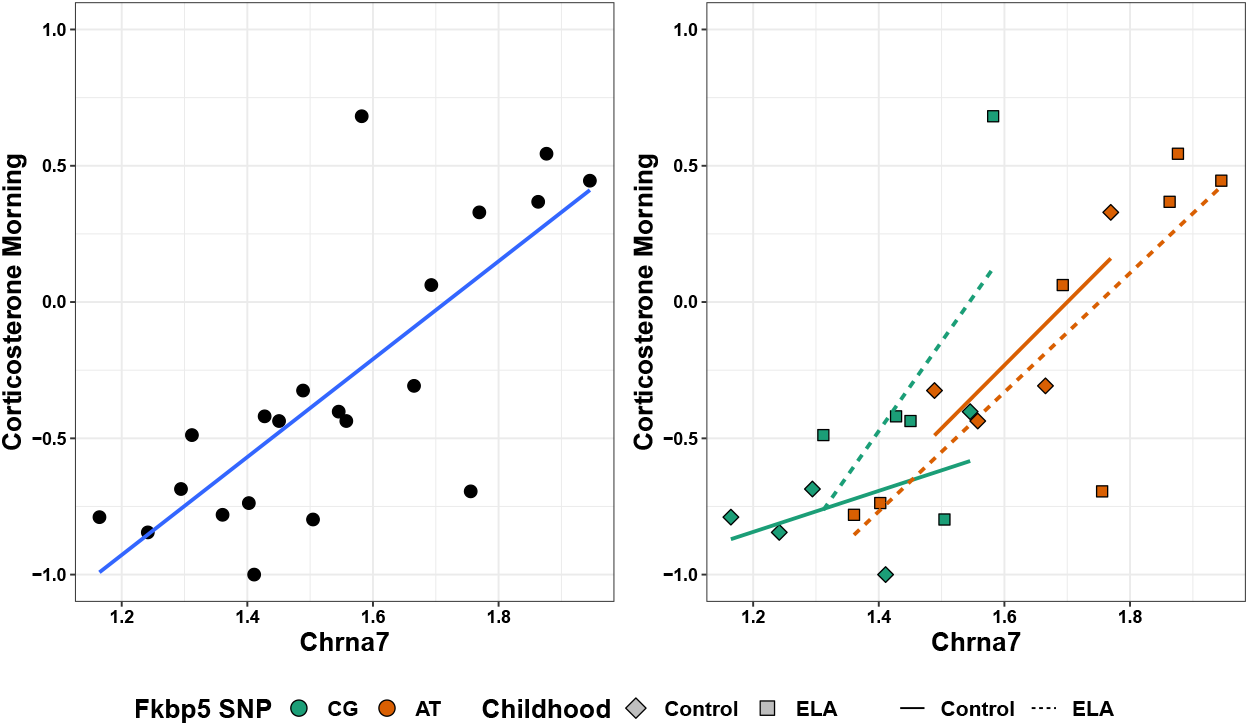
Regression of Morning Corticosterone Levels and *Chrna7*. Overall linear association (**left**) and regressions in the subgroups defined by *Fkbp5*-SNP and early life experience (**right**) are shown for the scaled morning corticosterone and log-transformed expression levels.

**Supplementary Table S3:**
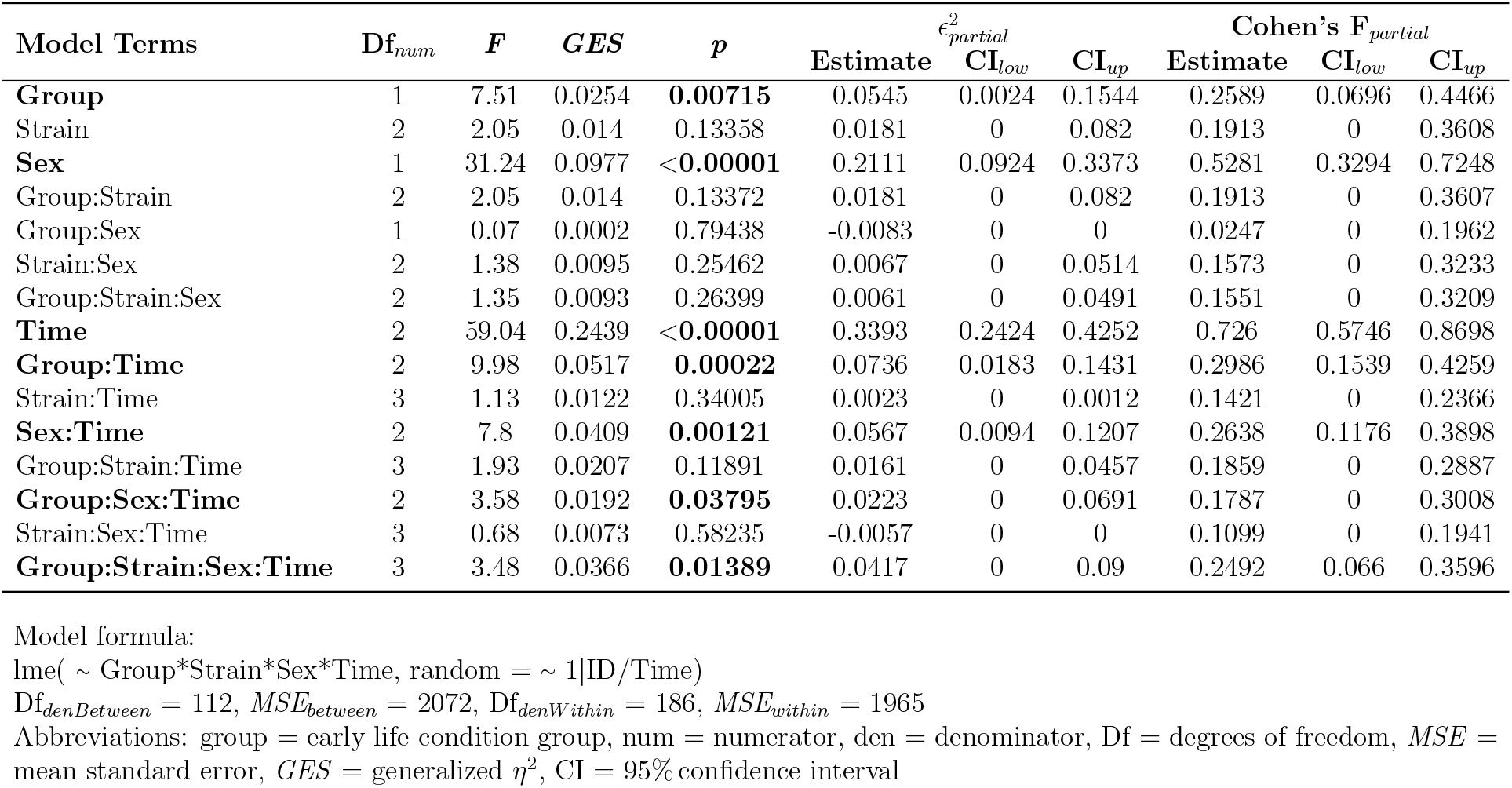
ANOVA Diurnal Corticosterone.

**Supplementary Table S4:**
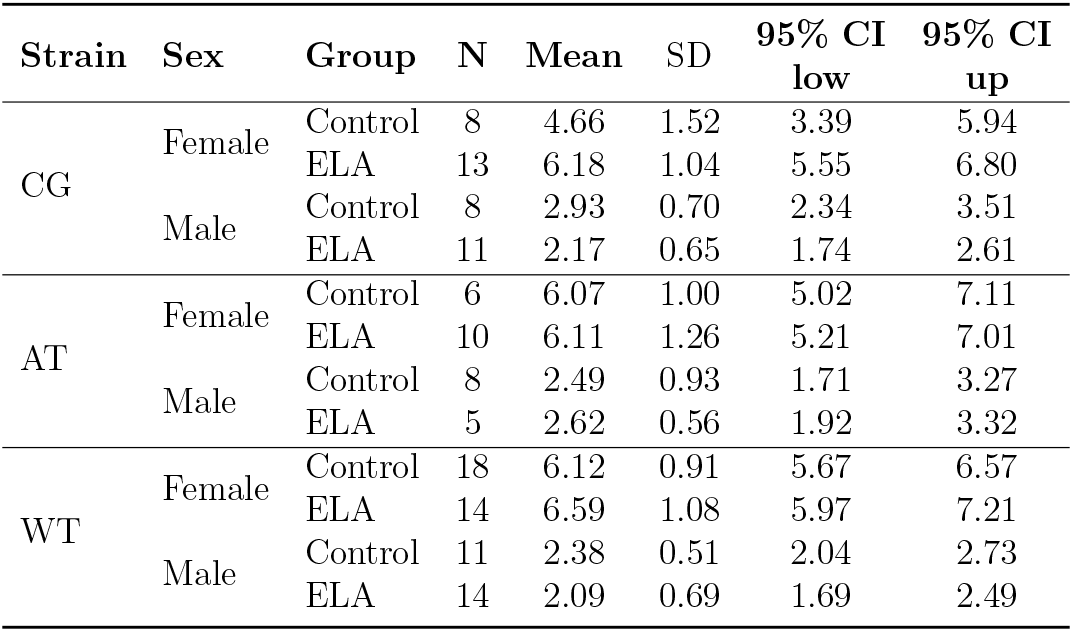
Descriptive Adrenal Weight [mg].

| Strain | Sex | Group | N | Mean | SD | 95% CI<br>low | 95% CI<br>up |
| --- | --- | --- | --- | --- | --- | --- | --- |
| CG | Female | Control | 8 | 4.66 | 1.52 | 3.39 | 5.94 |
|  |  | ELA | 13 | 6.18 | 1.04 | 5.55 | 6.80 |
|  | Male | Control | 8 | 2.93 | 0.70 | 2.34 | 3.51 |
|  |  | ELA | 11 | 2.17 | 0.65 | 1.74 | 2.61 |
| AT | Female | Control | 6 | 6.07 | 1.00 | 5.02 | 7.11 |
|  |  | ELA | 10 | 6.11 | 1.26 | 5.21 | 7.01 |
|  | Male | Control | 8 | 2.49 | 0.93 | 1.71 | 3.27 |
|  |  | ELA | 5 | 2.62 | 0.56 | 1.92 | 3.32 |
| WT | Female | Control | 18 | 6.12 | 0.91 | 5.67 | 6.57 |
|  |  | ELA | 14 | 6.59 | 1.08 | 5.97 | 7.21 |
|  | Male | Control | 11 | 2.38 | 0.51 | 2.04 | 2.73 |
|  |  | ELA | 14 | 2.09 | 0.69 | 1.69 | 2.49 |

**Supplementary Table S5:**
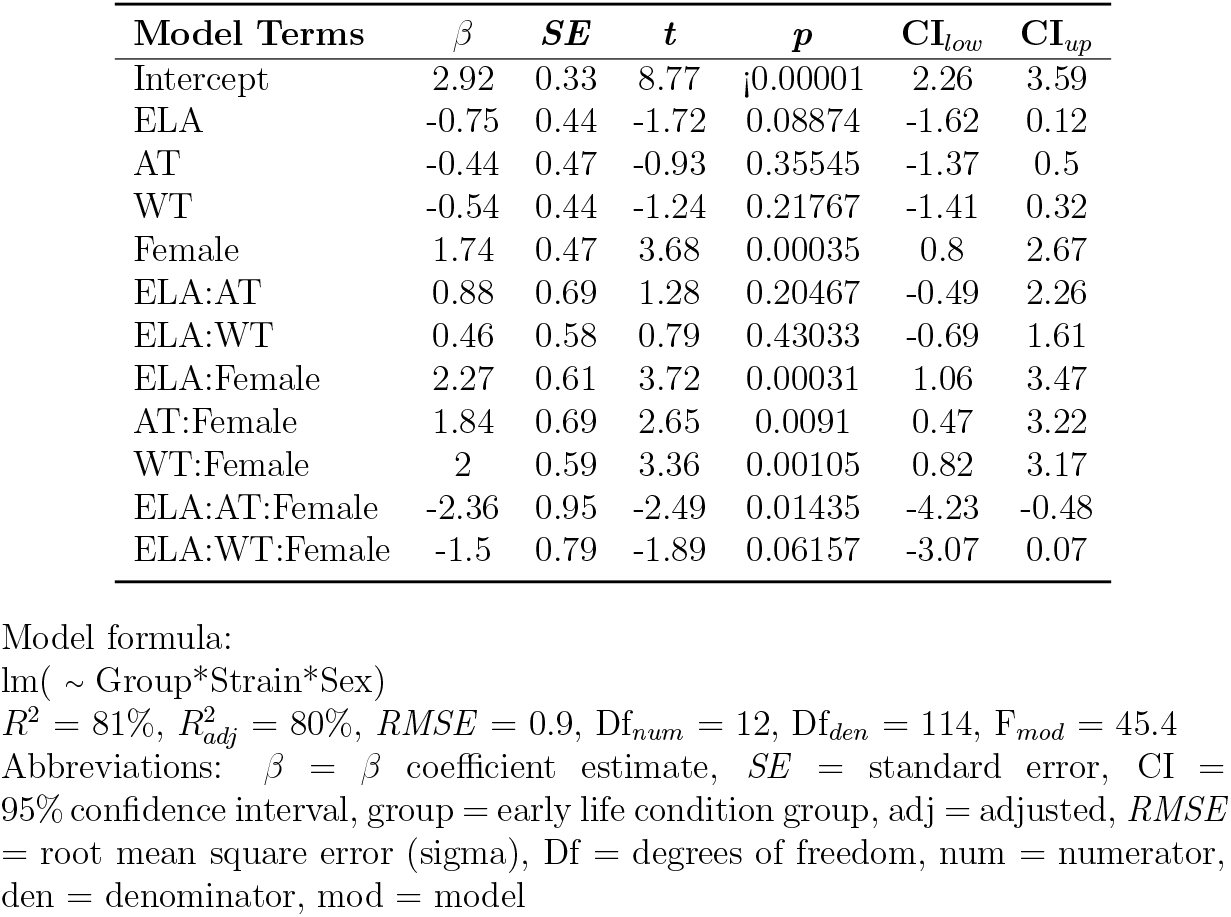
Model Summary Adrenal Weight.

**Supplementary Table S6:**
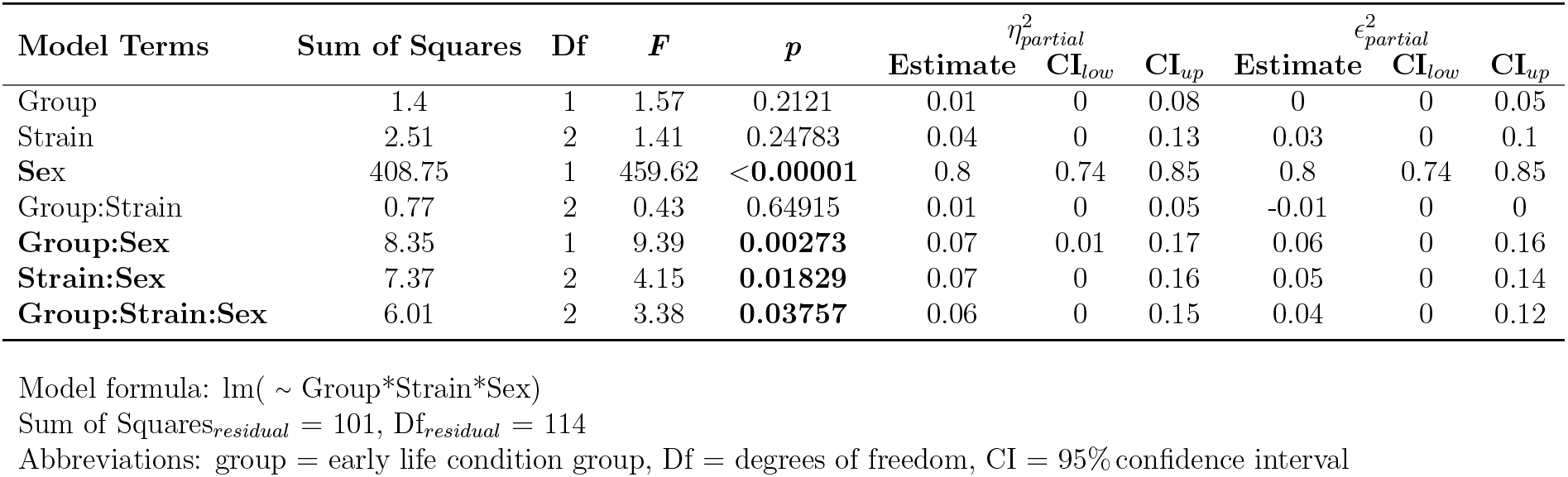
ANOVA Adrenal Weight.

**Supplementary Table S7:** Descriptive Beambreaks Novel Environment *Fkbp5*-Humanized and Wild Type Mice.

| Strain | Sex | Group | Minute | N | Mean | SD | 95% CI<br>low | 95% CI<br>up |
| --- | --- | --- | --- | --- | --- | --- | --- | --- |
| CG | Female | Control | 1 | 7 | 930.43 | 165.37 | 777.49 | 1083.37 |
|  |  |  | 2 | 7 | 883.71 | 113.22 | 779.00 | 988.43 |
|  |  |  | 5 | 7 | 740.57 | 129.63 | 620.68 | 860.46 |
|  |  |  | 10 | 7 | 680.57 | 159.12 | 533.41 | 827.74 |
|  |  |  | 15 | 7 | 640.00 | 135.22 | 514.94 | 765.06 |
|  |  | ELA | 1 | 11 | 933.73 | 165.27 | 822.70 | 1044.76 |
|  |  |  | 2 | 11 | 917.55 | 133.53 | 827.84 | 1007.25 |
|  |  |  | 5 | 11 | 826.27 | 126.28 | 741.44 | 911.11 |
|  |  |  | 10 | 11 | 755.27 | 146.47 | 656.88 | 853.67 |
|  |  |  | 15 | 11 | 713.73 | 151.06 | 612.25 | 815.21 |
|  | Male | Control | 1 | 7 | 584.86 | 186.19 | 412.66 | 757.05 |
|  |  |  | 2 | 7 | 636.29 | 217.71 | 434.94 | 837.63 |
|  |  |  | 5 | 7 | 598.71 | 68.18 | 535.66 | 661.77 |
|  |  |  | 10 | 7 | 573.71 | 52.56 | 525.11 | 622.32 |
|  |  |  | 15 | 7 | 442.14 | 160.16 | 294.01 | 590.27 |
|  |  | ELA | 1 | 11 | 640.45 | 206.55 | 501.69 | 779.22 |
|  |  |  | 2 | 11 | 714.09 | 193.50 | 584.10 | 844.08 |
|  |  |  | 5 | 11 | 635.36 | 182.28 | 512.91 | 757.82 |
|  |  |  | 10 | 11 | 590.00 | 155.13 | 485.78 | 694.22 |
|  |  |  | 15 | 11 | 583.27 | 182.96 | 460.36 | 706.18 |
| AT | Female | Control | 1 | 4 | 865.50 | 175.12 | 586.85 | 1144.15 |
|  |  |  | 2 | 4 | 825.25 | 41.77 | 758.78 | 891.72 |
|  |  |  | 5 | 4 | 808.50 | 117.71 | 621.20 | 995.80 |
|  |  |  | 10 | 4 | 686.25 | 198.54 | 370.33 | 1002.17 |
|  |  |  | 15 | 4 | 707.50 | 210.20 | 373.02 | 1041.98 |
|  |  | ELA | 1 | 10 | 817.10 | 263.28 | 628.76 | 1005.44 |
|  |  |  | 2 | 10 | 870.10 | 195.41 | 730.31 | 1009.89 |
|  |  |  | 5 | 10 | 708.70 | 123.42 | 620.41 | 796.99 |
|  |  |  | 10 | 10 | 650.50 | 166.73 | 531.23 | 769.77 |
|  |  |  | 15 | 10 | 649.60 | 188.51 | 514.75 | 784.45 |
|  | Male | Control | 1 | 7 | 631.14 | 188.82 | 456.52 | 805.77 |
|  |  |  | 2 | 7 | 691.86 | 87.43 | 611.00 | 772.72 |
|  |  |  | 5 | 7 | 705.86 | 112.21 | 602.08 | 809.63 |
|  |  |  | 10 | 7 | 547.00 | 135.95 | 421.26 | 672.74 |
|  |  |  | 15 | 7 | 472.86 | 101.99 | 378.53 | 567.19 |
|  |  | ELA | 1 | 5 | 457.40 | 157.74 | 261.54 | 653.26 |
|  |  |  | 2 | 5 | 602.60 | 165.87 | 396.65 | 808.55 |
|  |  |  | 5 | 5 | 631.00 | 112.92 | 490.79 | 771.21 |
|  |  |  | 10 | 5 | 605.20 | 155.49 | 412.14 | 798.26 |
|  |  |  | 15 | 5 | 532.60 | 84.01 | 428.28 | 636.92 |
| WT | Female | Control | 1 | 20 | 840.05 | 215.85 | 739.03 | 941.07 |
|  |  |  | 2 | 20 | 835.15 | 184.61 | 748.75 | 921.55 |
|  |  |  | 5 | 20 | 780.50 | 113.82 | 727.23 | 833.77 |
|  |  |  | 10 | 20 | 667.20 | 175.43 | 585.10 | 749.30 |
|  |  |  | 15 | 20 | 564.35 | 137.49 | 500.00 | 628.70 |
|  |  | ELA | 1 | 17 | 856.18 | 243.75 | 730.85 | 981.50 |
|  |  |  | 2 | 17 | 888.71 | 173.93 | 799.28 | 978.13 |
|  |  |  | 5 | 17 | 866.12 | 160.29 | 783.70 | 948.53 |
|  |  |  | 10 | 17 | 706.12 | 127.78 | 640.42 | 771.81 |
|  |  |  | 15 | 17 | 660.12 | 105.66 | 605.79 | 714.44 |
|  | Male | Control | 1 | 15 | 613.13 | 228.94 | 486.35 | 739.92 |
|  |  |  | 2 | 15 | 574.73 | 130.75 | 502.33 | 647.14 |
|  |  |  | 5 | 15 | 602.00 | 127.92 | 531.16 | 672.84 |
|  |  |  | 10 | 15 | 563.13 | 95.31 | 510.35 | 615.91 |
|  |  |  | 15 | 15 | 436.40 | 149.18 | 353.79 | 519.01 |
|  |  | ELA | 1 | 17 | 664.59 | 171.33 | 576.50 | 752.68 |
|  |  |  | 2 | 17 | 705.24 | 182.75 | 611.27 | 799.20 |
|  |  |  | 5 | 17 | 744.65 | 91.93 | 697.38 | 791.91 |
|  |  |  | 10 | 17 | 619.71 | 142.12 | 546.63 | 692.78 |
|  |  |  | 15 | 17 | 596.47 | 92.69 | 548.82 | 644.13 |

**Supplementary Table S8:** Model Summary Beambreaks Habituation OFT.

| **Model Terms** | $\beta$ | ***SE*** | **Df** | **CI<sub>low</sub>** | **CI<sub>up</sub>** | ***t*** | ***p*** | ***R*<sub>part</sub><sup>2</sup>** |
| --- | --- | --- | --- | --- | --- | --- | --- | --- |
| Intercept | 887.94 | 39.72 | 1822 | 810.04 | 965.84 | 22.35 | 0 | 0.4008 |
| ELA | 46 | 50.81 | 119 | -54.61 | 146.61 | 0.91 | 0.3671 | 0.0234 |
| AT | -41.87 | 65.87 | 119 | -172.29 | 88.56 | -0.64 | 0.5263 | 0.021 |
| WT | -5.27 | 46.15 | 119 | -96.66 | 86.11 | -0.11 | 0.9093 | 0.0032 |
| Male | -231.67 | 56.17 | 119 | -342.9 | -120.44 | -4.12 | 0.0001 | 0.0023 |
| Min | -19.04 | 2.68 | 1822 | -24.29 | -13.78 | -7.1 | 0 | 0.0011 |
| ELA:AT | -73.48 | 80.29 | 119 | -232.47 | 85.52 | -0.92 | 0.362 | 0.0011 |
| ELA:WT | -31.76 | 61.51 | 119 | -153.56 | 90.04 | -0.52 | 0.6066 | 0.0011 |
| ELA:Male | -2.86 | 71.86 | 119 | -145.14 | 139.43 | -0.04 | 0.9684 | 0.001 |
| AT:Male | 117.84 | 86.57 | 119 | -53.58 | 289.25 | 1.36 | 0.176 | 0.001 |
| WT:Male | 3.67 | 66.66 | 119 | -128.33 | 135.67 | 0.06 | 0.9562 | 0.0008 |
| ELA:Min | 4.49 | 3.43 | 1822 | -2.24 | 11.21 | 1.31 | 0.1908 | 0.0007 |
| AT:Min | 4.44 | 4.44 | 1822 | -4.28 | 13.15 | 1 | 0.3183 | 0.0005 |
| WT:Min | -2.08 | 3.11 | 1822 | -8.19 | 4.03 | -0.67 | 0.5039 | 0.0005 |
| Male:Min | 9.85 | 3.79 | 1822 | 2.41 | 17.28 | 2.6 | 0.0095 | 0.0003 |
| ELA:AT:Male | -86.08 | 113.21 | 119 | -310.24 | 138.08 | -0.76 | 0.4485 | 0.0002 |
| ELA:WT:Male | 77.84 | 88.04 | 119 | -96.49 | 252.18 | 0.88 | 0.3784 | 0.0001 |
| ELA:AT:Min | -2.33 | 5.42 | 1822 | -12.96 | 8.3 | -0.43 | 0.667 | 0.0001 |
| ELA:WT:Min | 1.11 | 4.15 | 1822 | -7.03 | 9.25 | 0.27 | 0.7895 | 0.0001 |
| ELA:Male:Min | -2.55 | 4.85 | 1822 | -12.06 | 6.96 | -0.53 | 0.5988 | 0 |
| AT:Male:Min | -8.99 | 5.84 | 1822 | -20.45 | 2.46 | -1.54 | 0.1239 | 0 |
| WT:Male:Min | -1.52 | 4.5 | 1822 | -10.34 | 7.31 | -0.34 | 0.7363 | 0 |
| ELA:AT:Male:Min | 11.74 | 7.64 | 1822 | -3.25 | 26.72 | 1.54 | 0.1246 | 0 |
| ELA:WT:Male:Min | -1.11 | 5.94 | 1822 | -12.76 | 10.55 | -0.19 | 0.8524 | 0 |

Model formula:
`lme( ~ Group*Strain*Sex*Minute, random = ~ 1|ID/Minute)`
$R^2_{marg} = 32.54\%$ , $R^2_{cond} = 93.57\%$
Abbreviations: $\beta$ = $\beta$ coefficient estimate, *SE* = standard error, Df = degrees of freedom, CI = 95% confidence interval, part = partial, group = early life condition group, Min = minute, marg = marginalized, cond = conditioned,

**Supplementary Table S9:**
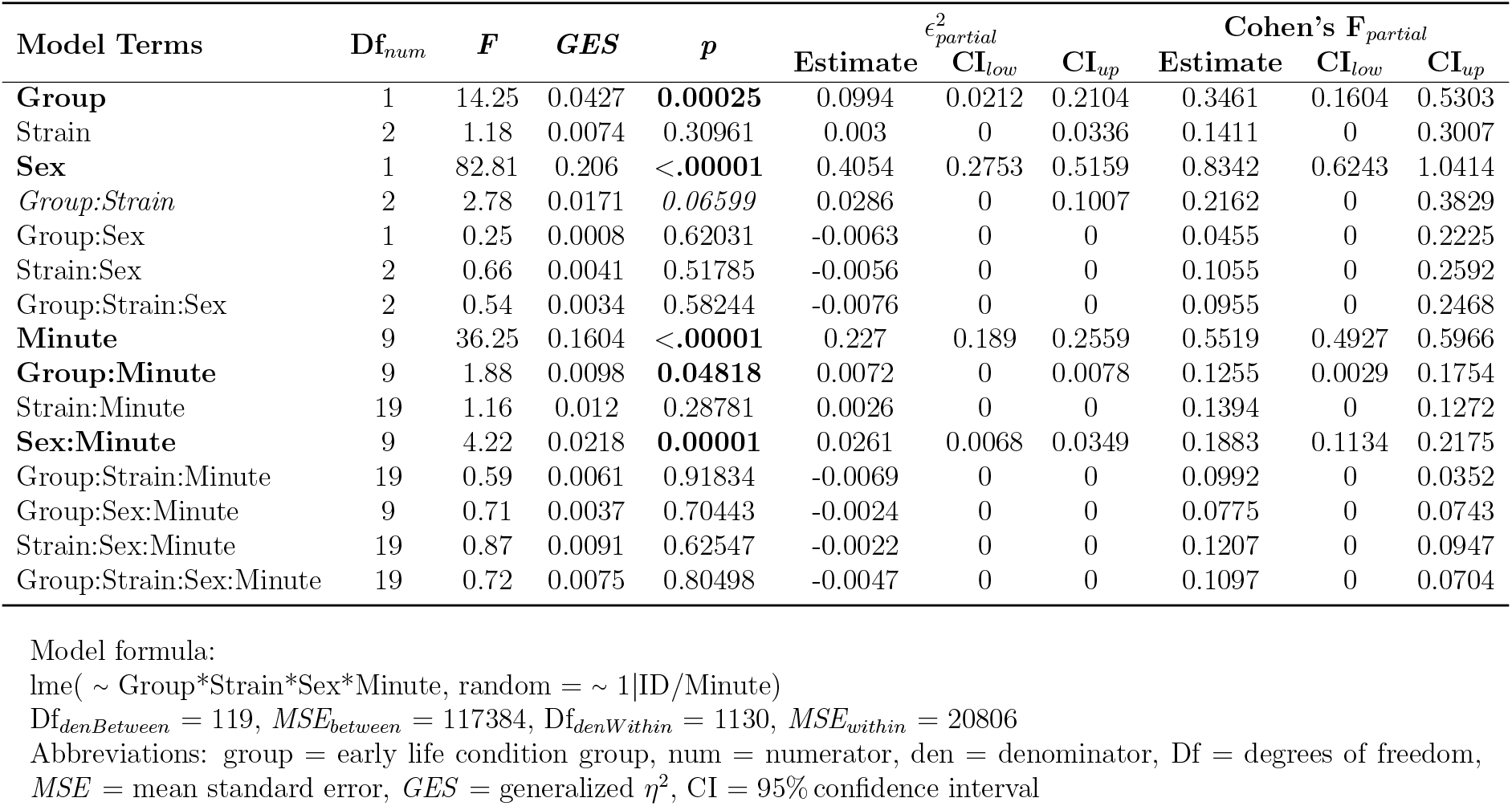
ANOVA Beambreaks Habituation OFT.

**Supplementary Table S10:** Descriptive Nocturnal Distance [m].

| Strain | Sex | Group | N | Mean | SD | 95% CI<br>low | 95% CI<br>up |
| --- | --- | --- | --- | --- | --- | --- | --- |
| CG | Female | Control | 7 | 2586.16 | 1117.10 | 1553.01 | 3619.30 |
|  |  | ELA | 11 | 4218.35 | 899.67 | 3613.95 | 4822.76 |
|  | Male | Control | 7 | 2259.04 | 353.83 | 1931.80 | 2586.28 |
|  |  | ELA | 11 | 2700.24 | 793.25 | 2167.33 | 3233.15 |
| AT | Female | Control | 4 | 3264.80 | 610.13 | 2293.94 | 4235.66 |
|  |  | ELA | 10 | 3044.38 | 864.49 | 2425.96 | 3662.80 |
|  | Male | Control | 7 | 2626.46 | 617.70 | 2055.18 | 3197.73 |
|  |  | ELA | 5 | 2455.84 | 376.49 | 1988.37 | 2923.31 |
| WT | Female | Control | 20 | 3199.28 | 948.47 | 2755.38 | 3643.17 |
|  |  | ELA | 17 | 3570.82 | 967.43 | 3073.41 | 4068.23 |
|  | Male | Control | 15 | 2466.61 | 662.09 | 2099.95 | 2833.26 |
|  |  | ELA | 17 | 2579.88 | 470.53 | 2337.96 | 2821.81 |

**Supplementary Table S11:** Model Summary Nocturnal Distance OFT.

| Model Terms | $\beta$ | $SE$ | $t$ | $p$ | $CI_{low}$ | $CI_{up}$ |
| --- | --- | --- | --- | --- | --- | --- |
| Intercept | 2586.16 | 300.2 | 8.61 | <0.00001 | 1991.73 | 3180.59 |
| ELA | 1632.2 | 384.02 | 4.25 | 0.00004 | 871.8 | 2392.59 |
| AT | 678.64 | 497.83 | 1.36 | 0.17539 | -307.11 | 1664.39 |
| WT | 613.12 | 348.8 | 1.76 | 0.08136 | -77.55 | 1303.78 |
| Male | -327.11 | 424.55 | -0.77 | 0.44253 | -1167.76 | 513.54 |
| ELA:AT | -1852.62 | 606.85 | -3.05 | 0.0028 | -3054.24 | -650.99 |
| ELA:WT | -1260.65 | 464.89 | -2.71 | 0.00768 | -2181.18 | -340.13 |
| ELA:Male | -1191 | 543.09 | -2.19 | 0.03025 | -2266.37 | -115.64 |
| AT:Male | -311.23 | 654.27 | -0.48 | 0.63517 | -1606.76 | 984.3 |
| WT:Male | -405.55 | 503.83 | -0.8 | 0.42246 | -1403.18 | 592.07 |
| ELA:AT:Male | 1240.81 | 855.59 | 1.45 | 0.14962 | -453.34 | 2934.95 |
| ELA:WT:Male | 932.74 | 665.4 | 1.4 | 0.16359 | -384.82 | 2250.3 |

Model formula:
$\text{lm}(\sim \text{Group} * \text{Strain} * \text{Sex})$
$R^2 = 34\%$ , $R^2_{adj} = 28\%$ , $RMSE = 794$ , $Df_{num} = 12$ , $Df_{den} = 119$ , $F_{mod} = 5.6$
Abbreviations: $\beta$ = $\beta$ coefficient estimate, $SE$ = standard error, $CI$ = 95% confidence interval, group = early life condition group, adj = adjusted, $RMSE$ = root mean square error (sigma), $Df$ = degrees of freedom, num = numerator, den = denominator, mod = model

**Supplementary Table S12:** ANOVA Nocturnal Distance OFT.

| Model Terms | Sum of Squares | Df | *F* | *p* | $\eta^2_{partial}$ | | | $\epsilon^2_{partial}$ | | |
| --- | --- | --- | --- | --- | --- | --- | --- | --- | --- | --- |
|  |  |  |  |  | Estimate | CI<sub>low</sub> | CI<sub>up</sub> | Estimate | CI<sub>low</sub> | CI<sub>up</sub> |
| **Group** | 4618308.48 | 1 | 7.32 | **0.00782** | 0.06 | 0 | 0.15 | 0.05 | 0 | 0.14 |
| Strain | 1068955.1 | 2 | 0.85 | 0.43117 | 0.01 | 0 | 0.05 | -0.01 | 0 | 0 |
| **Sex** | 24161371.46 | 1 | 38.3 | **<0.00001** | 0.23 | 0.11 | 0.36 | 0.23 | 0.11 | 0.35 |
| **Group:Strain** | 5942021.42 | 2 | 4.71 | **0.01075** | 0.08 | 0.01 | 0.18 | 0.07 | 0 | 0.16 |
| Group:Sex | 1630636.87 | 1 | 2.58 | 0.11054 | 0.02 | 0 | 0.1 | 0.01 | 0 | 0.08 |
| Strain:Sex | 646520.39 | 2 | 0.51 | 0.60036 | 0.01 | 0 | 0.06 | -0.01 | 0 | 0 |
| Group:Strain:Sex | 1691604.95 | 2 | 1.34 | 0.26558 | 0.02 | 0 | 0.09 | 0.01 | 0 | 0.05 |

Model formula: lm( ~ Group\*Strain\*Sex)
Sum of Squares<sub>residual</sub> = 8<sup>7</sup>, Df<sub>residual</sub> = 119
Abbreviations: group = early life condition group, Df = degrees of freedom, CI = 95% confidence interval

**Supplementary Table S13:** Descriptive Alternations [%] T-Maze.

| Read Out | Strain | Sex | Group | N | Mean | SD | 95% CI<br>low | 95% CI<br>up |
| --- | --- | --- | --- | --- | --- | --- | --- | --- |
| Alternations | CG | Female | Control | 7 | 57.14 | 19.34 | 39.25 | 75.03 |
|  |  |  | ELA | 11 | 61.04 | 18.45 | 48.64 | 73.43 |
|  |  | Male | Control | 7 | 55.10 | 27.27 | 29.89 | 80.32 |
|  |  |  | ELA | 11 | 56.49 | 11.27 | 48.92 | 64.07 |
|  | AT | Female | Control | 5 | 61.43 | 12.98 | 45.32 | 77.54 |
|  |  |  | ELA | 10 | 46.43 | 10.24 | 39.10 | 53.75 |
|  |  | Male | Control | 7 | 58.16 | 8.68 | 50.14 | 66.19 |
|  |  |  | ELA | 5 | 48.57 | 7.82 | 38.86 | 58.29 |
|  | WT | Female | Control | 20 | 51.07 | 21.40 | 41.06 | 61.09 |
|  |  |  | ELA | 17 | 55.04 | 15.92 | 46.85 | 63.23 |
|  |  | Male | Control | 15 | 55.71 | 18.35 | 45.55 | 65.88 |
|  |  |  | ELA | 17 | 60.50 | 17.33 | 51.59 | 69.42 |

**Supplementary Table S14:** Model Summary Alternations T-Maze.

| Model Terms | $\beta$ | *SE* | *t* | *p* | CI<sub>low</sub> | CI<sub>up</sub> |
| --- | --- | --- | --- | --- | --- | --- |
| Intercept | 57.14 | 6.54 | 8.73 | <0.00001 | 44.19 | 70.1 |
| ELA | 3.9 | 8.37 | 0.47 | 0.64249 | -12.68 | 20.47 |
| AT | 4.29 | 10.14 | 0.42 | 0.67326 | -15.79 | 24.36 |
| WT | -6.07 | 7.6 | -0.8 | 0.42618 | -21.13 | 8.98 |
| Male | -2.04 | 9.26 | -0.22 | 0.82585 | -20.37 | 16.28 |
| ELA:AT | -18.9 | 12.65 | -1.49 | 0.13787 | -43.94 | 6.15 |
| ELA:WT | 0.07 | 10.13 | 0.01 | 0.99415 | -19.99 | 20.14 |
| ELA:Male | -2.5 | 11.84 | -0.21 | 0.83281 | -25.95 | 20.94 |
| AT:Male | -1.22 | 13.73 | -0.09 | 0.92907 | -28.4 | 25.96 |
| WT:Male | 6.68 | 10.98 | 0.61 | 0.54399 | -15.06 | 28.43 |
| ELA:AT:Male | 7.91 | 18.25 | 0.43 | 0.6653 | -28.21 | 44.04 |
| ELA:WT:Male | 3.32 | 14.51 | 0.23 | 0.81914 | -25.4 | 32.04 |

Model formula:
lm( ~ Group\*Strain\*Sex)
$R^2 = 6\%$ , $R^2_{adj} = -2\%$ , $RMSE = 17.3$ , $Df_{num} = 12$ , $Df_{den} = 120$ , $F_{mod} = 0.75$
Abbreviations: $\beta$ = $\beta$ coefficient estimate, *SE* = standard error, CI = 95% confidence interval, group = early life condition group, adj = adjusted, *RMSE* = root mean square error (sigma), Df = degrees of freedom, num = numerator, den = denominator, mod = model

**Supplementary Table S15:**
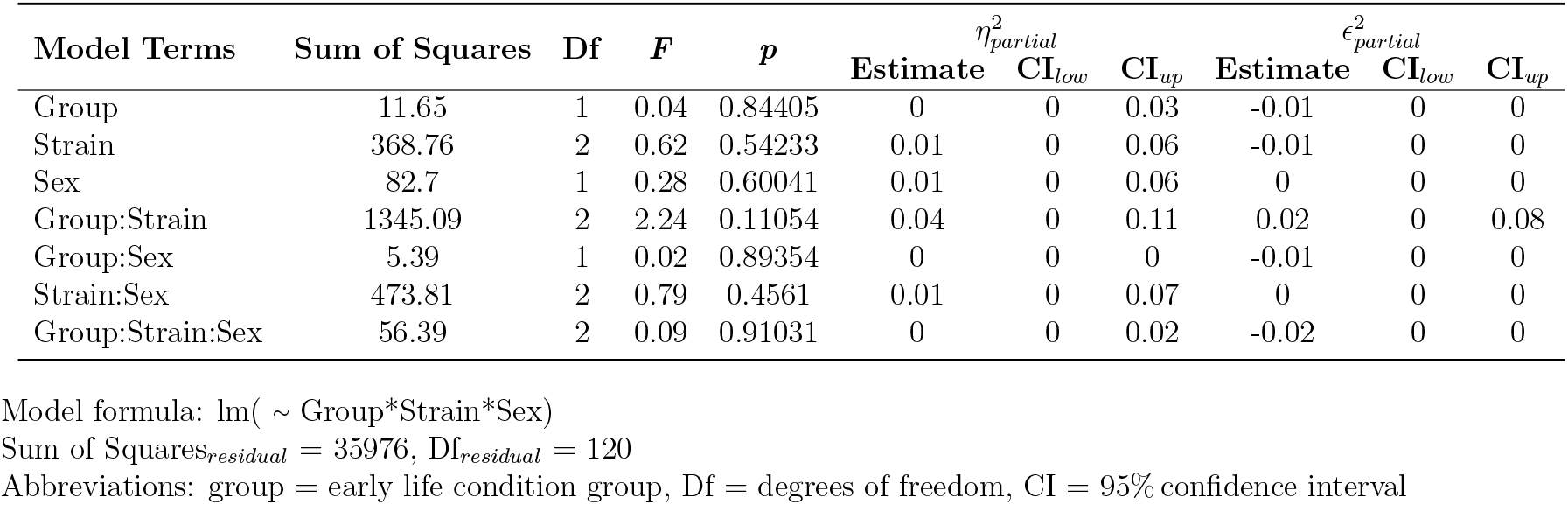
ANOVA T-Maze Alternations.

| Model Terms | Sum of Squares | Df | *F* | *p* | $\eta^2_{partial}$ | | | $\epsilon^2_{partial}$ | | |
| --- | --- | --- | --- | --- | --- | --- | --- | --- | --- | --- |
|  |  |  |  |  | Estimate | CI<sub>low</sub> | CI<sub>up</sub> | Estimate | CI<sub>low</sub> | CI<sub>up</sub> |
| Group | 11.65 | 1 | 0.04 | 0.84405 | 0 | 0 | 0.03 | -0.01 | 0 | 0 |
| Strain | 368.76 | 2 | 0.62 | 0.54233 | 0.01 | 0 | 0.06 | -0.01 | 0 | 0 |
| Sex | 82.7 | 1 | 0.28 | 0.60041 | 0.01 | 0 | 0.06 | 0 | 0 | 0 |
| Group:Strain | 1345.09 | 2 | 2.24 | 0.11054 | 0.04 | 0 | 0.11 | 0.02 | 0 | 0.08 |
| Group:Sex | 5.39 | 1 | 0.02 | 0.89354 | 0 | 0 | 0 | -0.01 | 0 | 0 |
| Strain:Sex | 473.81 | 2 | 0.79 | 0.4561 | 0.01 | 0 | 0.07 | 0 | 0 | 0 |
| Group:Strain:Sex | 56.39 | 2 | 0.09 | 0.91031 | 0 | 0 | 0.02 | -0.02 | 0 | 0 |

Model formula: $\text{lm}(\sim \text{Group} * \text{Strain} * \text{Sex})$
Sum of Squares<sub>residual</sub> = 35976, Df<sub>residual</sub> = 120
Abbreviations: group = early life condition group, Df = degrees of freedom, CI = 95% confidence interval

**Supplementary Table S16:** Descriptive Latency to Completion of the T-Maze.

| Read Out | Strain | Sex | Group | N | Mean | SD | 95% CI<br>low | 95% CI<br>up |
| --- | --- | --- | --- | --- | --- | --- | --- | --- |
| Latency | CG | Female | Control | 7 | 8.71 | 3.50 | 5.48 | 11.95 |
|  |  |  | ELA | 11 | 5.55 | 1.81 | 4.33 | 6.76 |
|  |  | Male | Control | 6 | 10.83 | 3.13 | 7.55 | 14.11 |
|  |  |  | ELA | 11 | 7.82 | 1.25 | 6.98 | 8.66 |
|  | AT | Female | Control | 5 | 7.60 | 1.67 | 5.52 | 9.68 |
|  |  |  | ELA | 10 | 6 | 2.21 | 4.42 | 7.58 |
|  |  | Male | Control | 7 | 9.57 | 2.30 | 7.45 | 11.70 |
|  |  |  | ELA | 5 | 8.20 | 3.77 | 3.52 | 12.88 |
|  | WT | Female | Control | 18 | 8.72 | 3.18 | 7.14 | 10.30 |
|  |  |  | ELA | 17 | 6.24 | 1.44 | 5.50 | 6.97 |
|  |  | Male | Control | 14 | 9.50 | 2.24 | 8.20 | 10.80 |
|  |  |  | ELA | 17 | 8.94 | 2.22 | 7.80 | 10.08 |

**Supplementary Table S17:** Model Summary Latency Completion T-maze.

| Model Terms | $\beta$ | *SE* | *t* | *p* | CI<sub>low</sub> | CI<sub>up</sub> |
| --- | --- | --- | --- | --- | --- | --- |
| Intercept | 8.71 | 0.9 | 9.64 | <0.00001 | 6.92 | 10.5 |
| ELA | -3.17 | 1.16 | -2.74 | 0.00711 | -5.46 | -0.88 |
| AT | -1.11 | 1.4 | -0.8 | 0.42782 | -3.89 | 1.66 |
| WT | 0.01 | 1.07 | 0.01 | 0.99407 | -2.1 | 2.12 |
| Male | 2.12 | 1.33 | 1.59 | 0.11396 | -0.52 | 4.75 |
| ELA:AT | 1.57 | 1.75 | 0.9 | 0.3711 | -1.89 | 5.03 |
| ELA:WT | 0.68 | 1.41 | 0.48 | 0.62983 | -2.11 | 3.48 |
| ELA:Male | 0.15 | 1.68 | 0.09 | 0.92711 | -3.17 | 3.47 |
| AT:Male | -0.15 | 1.93 | -0.08 | 0.93922 | -3.97 | 3.68 |
| WT:Male | -1.34 | 1.58 | -0.85 | 0.3977 | -4.47 | 1.79 |
| ELA:AT:Male | 0.07 | 2.55 | 0.03 | 0.97659 | -4.97 | 5.12 |
| ELA:WT:Male | 1.77 | 2.05 | 0.86 | 0.38889 | -2.29 | 5.84 |

Model formula:
lm( ~ Group\*Strain\*Sex)
$R^2 = 30\%$ , $R^2_{adj} = 23\%$ , $RMSE = 2.4$ , $Df_{num} = 12$ , $Df_{den} = 116$ , $F_{mod} = 4.5$
Abbreviations: $\beta$ = $\beta$ coefficient estimate, *SE* = standard error, CI = 95% confidence interval, group = early life condition group, adj = adjusted, *RMSE* = root mean square error (sigma), Df = degrees of freedom, num = numerator, den = denominator, mod = model

**Supplementary Table S18:**
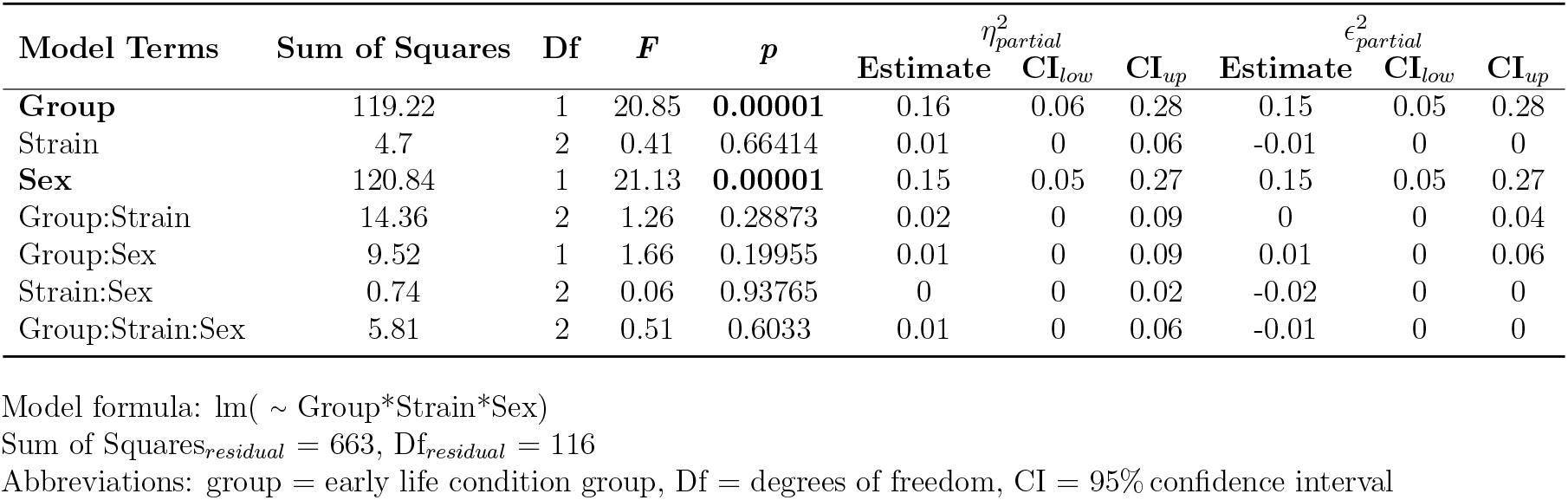
ANOVA T-Maze Latency Completion.

**Supplementary Table S19:** Descriptive Average Time [s/min] in the Dark Compartment.

| Strain | Sex | Group | N | Mean | SD | 95% CI<br>low | 95% CI<br>up |
| --- | --- | --- | --- | --- | --- | --- | --- |
| CG | Female | Control | 7 | 40.29 | 5.75 | 34.97 | 45.60 |
|  |  | ELA | 11 | 31.15 | 10.12 | 24.36 | 37.95 |
|  | Male | Control | 7 | 41.73 | 4.80 | 37.29 | 46.17 |
|  |  | ELA | 11 | 32.68 | 9.66 | 26.19 | 39.17 |
| AT | Female | Control | 5 | 40.70 | 4.85 | 34.68 | 46.72 |
|  |  | ELA | 10 | 38.93 | 9.71 | 31.98 | 45.88 |
|  | Male | Control | 7 | 42.24 | 6.66 | 36.09 | 48.40 |
|  |  | ELA | 5 | 41.42 | 2.75 | 38.00 | 44.84 |
| WT | Female | Control | 20 | 41.29 | 6.42 | 38.29 | 44.29 |
|  |  | ELA | 17 | 34.96 | 11.24 | 29.19 | 40.74 |
|  | Male | Control | 15 | 42.99 | 6.50 | 39.38 | 46.59 |
|  |  | ELA | 17 | 33.16 | 7.73 | 29.19 | 37.14 |

**Supplementary Table S20:**
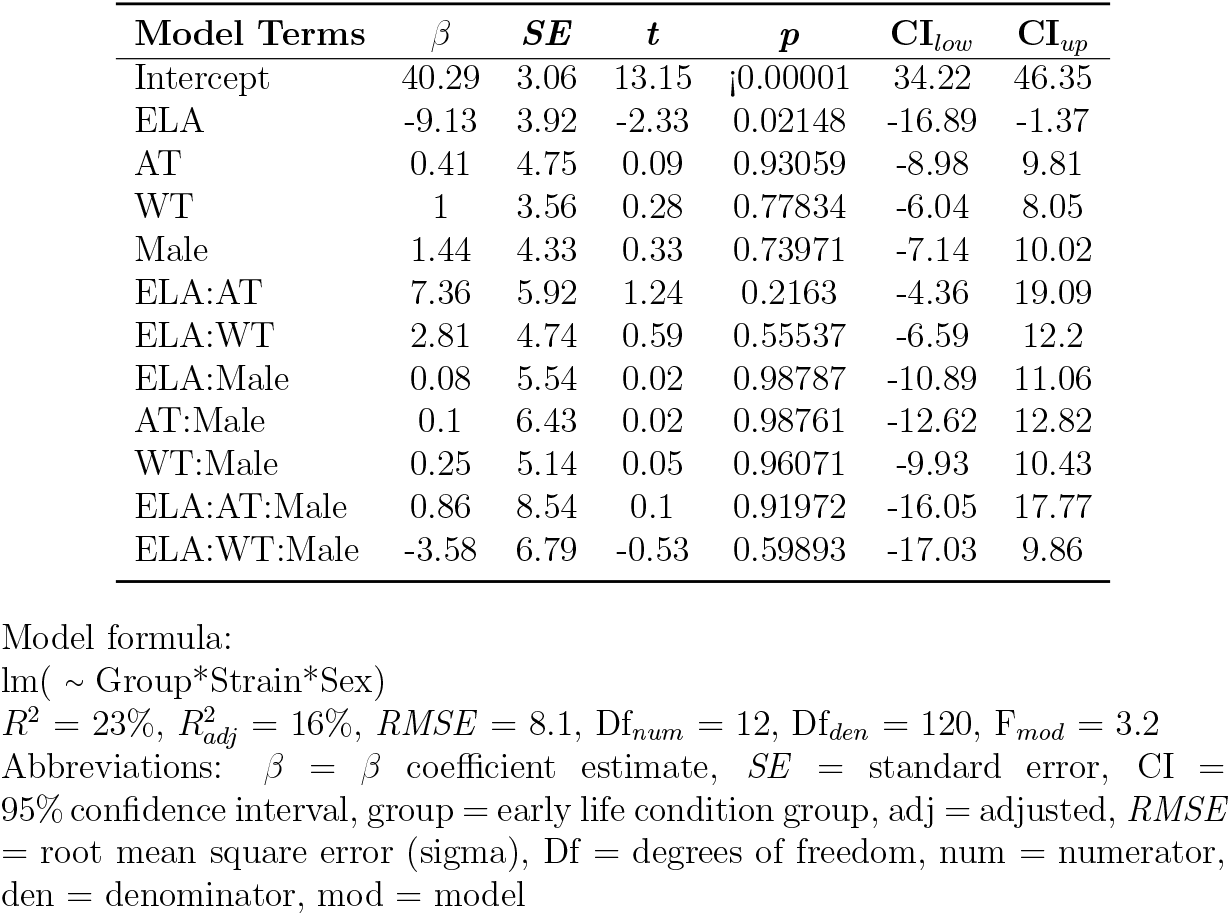
Model Summary Dark-Light-Test.

**Supplementary Table S21:** ANOVA Time in the Dark Compartment during Dark-Light Test.

| Model Terms | Sum of Squares | Df | *F* | *p* | $\eta_{partial}^2$ | | | $\epsilon_{partial}^2$ | | |
| --- | --- | --- | --- | --- | --- | --- | --- | --- | --- | --- |
|  |  |  |  |  | Estimate | CI<sub>low</sub> | CI<sub>up</sub> | Estimate | CI<sub>low</sub> | CI<sub>up</sub> |
| **Group** | 1548.2 | 1 | 23.56 | <0.00001 | 0.17 | 0.07 | 0.29 | 0.17 | 0.06 | 0.29 |
| *Strain* | 332.12 | 2 | 2.53 | 0.08412 | 0.04 | 0 | 0.12 | 0.03 | 0 | 0.1 |
| Sex | 20.11 | 1 | 0.31 | 0.58117 | 0 | 0 | 0.04 | -0.01 | 0 | 0 |
| Group:Strain | 251.9 | 2 | 1.92 | 0.15154 | 0.03 | 0 | 0.1 | 0.01 | 0 | 0.07 |
| Group:Sex | 22.11 | 1 | 0.34 | 0.56292 | 0 | 0 | 0.05 | -0.01 | 0 | 0 |
| Strain:Sex | 29.62 | 2 | 0.23 | 0.79854 | 0 | 0 | 0.04 | -0.01 | 0 | 0 |
| Group:Strain:Sex | 31.48 | 2 | 0.24 | 0.78737 | 0 | 0 | 0.04 | -0.01 | 0 | 0 |

Model formula: lm( ~ Group\*Strain\*Sex)
Sum of Squares<sub>residual</sub> = 7885, Df<sub>residual</sub> = 120
Abbreviations: group = early life condition group, Df = degrees of freedom, CI = 95% confidence interval

**Supplementary Table S22:** Descriptive Social Interaction [% time].

| Strain | Sex | Group | Compartment | N | Mean | SD | 95% CI<br>low | 95% CI<br>up |
| --- | --- | --- | --- | --- | --- | --- | --- | --- |
| CG | Female | Control | Reference | 7 | 3.343 | 1.767 | 1.709 | 4.977 |
|  |  |  | Social | 7 | 11.071 | 1.442 | 9.738 | 12.405 |
|  |  | ELA | Reference | 11 | 3.936 | 6.307 | -0.301 | 8.173 |
|  |  |  | Social | 11 | 5.864 | 4.883 | 2.583 | 9.144 |
|  | Male | Control | Reference | 7 | 1.957 | 1.811 | 0.282 | 3.632 |
|  |  |  | Social | 7 | 10.986 | 6.998 | 4.513 | 17.458 |
|  |  | ELA | Reference | 11 | 3.000 | 1.872 | 1.742 | 4.258 |
|  |  |  | Social | 11 | 10.100 | 6.727 | 5.581 | 14.619 |
| AT | Female | Control | Reference | 5 | 1.960 | 1.260 | 0.395 | 3.525 |
|  |  |  | Social | 5 | 6.120 | 1.840 | 3.835 | 8.405 |
|  |  | ELA | Reference | 10 | 5.070 | 7.379 | -0.209 | 10.349 |
|  |  |  | Social | 10 | 7.680 | 4.878 | 4.190 | 11.170 |
|  | Male | Control | Reference | 7 | 2.643 | 1.512 | 1.244 | 4.041 |
|  |  |  | Social | 7 | 12.771 | 3.582 | 9.459 | 16.084 |
|  |  | ELA | Reference | 5 | 1.380 | 1.341 | -0.284 | 3.044 |
|  |  |  | Social | 5 | 6.000 | 3.694 | 1.413 | 10.587 |
| WT | Female | Control | Reference | 16 | 4.088 | 3.509 | 2.218 | 5.957 |
|  |  |  | Social | 16 | 8.144 | 5.467 | 5.231 | 11.057 |
|  |  | ELA | Reference | 16 | 3.269 | 2.256 | 2.066 | 4.471 |
|  |  |  | Social | 16 | 8.781 | 5.098 | 6.065 | 11.498 |
|  | Male | Control | Reference | 11 | 3.782 | 2.020 | 2.425 | 5.139 |
|  |  |  | Social | 11 | 10.927 | 4.814 | 7.693 | 14.161 |
|  |  | ELA | Reference | 14 | 3.129 | 1.715 | 2.138 | 4.119 |
|  |  |  | Social | 14 | 9.871 | 4.787 | 7.107 | 12.635 |

**Supplementary Table S23:** Model Summary Interaction Time SCT.

| **Model Terms** | $\beta$ | ***SE*** | **Df** | **CI<sub>low</sub>** | **CI<sub>up</sub>** | ***t*** | ***p*** | **$R^2_{part}$** |
| --- | --- | --- | --- | --- | --- | --- | --- | --- |
| Intercept | 3.34 | 1.64 | 108 | 0.1 | 6.59 | 2.04 | 0.0436 | 0.3595 |
| ELA | 0.59 | 2.09 | 108 | -3.56 | 4.74 | 0.28 | 0.7774 | 0.0451 |
| AT | -1.38 | 2.54 | 108 | -6.41 | 3.64 | -0.55 | 0.5867 | 0.0163 |
| WT | 0.74 | 1.96 | 108 | -3.15 | 4.64 | 0.38 | 0.7052 | 0.016 |
| Male | -1.39 | 2.32 | 108 | -5.98 | 3.2 | -0.6 | 0.5508 | 0.0074 |
| Social | 7.73 | 2.32 | 108 | 3.14 | 12.32 | 3.34 | 0.0012 | 0.0062 |
| ELA:AT | 2.52 | 3.16 | 108 | -3.76 | 8.79 | 0.8 | 0.4283 | 0.0049 |
| ELA:WT | -1.41 | 2.59 | 108 | -6.56 | 3.73 | -0.54 | 0.5873 | 0.0047 |
| ELA:Male | 0.45 | 2.96 | 108 | -5.42 | 6.32 | 0.15 | 0.8797 | 0.0042 |
| AT:Male | 2.07 | 3.43 | 108 | -4.74 | 8.88 | 0.6 | 0.5482 | 0.0039 |
| WT:Male | 1.08 | 2.87 | 108 | -4.61 | 6.77 | 0.38 | 0.7075 | 0.0038 |
| ELA:Social | -5.8 | 2.96 | 108 | -11.67 | 0.07 | -1.96 | 0.0527 | 0.0036 |
| AT:Social | -3.57 | 3.59 | 108 | -10.68 | 3.54 | -0.99 | 0.322 | 0.0027 |
| WT:Social | -3.67 | 2.78 | 108 | -9.17 | 1.83 | -1.32 | 0.1887 | 0.0015 |
| Male:Social | 1.3 | 3.27 | 108 | -5.19 | 7.79 | 0.4 | 0.6921 | 0.0015 |
| ELA:AT:Male | -4.82 | 4.56 | 108 | -13.87 | 4.23 | -1.06 | 0.2931 | 0.0013 |
| ELA:WT:Male | -0.28 | 3.76 | 108 | -7.74 | 7.18 | -0.08 | 0.94 | 0.0013 |
| ELA:AT:Social | 4.25 | 4.48 | 108 | -4.62 | 13.12 | 0.95 | 0.3443 | 0.0008 |
| ELA:WT:Social | 7.26 | 3.67 | 108 | -0.02 | 14.53 | 1.98 | 0.0505 | 0.0007 |
| ELA:Male:Social | 3.87 | 4.19 | 108 | -4.43 | 12.18 | 0.92 | 0.3573 | 0.0006 |
| AT:Male:Social | 4.67 | 4.86 | 108 | -4.96 | 14.3 | 0.96 | 0.3386 | 0.0006 |
| WT:Male:Social | 1.79 | 4.06 | 108 | -6.26 | 9.84 | 0.44 | 0.6603 | 0.0003 |
| ELA:AT:Male:Social | -7.83 | 6.46 | 108 | -20.63 | 4.96 | -1.21 | 0.2277 | 0.0001 |
| ELA:WT:Male:Social | -5.73 | 5.32 | 108 | -16.28 | 4.82 | -1.08 | 0.2839 | 0 |

Model formula:
`lme( ~ Group*Strain*Sex*Compartment, random = ~ 1|ID/Compartment)`
$R^2_{marg} = 35.66\%$ , $R^2_{cond} > 99.99\%$
Abbreviations: $\beta$ = $\beta$ coefficient estimate, *SE* = standard error, Df = degrees of freedom, CI = 95% confidence interval, part = partial, group = early life condition group, marg = marginalized, cond = conditioned,

**Supplementary Table S24:**
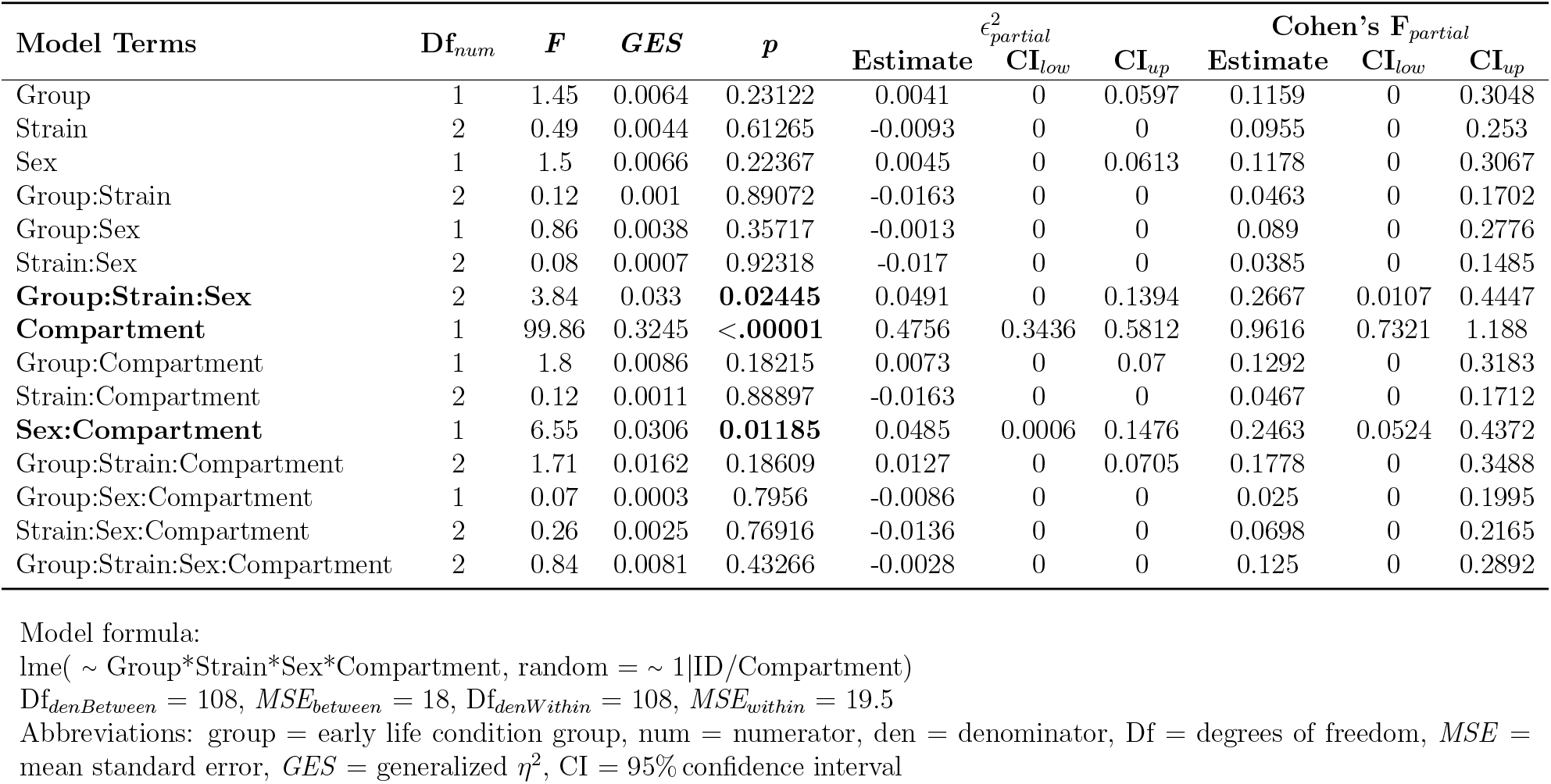
ANOVA Percent Interaction Time SCT.

| Model Terms | Df<sub>num</sub> | *F* | *GES* | *p* | $\epsilon^2_{\text{partial}}$ | | | Cohen's $F_{\text{partial}}$ | | |
| --- | --- | --- | --- | --- | --- | --- | --- | --- | --- | --- |
|  |  |  |  |  | Estimate | CI<sub>low</sub> | CI<sub>up</sub> | Estimate | CI<sub>low</sub> | CI<sub>up</sub> |
| Group | 1 | 1.45 | 0.0064 | 0.23122 | 0.0041 | 0 | 0.0597 | 0.1159 | 0 | 0.3048 |
| Strain | 2 | 0.49 | 0.0044 | 0.61265 | -0.0093 | 0 | 0 | 0.0955 | 0 | 0.253 |
| Sex | 1 | 1.5 | 0.0066 | 0.22367 | 0.0045 | 0 | 0.0613 | 0.1178 | 0 | 0.3067 |
| Group:Strain | 2 | 0.12 | 0.001 | 0.89072 | -0.0163 | 0 | 0 | 0.0463 | 0 | 0.1702 |
| Group:Sex | 1 | 0.86 | 0.0038 | 0.35717 | -0.0013 | 0 | 0 | 0.089 | 0 | 0.2776 |
| Strain:Sex | 2 | 0.08 | 0.0007 | 0.92318 | -0.017 | 0 | 0 | 0.0385 | 0 | 0.1485 |
| **Group:Strain:Sex** | 2 | 3.84 | 0.033 | **0.02445** | 0.0491 | 0 | 0.1394 | 0.2667 | 0.0107 | 0.4447 |
| **Compartment** | 1 | 99.86 | 0.3245 | **<.00001** | 0.4756 | 0.3436 | 0.5812 | 0.9616 | 0.7321 | 1.188 |
| Group:Compartment | 1 | 1.8 | 0.0086 | 0.18215 | 0.0073 | 0 | 0.07 | 0.1292 | 0 | 0.3183 |
| Strain:Compartment | 2 | 0.12 | 0.0011 | 0.88897 | -0.0163 | 0 | 0 | 0.0467 | 0 | 0.1712 |
| **Sex:Compartment** | 1 | 6.55 | 0.0306 | **0.01185** | 0.0485 | 0.0006 | 0.1476 | 0.2463 | 0.0524 | 0.4372 |
| Group:Strain:Compartment | 2 | 1.71 | 0.0162 | 0.18609 | 0.0127 | 0 | 0.0705 | 0.1778 | 0 | 0.3488 |
| Group:Sex:Compartment | 1 | 0.07 | 0.0003 | 0.7956 | -0.0086 | 0 | 0 | 0.025 | 0 | 0.1995 |
| Strain:Sex:Compartment | 2 | 0.26 | 0.0025 | 0.76916 | -0.0136 | 0 | 0 | 0.0698 | 0 | 0.2165 |
| Group:Strain:Sex:Compartment | 2 | 0.84 | 0.0081 | 0.43266 | -0.0028 | 0 | 0 | 0.125 | 0 | 0.2892 |

Model formula:
lme( ~ Group\*Strain\*Sex\*Compartment, random = ~ 1|ID/Compartment)
Df<sub>denBetween</sub> = 108, MSE<sub>between</sub> = 18, Df<sub>denWithin</sub> = 108, MSE<sub>within</sub> = 19.5
Abbreviations: group = early life condition group, num = numerator, den = denominator, Df = degrees of freedom, MSE = mean standard error, GES = generalized $\eta^2$ , CI = 95% confidence interval

**Supplementary Table S25:** Descriptive Social Distance [% time].

| Strain | Sex | Group | Compartment | N | Mean | SD | 95% CI<br>low | 95% CI<br>up |
| --- | --- | --- | --- | --- | --- | --- | --- | --- |
| CG | Female | Control | Reference | 7 | 7.51 | 2.50 | 5.20 | 9.83 |
|  |  |  | Social | 7 | 10.93 | 2.26 | 8.84 | 13.02 |
|  |  | ELA | Reference | 11 | 8.56 | 4.22 | 5.73 | 11.40 |
|  |  |  | Social | 11 | 21.39 | 7.70 | 16.22 | 26.57 |
|  | Male | Control | Reference | 7 | 8.33 | 3.92 | 4.71 | 11.95 |
|  |  |  | Social | 7 | 15.87 | 7.39 | 9.03 | 22.71 |
|  |  | ELA | Reference | 11 | 7.70 | 3.00 | 5.69 | 9.71 |
|  |  |  | Social | 11 | 15.73 | 2.81 | 13.84 | 17.62 |
| AT | Female | Control | Reference | 5 | 7.60 | 1.75 | 5.42 | 9.78 |
|  |  |  | Social | 5 | 15.36 | 10.48 | 2.34 | 28.38 |
|  |  | ELA | Reference | 10 | 7.59 | 4.52 | 4.36 | 10.82 |
|  |  |  | Social | 10 | 13.07 | 6.70 | 8.28 | 17.86 |
|  | Male | Control | Reference | 7 | 7.20 | 2.49 | 4.89 | 9.51 |
|  |  |  | Social | 7 | 18.43 | 4.53 | 14.24 | 22.62 |
|  |  | ELA | Reference | 5 | 3.90 | 2.74 | 0.50 | 7.30 |
|  |  |  | Social | 5 | 11.72 | 9.51 | -0.08 | 23.52 |
| WT | Female | Control | Reference | 16 | 8.77 | 3.78 | 6.76 | 10.78 |
|  |  |  | Social | 16 | 11.49 | 4.82 | 8.93 | 14.06 |
|  |  | ELA | Reference | 16 | 7.40 | 3.59 | 5.49 | 9.31 |
|  |  |  | Social | 16 | 13.06 | 5.91 | 9.91 | 16.21 |
|  | Male | Control | Reference | 11 | 7.04 | 4.33 | 4.13 | 9.95 |
|  |  |  | Social | 11 | 13.89 | 5.47 | 10.21 | 17.57 |
|  |  | ELA | Reference | 14 | 6.82 | 3.67 | 4.70 | 8.94 |
|  |  |  | Social | 14 | 13.89 | 7.01 | 9.84 | 17.93 |

**Supplementary Table S26:** Model Summary Distant Observation Time SCT.

| **Model Terms** | $\beta$ | ***SE*** | **Df** | $CI_{low}$ | $CI_{up}$ | $t$ | $p$ | $R^2_{part}$ |
| --- | --- | --- | --- | --- | --- | --- | --- | --- |
| Intercept | 7.51 | 1.92 | 108 | 3.7 | 11.33 | 3.9 | 0.0002 | 0.3928 |
| ELA | 1.05 | 2.46 | 108 | -3.83 | 5.93 | 0.43 | 0.6708 | 0.03 |
| AT | 0.09 | 2.98 | 108 | -5.82 | 6 | 0.03 | 0.9771 | 0.0205 |
| WT | 1.25 | 2.31 | 108 | -3.32 | 5.83 | 0.54 | 0.5878 | 0.0137 |
| Male | 0.81 | 2.72 | 108 | -4.58 | 6.21 | 0.3 | 0.7654 | 0.0095 |
| Social | 3.41 | 2.72 | 108 | -1.97 | 8.8 | 1.26 | 0.2115 | 0.0066 |
| ELA:AT | -1.06 | 3.72 | 108 | -8.43 | 6.31 | -0.28 | 0.7764 | 0.0049 |
| ELA:WT | -2.42 | 3.05 | 108 | -8.46 | 3.63 | -0.79 | 0.4296 | 0.0045 |
| ELA:Male | -1.68 | 3.48 | 108 | -8.58 | 5.22 | -0.48 | 0.6308 | 0.0045 |
| AT:Male | -1.21 | 4.04 | 108 | -9.22 | 6.79 | -0.3 | 0.7641 | 0.0042 |
| WT:Male | -2.55 | 3.37 | 108 | -9.23 | 4.14 | -0.75 | 0.452 | 0.0027 |
| ELA:Social | 9.41 | 3.47 | 108 | 2.53 | 16.3 | 2.71 | 0.0078 | 0.0024 |
| AT:Social | 4.35 | 4.21 | 108 | -4 | 12.69 | 1.03 | 0.304 | 0.0017 |
| WT:Social | -0.69 | 3.26 | 108 | -7.14 | 5.77 | -0.21 | 0.8328 | 0.0013 |
| Male:Social | 4.13 | 3.84 | 108 | -3.49 | 11.74 | 1.07 | 0.2849 | 0.001 |
| ELA:AT:Male | -1.61 | 5.37 | 108 | -12.25 | 9.02 | -0.3 | 0.7644 | 0.0008 |
| ELA:WT:Male | 2.83 | 4.42 | 108 | -5.94 | 11.6 | 0.64 | 0.5235 | 0.0004 |
| ELA:AT:Social | -11.69 | 5.25 | 108 | -22.1 | -1.29 | -2.23 | 0.028 | 0.0004 |
| ELA:WT:Social | -6.48 | 4.3 | 108 | -15.01 | 2.05 | -1.51 | 0.135 | 0.0004 |
| ELA:Male:Social | -8.93 | 4.91 | 108 | -18.67 | 0.81 | -1.82 | 0.072 | 0.0003 |
| AT:Male:Social | -0.66 | 5.7 | 108 | -11.95 | 10.63 | -0.12 | 0.908 | 0.0002 |
| WT:Male:Social | 0 | 4.76 | 108 | -9.44 | 9.44 | 0 | 0.9998 | 0.0001 |
| ELA:AT:Male:Social | 7.8 | 7.57 | 108 | -7.21 | 22.81 | 1.03 | 0.3053 | 0 |
| ELA:WT:Male:Social | 6.21 | 6.24 | 108 | -6.17 | 18.58 | 0.99 | 0.3224 | 0 |

Model formula:
`lme( ~ Group*Strain*Sex*Compartment, random = ~ 1|ID/Compartment)`
$R^2_{marg} = 38.98\%$ , $R^2_{cond} = 98.93\%$
Abbreviations: $\beta$ = $\beta$ coefficient estimate, $SE$ = standard error, Df = degrees of freedom, CI = 95% confidence interval, part = partial, group = early life condition group, marg = marginalized, cond = conditioned,

**Supplementary Table S27:**
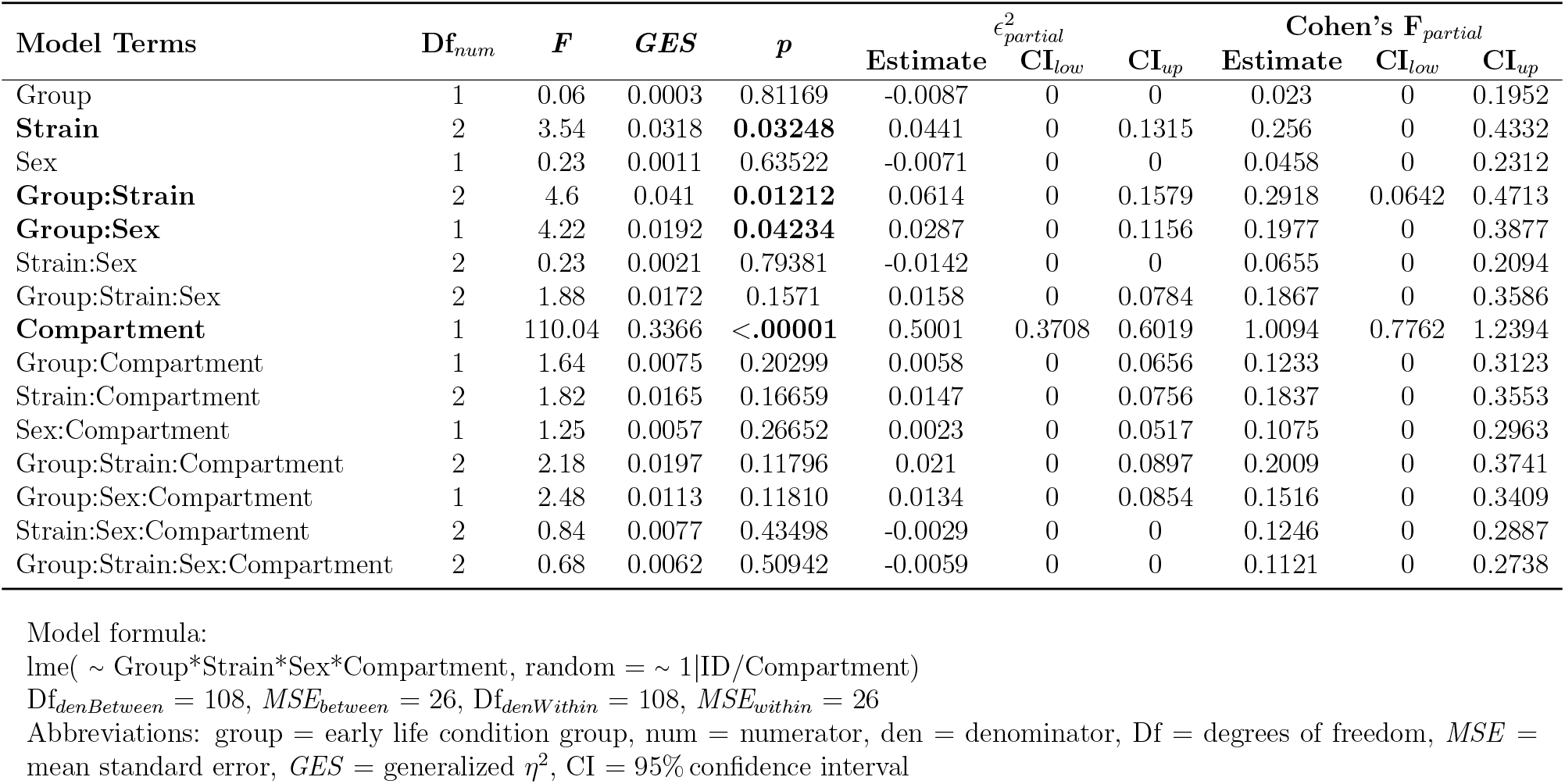
ANOVA Percent Distant Observation Time SCT.

| Model Terms | Df<sub>num</sub> | F | GES | p | $\epsilon_{partial}^2$ | | | Cohen's F<sub>partial</sub> | | |
| --- | --- | --- | --- | --- | --- | --- | --- | --- | --- | --- |
|  |  |  |  |  | Estimate | CI<sub>low</sub> | CI<sub>up</sub> | Estimate | CI<sub>low</sub> | CI<sub>up</sub> |
| Group | 1 | 0.06 | 0.0003 | 0.81169 | -0.0087 | 0 | 0 | 0.023 | 0 | 0.1952 |
| **Strain** | 2 | 3.54 | 0.0318 | **0.03248** | 0.0441 | 0 | 0.1315 | 0.256 | 0 | 0.4332 |
| Sex | 1 | 0.23 | 0.0011 | 0.63522 | -0.0071 | 0 | 0 | 0.0458 | 0 | 0.2312 |
| **Group:Strain** | 2 | 4.6 | 0.041 | **0.01212** | 0.0614 | 0 | 0.1579 | 0.2918 | 0.0642 | 0.4713 |
| **Group:Sex** | 1 | 4.22 | 0.0192 | **0.04234** | 0.0287 | 0 | 0.1156 | 0.1977 | 0 | 0.3877 |
| Strain:Sex | 2 | 0.23 | 0.0021 | 0.79381 | -0.0142 | 0 | 0 | 0.0655 | 0 | 0.2094 |
| Group:Strain:Sex | 2 | 1.88 | 0.0172 | 0.1571 | 0.0158 | 0 | 0.0784 | 0.1867 | 0 | 0.3586 |
| **Compartment** | 1 | 110.04 | 0.3366 | **<.00001** | 0.5001 | 0.3708 | 0.6019 | 1.0094 | 0.7762 | 1.2394 |
| Group:Compartment | 1 | 1.64 | 0.0075 | 0.20299 | 0.0058 | 0 | 0.0656 | 0.1233 | 0 | 0.3123 |
| Strain:Compartment | 2 | 1.82 | 0.0165 | 0.16659 | 0.0147 | 0 | 0.0756 | 0.1837 | 0 | 0.3553 |
| Sex:Compartment | 1 | 1.25 | 0.0057 | 0.26652 | 0.0023 | 0 | 0.0517 | 0.1075 | 0 | 0.2963 |
| Group:Strain:Compartment | 2 | 2.18 | 0.0197 | 0.11796 | 0.021 | 0 | 0.0897 | 0.2009 | 0 | 0.3741 |
| Group:Sex:Compartment | 1 | 2.48 | 0.0113 | 0.11810 | 0.0134 | 0 | 0.0854 | 0.1516 | 0 | 0.3409 |
| Strain:Sex:Compartment | 2 | 0.84 | 0.0077 | 0.43498 | -0.0029 | 0 | 0 | 0.1246 | 0 | 0.2887 |
| Group:Strain:Sex:Compartment | 2 | 0.68 | 0.0062 | 0.50942 | -0.0059 | 0 | 0 | 0.1121 | 0 | 0.2738 |

Model formula:
lme( ~ Group\*Strain\*Sex\*Compartment, random = ~ 1|ID/Compartment)
Df<sub>denBetween</sub> = 108, MSE<sub>between</sub> = 26, Df<sub>denWithin</sub> = 108, MSE<sub>within</sub> = 26
Abbreviations: group = early life condition group, num = numerator, den = denominator, Df = degrees of freedom, MSE = mean standard error, GES = generalized $\eta^2$ , CI = 95% confidence interval

**Supplementary Table S28:** Model Summary Restraint Stress Corticosterone.

| **Model Terms** | $\beta$ | ***SE*** | **Df** | ***CI<sub>low</sub>*** | ***CI<sub>up</sub>*** | ***t*** | ***p*** | ***R<sub>part</sub><sup>2</sup>*** |
| --- | --- | --- | --- | --- | --- | --- | --- | --- |
| Intercept | 12.29 | 14.04 | 122 | -14.21 | 38.78 | 0.88 | 0.3832 | 0.7238 |
| ELA | 5.35 | 17.66 | 122 | -27.98 | 38.69 | 0.3 | 0.7623 | 0.0708 |
| AT | 6.86 | 19.85 | 122 | -30.61 | 44.33 | 0.35 | 0.7304 | 0.0058 |
| WT | 5.4 | 17 | 122 | -26.68 | 37.49 | 0.32 | 0.7511 | 0.0031 |
| Female | 16.57 | 19.85 | 122 | -20.9 | 54.04 | 0.83 | 0.4055 | 0.0029 |
| Restraint | 85 | 19.48 | 118 | 48.22 | 121.78 | 4.36 | 0 | 0.002 |
| ELA:AT | -7.76 | 28.01 | 122 | -60.64 | 45.11 | -0.28 | 0.7822 | 0.0013 |
| ELA:WT | -0.25 | 22.14 | 122 | -42.04 | 41.54 | -0.01 | 0.991 | 0.0007 |
| ELA:Female | 2.97 | 25.19 | 122 | -44.57 | 50.51 | 0.12 | 0.9063 | 0.0007 |
| AT:Female | 4.79 | 28.65 | 122 | -49.29 | 58.87 | 0.17 | 0.8676 | 0.0006 |
| WT:Female | -4.87 | 23.56 | 122 | -49.33 | 39.6 | -0.21 | 0.8366 | 0.0005 |
| ELA:Restraint | 0.28 | 24.51 | 118 | -46 | 46.56 | 0.01 | 0.991 | 0.0004 |
| AT:Restraint | -0.43 | 27.55 | 118 | -52.44 | 51.58 | -0.02 | 0.9876 | 0.0004 |
| WT:Restraint | -9.09 | 23.59 | 118 | -53.63 | 35.45 | -0.39 | 0.7007 | 0.0003 |
| Female:Restraint | 33.29 | 27.55 | 118 | -18.73 | 85.3 | 1.21 | 0.2293 | 0.0003 |
| ELA:AT:Female | 9.44 | 38.24 | 122 | -62.74 | 81.62 | 0.25 | 0.8055 | 0.0003 |
| ELA:WT:Female | 0.22 | 31.03 | 122 | -58.34 | 58.78 | 0.01 | 0.9943 | 0.0002 |
| ELA:AT:Restraint | -27.88 | 39.75 | 118 | -102.93 | 47.18 | -0.7 | 0.4845 | 0.0001 |
| ELA:WT:Restraint | 3.89 | 30.72 | 118 | -54.11 | 61.9 | 0.13 | 0.8993 | 0.0001 |
| ELA:Female:Restraint | 14.89 | 34.95 | 118 | -51.1 | 80.88 | 0.43 | 0.6708 | 0.0001 |
| AT:Female:Restraint | -23.17 | 40.33 | 118 | -99.32 | 52.98 | -0.57 | 0.5667 | 0 |
| WT:Female:Restraint | 13.98 | 32.75 | 118 | -47.84 | 75.81 | 0.43 | 0.6701 | 0 |
| ELA:AT:Female:Restraint | 47.52 | 54.14 | 118 | -54.7 | 149.74 | 0.88 | 0.3819 | 0 |
| ELA:WT:Female:Restraint | -11.53 | 43.15 | 118 | -93.01 | 69.95 | -0.27 | 0.7897 | 0 |

Model formula:
`lme( ~ Group*Strain*Sex*SampleType, random = ~ 1|ID/SampleType)`
$R^2_{marg} = 72.52\%$ , $R^2_{cond} = 98.61\%$
Abbreviations: $\beta$ = $\beta$ coefficient estimate, *SE* = standard error, Df = degrees of freedom, CI = 95% confidence interval, part = partial, group = early life condition group, marg = marginalized, cond = conditioned,

**Supplementary Table S29:** ANOVA Restraint Stress Corticosterone.

| Model Terms | $Df_{num}$ | $F$ | $GES$ | $p$ | $\epsilon_{partial}^2$ | | | Cohen's $F_{partial}$ | | |
| --- | --- | --- | --- | --- | --- | --- | --- | --- | --- | --- |
| | | | | | Estimate | $CI_{low}$ | $CI_{up}$ | Estimate | $CI_{low}$ | $CI_{up}$ |
| **Group** | 1 | 4.27 | 0.0184 | **0.04101** | 0.0267 | 0 | 0.1075 | 0.1902 | 0 | 0.3719 |
| Strain | 2 | 0.15 | 0.0013 | 0.86509 | -0.0145 | 0 | 0 | 0.0496 | 0 | 0.1753 |
| **Sex** | 1 | 70.59 | 0.2369 | **<0.00001** | 0.369 | 0.2377 | 0.484 | 0.7735 | 0.5666 | 0.9778 |
| Group:Strain | 2 | 0.02 | 0.0002 | 0.98166 | -0.0166 | 0 | 0 | 0.0177 | 0 | 0 |
| Group:Sex | 1 | 2.04 | 0.0089 | 0.15560 | 0.0087 | 0 | 0.0699 | 0.1316 | 0 | 0.3125 |
| Strain:Sex | 2 | 0.32 | 0.0028 | 0.72473 | -0.0114 | 0 | 0 | 0.074 | 0 | 0.2182 |
| Group:Strain:Sex | 2 | 1.07 | 0.0093 | 0.34799 | 0.0011 | 0 | 0.0156 | 0.1344 | 0 | 0.2937 |
| **SampleType** | 1 | 516.69 | 0.6781 | **<0.00001** | 0.8125 | 0.7558 | 0.8522 | 2.0925 | 1.7691 | 2.413 |
| Group:SampleType | 1 | 0.5 | 0.002 | 0.48276 | -0.0043 | 0 | 0 | 0.0648 | 0 | 0.2447 |
| Strain:SampleType | 2 | 0.37 | 0.003 | 0.69132 | -0.0106 | 0 | 0 | 0.0792 | 0 | 0.2257 |
| **Sex:SampleType** | 1 | 24.08 | 0.0894 | **<0.00001** | 0.1624 | 0.0591 | 0.2831 | 0.4517 | 0.2614 | 0.6402 |
| Group:Strain:SampleType | 2 | 0.01 | 0.0001 | 0.99149 | -0.0168 | 0 | 0 | 0.012 | 0 | 0 |
| Group:Sex:SampleType | 1 | 0.89 | 0.0036 | 0.34817 | -0.0009 | 0 | 0 | 0.0867 | 0 | 0.2672 |
| Strain:Sex:SampleType | 2 | 0.08 | 0.0007 | 0.91888 | -0.0155 | 0 | 0 | 0.0379 | 0 | 0.1443 |
| Group:Strain:Sex:SampleType | 2 | 0.76 | 0.0061 | 0.47092 | -0.0041 | 0 | 0 | 0.1133 | 0 | 0.2692 |

Model formula:
`lme( ~ Group*Strain*Sex*SampleType, random = ~ 1|ID/SampleType)`
$Df_{denBetween} = 118$ , $MSE_{between} = 1454$ , $Df_{denWithin} = 118$ , $MSE_{within} = 1348$
Abbreviations: group = early life condition group, num = numerator, den = denominator, Df = degrees of freedom, $MSE$ = mean standard error, $GES$ = generalized $\eta^2$ , CI = 95% confidence interval

**Supplementary Table S30:** Model Summary Dexamethasone Suppression.

| **Model Terms** | $\beta$ | ***SE*** | **Df** | **CI<sub>low</sub>** | **CI<sub>up</sub>** | ***t*** | ***p*** | **$R^2_{part}$** |
| --- | --- | --- | --- | --- | --- | --- | --- | --- |
| Intercept | 12.29 | 4.03 | 123 | 4.68 | 19.89 | 3.05 | 0.0028 | 0.6342 |
| ELA | 5.35 | 5.07 | 123 | -4.22 | 14.92 | 1.06 | 0.2929 | 0.034 |
| AT | 6.86 | 5.7 | 123 | -3.9 | 17.61 | 1.2 | 0.2309 | 0.0161 |
| WT | 5.4 | 4.88 | 123 | -3.81 | 14.61 | 1.11 | 0.2701 | 0.0064 |
| Female | 16.57 | 5.7 | 123 | 5.82 | 27.33 | 2.91 | 0.0043 | 0.006 |
| Dex | -11.29 | 5.66 | 120 | -21.97 | -0.6 | -1.99 | 0.0484 | 0.0051 |
| ELA:AT | -7.76 | 8.04 | 123 | -22.94 | 7.42 | -0.97 | 0.3361 | 0.0046 |
| ELA:WT | -0.26 | 6.35 | 123 | -12.25 | 11.74 | -0.04 | 0.9675 | 0.0039 |
| ELA:Female | 2.97 | 7.23 | 123 | -10.68 | 16.62 | 0.41 | 0.6817 | 0.0031 |
| AT:Female | 4.79 | 8.22 | 123 | -10.74 | 20.31 | 0.58 | 0.5615 | 0.0027 |
| WT:Female | -4.87 | 6.76 | 123 | -17.63 | 7.9 | -0.72 | 0.4727 | 0.0026 |
| ELA:Dex | -3.52 | 7.12 | 120 | -16.97 | 9.93 | -0.49 | 0.622 | 0.0022 |
| AT:Dex | -3.14 | 8 | 120 | -18.26 | 11.97 | -0.39 | 0.6952 | 0.0019 |
| WT:Dex | -3.63 | 6.89 | 120 | -16.64 | 9.39 | -0.53 | 0.5998 | 0.0014 |
| Female:Dex | -10 | 8 | 120 | -25.12 | 5.12 | -1.25 | 0.2139 | 0.0013 |
| ELA:AT:Female | 9.44 | 10.97 | 123 | -11.28 | 30.16 | 0.86 | 0.3914 | 0.0013 |
| ELA:WT:Female | 0.23 | 8.9 | 123 | -16.58 | 17.04 | 0.03 | 0.9795 | 0.0011 |
| ELA:AT:Dex | 4.42 | 11.29 | 120 | -16.92 | 25.75 | 0.39 | 0.6965 | 0.001 |
| ELA:WT:Dex | 7.24 | 8.93 | 120 | -9.64 | 24.11 | 0.81 | 0.4195 | 0.0007 |
| ELA:Female:Dex | -6.83 | 10.15 | 120 | -26.01 | 12.35 | -0.67 | 0.5024 | 0.0006 |
| AT:Female:Dex | -9.24 | 11.55 | 120 | -31.06 | 12.58 | -0.8 | 0.4254 | 0.0006 |
| WT:Female:Dex | 5.42 | 9.53 | 120 | -12.57 | 23.41 | 0.57 | 0.5704 | 0.0001 |
| ELA:AT:Female:Dex | -1.75 | 15.45 | 120 | -30.93 | 27.44 | -0.11 | 0.9102 | 0 |
| ELA:WT:Female:Dex | -6.96 | 12.51 | 120 | -30.6 | 16.67 | -0.56 | 0.579 | 0 |

Model formula:
`lme( ~ Group*Strain*Sex*SampleType, random = ~ 1|ID/SampleType)`
$R^2_{marg} = 63.25\%$ , $R^2_{cond} = 98.86\%$
Abbreviations: $\beta$ = $\beta$ coefficient estimate, *SE* = standard error, Df = degrees of freedom, CI = 95% confidence interval, part = partial, group = early life condition group, marg = marginalized, cond = conditioned,

**Supplementary Table S31:** ANOVA Dexamethasone Suppression Test.

| Model Terms | $Df_{num}$ | $F$ | $GES$ | $p$ | $\epsilon_{partial}^2$ | | | Cohen's $F_{partial}$ | | |
| --- | --- | --- | --- | --- | --- | --- | --- | --- | --- | --- |
| | | | | | Estimate | $CI_{low}$ | $CI_{up}$ | Estimate | $CI_{low}$ | $CI_{up}$ |
| **Group** | 1 | 8.48 | 0.0345 | **0.00429** | 0.0582 | 0.0043 | 0.1559 | 0.2658 | 0.083 | 0.4473 |
| **Strain** | 2 | 4.16 | 0.0339 | **0.01793** | 0.0492 | 0 | 0.1346 | 0.2633 | 0.0402 | 0.4325 |
| **Sex** | 1 | 58.99 | 0.1992 | **<0.00001** | 0.324 | 0.1949 | 0.4422 | 0.7011 | 0.5003 | 0.8996 |
| Group:Strain | 2 | 0.54 | 0.0045 | 0.58400 | -0.0076 | 0 | 0 | 0.0949 | 0 | 0.2455 |
| Group:Sex | 1 | 0.09 | 0.0004 | 0.76401 | -0.0076 | 0 | 0 | 0.0275 | 0 | 0.1964 |
| Strain:Sex | 2 | 2.75 | 0.0227 | 0.06807 | 0.0279 | 0 | 0.099 | 0.214 | 0 | 0.3799 |
| Group:Strain:Sex | 2 | 1.05 | 0.0088 | 0.35248 | 0.0008 | 0 | 0.0116 | 0.1324 | 0 | 0.2903 |
| **SampleType** | 1 | 276.49 | 0.5323 | **<0.00001** | 0.6948 | 0.6082 | 0.7578 | 1.5179 | 1.254 | 1.7787 |
| Group:SampleType | 1 | 2.38 | 0.0097 | 0.12583 | 0.0112 | 0 | 0.0755 | 0.1407 | 0 | 0.3202 |
| Strain:SampleType | 2 | 1.86 | 0.0151 | 0.15992 | 0.0139 | 0 | 0.0701 | 0.1761 | 0 | 0.339 |
| **Sex:SampleType** | 1 | 31.42 | 0.1145 | **<0.00001** | 0.2009 | 0.088 | 0.3227 | 0.5117 | 0.3205 | 0.701 |
| Group:Strain:SampleType | 2 | 0.36 | 0.003 | 0.69861 | -0.0106 | 0 | 0 | 0.0774 | 0 | 0.2222 |
| **Group:Sex:SampleType** | 1 | 4.26 | 0.0172 | **0.04108** | 0.0263 | 0 | 0.1058 | 0.1885 | 0 | 0.3686 |
| Strain:Sex:SampleType | 2 | 1.15 | 0.0094 | 0.32064 | 0.0024 | 0 | 0.0294 | 0.1383 | 0 | 0.297 |
| Group:Strain:Sex:SampleType | 2 | 0.4 | 0.0033 | 0.66930 | -0.0099 | 0 | 0 | 0.0819 | 0 | 0.2285 |

Model formula:
$\text{lme}(\sim \text{Group} * \text{Strain} * \text{Sex} * \text{SampleType}, \text{random} = \sim 1 | \text{ID} / \text{SampleType})$
$Df_{denBetween} = 120$ , $MSE_{between} = 108$ , $Df_{denWithin} = 120$ , $MSE_{within} = 106$
Abbreviations: group = early life condition group, num = numerator, den = denominator, Df = degrees of freedom, $MSE$ = mean standard error, $GES$ = generalized $\eta^2$ , CI = 95% confidence interval

**Supplementary Table S32:** Descriptive Statistics of Glucocorticoid Signaling Regulating Genes Separated by Brain Region, *Fkbp5*-Genotype and Early Life Condition.

| Tissue | Strain | Group | Gene | N | Mean | SD | 95% CI<br>low | 95% CI<br>up |
| --- | --- | --- | --- | --- | --- | --- | --- | --- |
| Hypothalamus | CG | Control | *Nr3c2* | 5 | 4.32 | 0.98 | 3.10 | 5.54 |
|  | CG | ELA |  | 5 | 4.20 | 0.25 | 3.89 | 4.51 |
|  | AT | Control |  | 4 | 4.26 | 0.41 | 3.62 | 4.91 |
|  | AT | ELA |  | 7 | 3.74 | 1.54 | 2.32 | 5.16 |
| Ventral<br>Hippocampus | CG | Control |  | 4 | 26.58 | 3.88 | 20.41 | 32.76 |
|  | CG | ELA |  | 5 | 19.32 | 8.35 | 8.95 | 29.69 |
|  | AT | Control |  | 3 | 21.26 | 7.60 | 2.37 | 40.15 |
|  | AT | ELA |  | 7 | 13.93 | 5.79 | 8.57 | 19.28 |
| Dorsal<br>Hippocampus | CG | Control |  | 4 | 26.58 | 3.88 | 20.41 | 32.76 |
|  | CG | ELA |  | 5 | 19.32 | 8.35 | 8.95 | 29.69 |
|  | AT | Control |  | 3 | 21.26 | 7.60 | 2.37 | 40.15 |
|  | AT | ELA |  | 7 | 13.93 | 5.79 | 8.57 | 19.28 |
| Hypothalamus | CG | Control | *Nr3c1* | 5 | 8.32 | 4.85 | 2.30 | 14.35 |
|  | CG | ELA |  | 5 | 8.95 | 2.62 | 5.70 | 12.21 |
|  | AT | Control |  | 4 | 7.36 | 4.36 | 0.42 | 14.30 |
|  | AT | ELA |  | 7 | 7.41 | 3.30 | 4.36 | 10.46 |
| Ventral<br>Hippocampus | CG | Control |  | 4 | 9.30 | 0.87 | 7.93 | 10.68 |
|  | CG | ELA |  | 5 | 8.46 | 2.14 | 5.81 | 11.11 |
|  | AT | Control |  | 3 | 9.74 | 3.32 | 1.49 | 17.99 |
|  | AT | ELA |  | 7 | 6.54 | 3.59 | 3.22 | 9.86 |
| Dorsal<br>Hippocampus | CG | Control |  | 4 | 9.30 | 0.87 | 7.93 | 10.68 |
|  | CG | ELA |  | 5 | 8.46 | 2.14 | 5.81 | 11.11 |
|  | AT | Control |  | 3 | 9.74 | 3.32 | 1.49 | 17.99 |
|  | AT | ELA |  | 7 | 6.54 | 3.59 | 3.22 | 9.86 |
| Hypothalamus | CG | Control | *Hsp90ab1* | 5 | 719.29 | 58.80 | 646.29 | 792.29 |
|  | CG | ELA |  | 5 | 747.51 | 27.66 | 713.16 | 781.85 |
|  | AT | Control |  | 4 | 645.67 | 46.85 | 571.11 | 720.22 |
|  | AT | ELA |  | 7 | 642.06 | 54.79 | 591.39 | 692.73 |
| Ventral<br>Hippocampus | CG | Control |  | 4 | 772.59 | 32.21 | 721.34 | 823.84 |
|  | CG | ELA |  | 5 | 768.25 | 51.96 | 703.73 | 832.77 |
|  | AT | Control |  | 3 | 707.60 | 63.99 | 548.63 | 866.58 |
|  | AT | ELA |  | 7 | 645.62 | 98.51 | 554.52 | 736.73 |
| Dorsal<br>Hippocampus | CG | Control |  | 4 | 772.59 | 32.21 | 721.34 | 823.84 |
|  | CG | ELA |  | 5 | 768.25 | 51.96 | 703.73 | 832.77 |
|  | AT | Control |  | 3 | 707.60 | 63.99 | 548.63 | 866.58 |
|  | AT | ELA |  | 7 | 645.62 | 98.51 | 554.52 | 736.73 |
| Hypothalamus | CG | Control | *Fkbp5* | 5 | 2.80 | 0.61 | 2.04 | 3.56 |
|  | CG | ELA |  | 5 | 2.65 | 0.43 | 2.12 | 3.18 |
|  | AT | Control |  | 4 | 3.20 | 0.47 | 2.45 | 3.95 |
|  | AT | ELA |  | 7 | 2.47 | 1.40 | 1.18 | 3.77 |
| Ventral<br>Hippocampus | CG | Control |  | 4 | 8.59 | 1.98 | 5.44 | 11.75 |
|  | CG | ELA |  | 5 | 7.18 | 2.53 | 4.04 | 10.32 |
|  | AT | Control |  | 3 | 5.92 | 0.36 | 5.03 | 6.81 |
|  | AT | ELA |  | 7 | 4.57 | 1.97 | 2.75 | 6.39 |
| Dorsal<br>Hippocampus | CG | Control |  | 4 | 8.59 | 1.98 | 5.44 | 11.75 |
|  | CG | ELA |  | 5 | 7.18 | 2.53 | 4.04 | 10.32 |
|  | AT | Control |  | 3 | 5.92 | 0.36 | 5.03 | 6.81 |
|  | AT | ELA |  | 7 | 4.57 | 1.97 | 2.75 | 6.39 |
| Hypothalamus | CG | Control | *Fkbp4* | 5 | 118.83 | 14.68 | 100.60 | 137.06 |
|  | CG | ELA |  | 5 | 122.42 | 5.26 | 115.89 | 128.95 |
|  | AT | Control |  | 4 | 118.67 | 17.79 | 90.36 | 146.97 |
|  | AT | ELA |  | 7 | 122.26 | 11.40 | 111.72 | 132.80 |
| Ventral<br>Hippocampus | CG | Control |  | 4 | 79.13 | 8.88 | 65.00 | 93.26 |
|  | CG | ELA |  | 5 | 91.14 | 16.06 | 71.19 | 111.09 |
|  | AT | Control |  | 3 | 83.57 | 16.58 | 42.40 | 124.75 |
|  | AT | ELA |  | 7 | 79.59 | 11.16 | 69.27 | 89.92 |
| Dorsal<br>Hippocampus | CG | Control |  | 4 | 79.13 | 8.88 | 65.00 | 93.26 |
|  | CG | ELA |  | 5 | 91.14 | 16.06 | 71.19 | 111.09 |
|  | AT | Control |  | 3 | 83.57 | 16.58 | 42.40 | 124.75 |
|  | AT | ELA |  | 7 | 79.59 | 11.16 | 69.27 | 89.92 |

**Supplementary Table S33:** Significant Results of the ANOVA on TissueWise Expression of Glucocorticoid Signaling Regulators in Dependence of *Fkbp5*-Genotype × Early Life Condition.

| **Tissue** | **Gene** | **Term** | **Df** | ***F*** | ***p*** | ***R*<sup>2</sup>** | ***R*<sub>adj</sub><sup>2</sup>** |
| --- | --- | --- | --- | --- | --- | --- | --- |
| Hypothalamus | *Hsp90ab1* | Strain | 1 | 17.7 | .0006 | 52 | 43 |
| Ventral<br>Hippocampus | *Hsp90ab1* | Strain | 1 | 8.9 | .009 | 43 | 32 |
|  | *Nr3c2* | Early Life | 1 | 5.4 | .03 | 40 | 28 |
|  | *Fkbp5* | Strain | 1 | 7.9 | .01 | 43 | 32 |
| Dorsal<br>Hippocampus | *Hsp90ab1* | Strain | 1 | 8.9 | 0.009 | 43 | 32 |
|  | *Nr3c2* | Early Life | 1 | 5.4 | 0.03 | 40 | 28 |
|  | *Fkbp5* | Strain | 1 | 7.9 | 0.01 | 43 | 32 |

**Supplementary Table S34:**
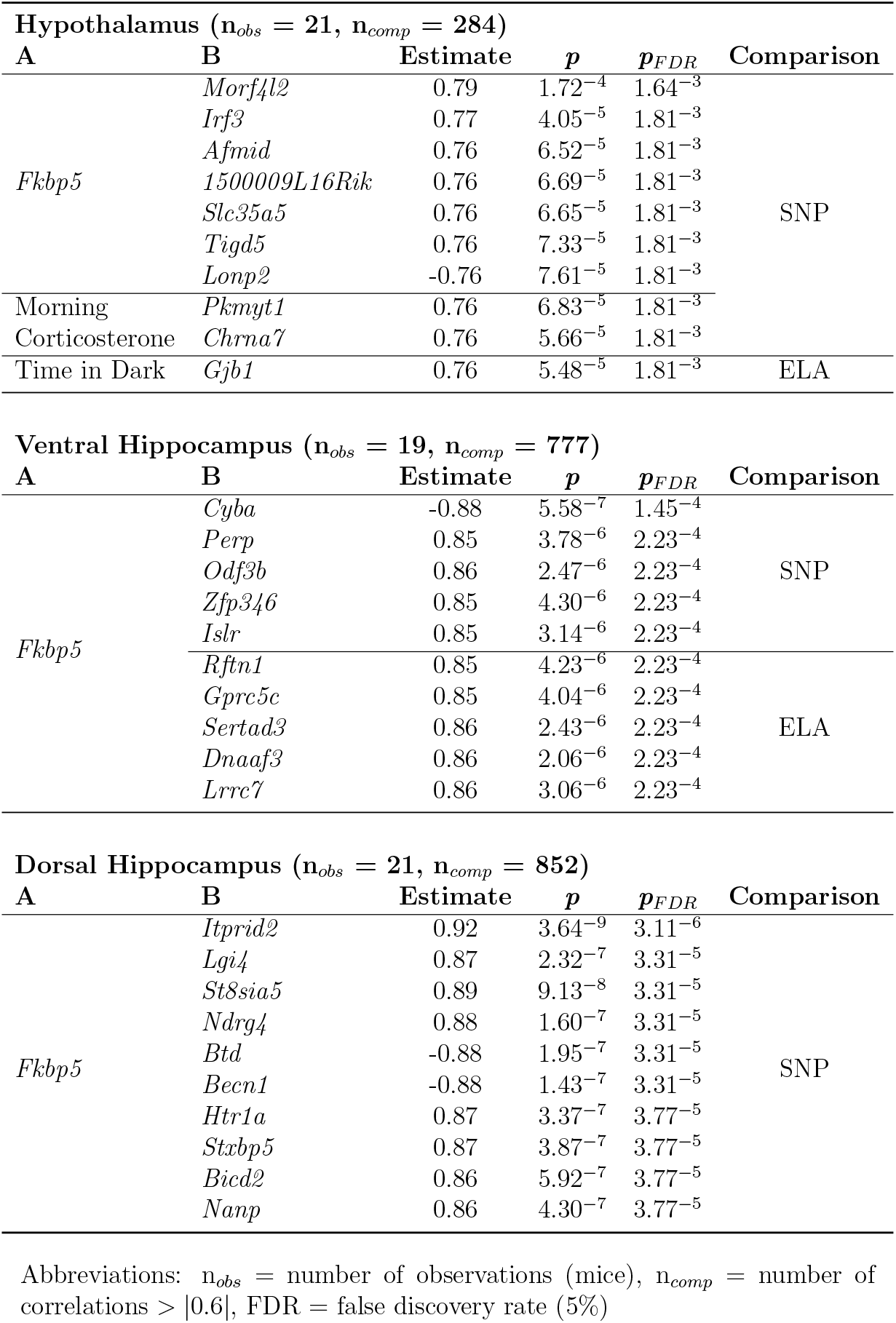
Top 10 Pearson Correlations between DEGs, Behavior, Physiology and *Fkbp5* Expression.

## References

[1] Selye, H. Stress and the general adaptation syndrome. BMJ 1, 1383–1392 (1950). URL https://doi.org/10.1136/bmj.1.4667.1383.

[2] de Kloet, E. R., Joëls, M. & Holsboer, F. Stress and the brain: from adaptation to disease. Nature Reviews Neuroscience 6, 463–475 (2005). URL https://doi.org/10.1038/nrn1683.

[3] Matosin, N., Halldorsdottir, T. & Binder, E. B. Understanding the molecular mechanisms underpinning gene by environment interactions in psychiatric disorders: The FKBP5 model. Biological Psychiatry 83, 821–830 (2018). URL https://doi.org/10.1016/j.biopsych.2018.01.021.

[4] Opendak, M., Gould, E. & Sullivan, R. Early life adversity during the infant sensitive period for attachment: Programming of behavioral neurobiology of threat processing and social behavior. Developmental Cognitive Neuroscience 25, 145–159 (2017). URL https://doi.org/10.1016/j.dcn.2017.02.002.

[5] MacMillan, H. L. et al. Childhood abuse and lifetime psychopathology in a community sample. American Journal of Psychiatry 158, 1878–1883 (2001). URL https://doi.org/10.1176/appi.ajp.158.11.1878.

[6] Nemeroff, C. B. Paradise lost: The neurobiological and clinical consequences of child abuse and neglect. Neuron 89, 892–909 (2016). URL https://doi.org/10.1016/j.neuron.2016.01.019.

[7] Nold, V., Richter, N., Hengerer, B., Kolassa, I.-T. & Allers, K. A. FKBP5 polymorphisms induce differential glucocorticoid responsiveness in primary CNS cells – first insights from novel humanized mice. European Journal of Neuro-science (2020). URL https://doi.org/10.1111/ejn.14999.

[8] Pirkl, F. & Buchner, J. Functional analysis of the hsp90-associated human peptidyl prolyl cis/trans isomerases FKBP51, FKBP52 and cyp40 1 1edited by r. huber. Journal of Molecular Biology 308, 795–806 (2001). URL https://doi.org/10.1006/jmbi.2001.4595.

[9] Binder, E. B. et al. Polymorphisms in FKBP5 are associated with increased recurrence of depressive episodes and rapid response to antidepressant treatment. Nature Genetics 36, 1319–1325 (2004). URL https://doi.org/10.1038/ng1479.

[10] Ising, M. et al. Polymorphisms in the FKBP5 gene region modulate recovery from psychosocial stress in healthy controls. European Journal of Neuroscience 28, 389–398 (2008). URL https://doi.org/10.1111/j.1460-9568.2008.06332.x.

[11] Dackis, M. N., Rogosch, F. A., Oshri, A. & Cicchetti, D. The role of limbic system irritability in linking history of childhood maltreatment and psychiatric outcomes in low-income, high-risk women: Moderation by FK506 binding protein 5 haplotype. Development and Psychopathology 24, 1237–1252 (2012). URL https://doi.org/10.1017/s0954579412000673.

[12] VanZomeren-Dohm, A. A., Pitula, C. E., Koss, K. J., Thomas, K. & Gunnar, M. R. FKBP5 moderation of depressive symptoms in peer victimized, post-institutionalized children. Psychoneuroendocrinology 51, 426–430 (2015). URL https://doi.org/10.1016/j.psyneuen.2014.10.003.

[13] Millstein, R. A. & Holmes, A. Effects of repeated maternal separation on anxiety- and depression-related phenotypes in different mouse strains. Neuroscience & Biobehavioral Reviews 31, 3–17 (2007). URL https://doi.org/10.1016/j.neubiorev.2006.05.003.

[14] Campos, A. C., Fogaca, M. V., Aguiar, D. C. & Guimaraes, F. S. Animal models of anxiety disorders and stress. Revista Brasileira de Psiquiatria 35, S101–S111 (2013). URL https://doi.org/10.1590/1516-4446-2013-1139.

[15] Floriou-Servou, A. et al. Distinct proteomic, transcriptomic, and epigenetic stress responses in dorsal and ventral hippocampus. Biological Psychiatry 84, 531–541 (2018). URL https://doi.org/10.1016/j.biopsych.2018.02.003.

[16] Goodwill, H. L. et al. Early life stress leads to sex differences in development of depressive-like outcomes in a mouse model. Neuropsychopharmacology 44, 711–720 (2018). URL https://doi.org/10.1038/s41386-018-0195-5.

[17] Davis, E. P. & Pfaff, D. Sexually dimorphic responses to early adversity: Implications for affective problems and autism spectrum disorder. Psychoneuroen-docrinology 49, 11–25 (2014). URL https://doi.org/10.1016/j.psyneuen.2014.06.014.

[18] Moisan, M.-P. Sexual dimorphism in glucocorticoid stress response. International Journal of Molecular Sciences 22, 3139 (2021). URL https://doi.org/10.3390/ijms22063139.

[19] Lee, R. S. et al. DNA methylation and sex-specific expression of FKBP5 as correlates of one-month bedtime cortisol levels in healthy individuals. Psy-choneuroendocrinology 97, 164–173 (2018). URL https://doi.org/10.1016/j.psyneuen.2018.07.003.

[20] Walker, J. J. et al. The origin of glucocorticoid hormone oscillations. PLoS Biology 10, e1001341 (2012). URL https://doi.org/10.1371/journal.pbio.1001341.

[21] McEwen, B. S. Mood disorders and allostatic load. Biological Psychiatry 54, 200–207 (2003). URL https://doi.org/10.1016/s0006-3223(03)00177-x.

[22] Reppert, S. M. & Weaver, D. R. Coordination of circadian timing in mammals. Nature 418, 935–941 (2002). URL https://doi.org/10.1038/nature00965.

[23] Wuäst, S. et al. The cortisol awakening response - normal values and confounds. Noise Health 2, 79–88 (2000).

[24] Fu, Y. et al. Intrinsically photosensitive retinal ganglion cells detect light with a vitamin a-based photopigment, melanopsin. Proceedings of the National Academy of Sciences 102, 10339–10344 (2005). URL https://doi.org/10.1073/pnas.0501866102.

[25] Aschoff, J., Gerecke, U. & Wever, R. DESYNCHRONIZATION OF HUMAN CIRCADIAN RHYTHMS. The Japanese Journal of Physiology 17, 450–457 (1967). URL https://doi.org/10.2170/jjphysiol.17.450.

[26] Fries, E., Dettenborn, L. & Kirschbaum, C. The cortisol awakening response (CAR): Facts and future directions. International Journal of Psychophysiology 72, 67–73 (2009). URL https://doi.org/10.1016/j.ijpsycho.2008.03.014.

[27] Vrshek-Schallhorn, S. et al. The cortisol awakening response predicts major depression: predictive stability over a 4-year follow-up and effect of depression history. Psychological Medicine 43, 483–493 (2012). URL https://doi.org/10.1017/s0033291712001213.

[28] Huber, T. J., Issa, K., Schik, G. & Wolf, O. T. The cortisol awakening response is blunted in psychotherapy inpatients suffering from depression. Psychoneuroendocrinology 31, 900–904 (2006). URL https://doi.org/10.1016/j.psyneuen.2006.03.005.

[29] Coppen, A. et al. Dexamethasone suppression test in depression and other psychiatric illness. British Journal of Psychiatry 142, 498–504 (1983). URL https://doi.org/10.1192/bjp.142.5.498.

[30] Stapelberg, N. et al. From feedback loop transitions to biomarkers in the psycho-immune-neuroendocrine network: Detecting the critical transition from health to major depression. Neuroscience & Biobehavioral Reviews 90, 1–15 (2018). URL https://doi.org/10.1016/j.neubiorev.2018.03.005.

[31] Koch, C., Leinweber, B., Drengberg, B., Blaum, C. & Oster, H. Interaction between circadian rhythms and stress. Neurobiology of Stress 6, 57–67 (2017). URL https://doi.org/10.1016/j.ynstr.2016.09.001.

[32] Nold, V. et al. Activation of the kynurenine pathway and mitochondrial respiration to face allostatic load in a double-hit model of stress. Psychoneuroendocrinology 107, 148–159 (2019). URL https://doi.org/10.1016/j.psyneuen.2019.04.006.

[33] Conrad, C. D. Chronic stress-induced hippocampal vulnerability: The glucocorticoid vulnerability hypothesis. Reviews in the Neurosciences 19 (2008). URL https://doi.org/10.1515/revneuro.2008.19.6.395.

[34] McEwen, B. S., Gould, E. A. & Sakai, R. R. The vulnerability of the hippocampus to protective and destructive effects of glucocorticoids in relation to stress. British Journal of Psychiatry 160, 18–23 (1992). URL https://doi.org/10.1192/s0007125000296645.

[35] Harrell, C., Gillespie, C. & Neigh, G. Energetic stress: The reciprocal relationship between energy availability and the stress response. Physiology & Behavior 166, 43–55 (2016). URL https://doi.org/10.1016/j.physbeh.2015.10.009.

[36] Sheline, Y. I., Sanghavi, M., Mintun, M. A. & Gado, M. H. Depression duration but not age predicts hippocampal volume loss in medically healthy women with recurrent major depression. The Journal of Neuroscience 19, 5034–5043 (1999). URL https://doi.org/10.1523/jneurosci.19-12-05034.1999.

[37] Small, T. W. et al. Stress-responsiveness influences baseline glucocorticoid levels: Revisiting the under 3 min sampling rule. General and Comparative En-docrinology 247, 152–165 (2017). URL https://doi.org/10.1016/j.ygcen.2017.01.028.

[38] Chou, B.-K. et al. A facile method to establish human induced pluripotent stem cells from adult blood cells under feeder-free and xeno-free culture conditions: A clinically compliant approach. STEM CELLS Translational Medicine 4, 320–332 (2015). URL https://doi.org/10.5966/sctm.2014-0214.

[39] Shi, Y., Kirwan, P. & Livesey, F. J. Directed differentiation of human pluripotent stem cells to cerebral cortex neurons and neural networks. Nature Protocols 7, 1836–1846 (2012). URL https://doi.org/10.1038/nprot.2012.116.

[40] Yuan, S. H. et al. Cell-surface marker signatures for the isolation of neural stem cells, glia and neurons derived from human pluripotent stem cells. PLoS ONE 6, e17540 (2011). URL https://doi.org/10.1371/journal.pone.0017540.

[41] Steiger, J. H. Beyond the f test: Effect size confidence intervals and tests of close fit in the analysis of variance and contrast analysis. Psychological Methods 9, 164–182 (2004). URL https://doi.org/10.1037/1082-989x.9.2.164.

[42] Kelley, T. L. An unbiased correlation ratio measure. Proceedings of the National Academy of Sciences 21, 554–559 (1935). URL https://doi.org/10.1073/pnas.21.9.554.

[43] Olejnik, S. & Algina, J. Generalized eta and omega squared statistics: Measures of effect size for some common research designs. Psychological Methods 8, 434–447 (2003). URL https://doi.org/10.1037/1082-989x.8.4.434.

[44] Allen, R. Statistics and Experimental Design for Psychologists (WORLD SCIENTIFIC (EUROPE), 2017). URL https://doi.org/10.1142/q0019.

